# Massively multiplexed microfluidics maps combinatorial and sequential antibiotic responses in 3D

**DOI:** 10.1101/2025.10.30.684665

**Authors:** Yoon Jeong, Gabriel Mercado-Vasquez, Abinash Padhi, Yiyu Deng, Madhumitha Prakash, Jonathan M. Matthews, Sefika Ozcan, Erdal Toprak, Savaş Tay

## Abstract

Current microbial culture and antibiotic susceptibility testing platforms lack dynamic chemical control and do not scale to high-throughput 3D culture. We present an automated microfluidic system that creates 512 independently programmable 3D hydrogel-culture chambers and delivers each chamber a distinct combinatorial and time-varying drug protocol. Automated workflows execute thousands of micro-pipetting operations and track 10,000 bacterial colonies via live-cell microscopy, and generate half a million single colony images to quantify growth, drug response and morphology. Using this system, we study how hydrogel stiffness and nutrient timing alter colony architecture and modulate antibiotic susceptibility and resistance. We then profile the efficacy of antibiotic pairs using simultaneous and sequential (temporally ordered) dosing across 2,700 drug–dose combinations, revealing synergy, antagonism and order dependent shifts in drug efficacy. This platform integrates serial and parallel multiplexing, automation, and dynamic 3D microenvironments to map the chemical and mechanical determinants of antibiotic response and drug interactions.

## Main Text

The chemical composition of cellular microenvironments changes across a wide range of spatial and temporal scales^1^, exposing cells to time-varying combinations of nutrients, signaling molecules and drugs. Cells adapt to these conditions and organize into complex three-dimensional (3D) structures that support their function and growth^2^. Reconstituting such dynamic microenvironments *in vitro* enables mechanistic studies of drug response and cell-cell interactions. Lessons learned from studying cells under precisely controlled dynamic conditions can inform therapeutic strategies. For example, sequential antibiotic administration where different drugs are delivered according to a specific order and timing holds promise for managing challenging bacterial infections^3,4^. Drug timing is critical in sequential therapies, and the order and duration of drug exposure can influence synergy or antagonism between individual drugs, or can lead to resistance^5,6^. Overlapping (simultaneous) or sequential (temporally ordered) drug delivery can alter microbial responses and therapeutic outcomes idiosyncratically in clinical settings^7^. Studying such scenarios enables designing better therapies and understanding the mechanisms behind antibiotic susceptibility, resistance, and other antibiotic survival mechanisms such as persistence and tolerance.

Live-cell experiments are ideal for systematic testing of drug dosage, timing and duration, and monitoring phenotypic shifts due to cell density or nutrient availability. Longitudinal imaging of live cells and colonies allow real-time observation of functional outcomes like adaptation, resistance, or phenotypic variability^8^. Unfortunately, current live-cell analysis methods suffer from limitations in sensitivity, throughput, and flexibility. Conventional techniques such as microdilution lack the capability to dynamically image cells and adjust the administration of antibiotics, and are not suitable to study drug order, timing and concentration in high throughput^9^. The inability to fine-tune drug delivery parameters like timing and combinations prevents researchers from fully exploring and optimizing sequential combination regimens^10^. Robotic systems are unable to combine 3D cell culture and dynamic drug exchanges with real-time tracking of cells. Besides, manual or robotic pipetting-based methods cannot readily mimic clinically relevant drug changes without altering the cell density, which itself influences growth and drug susceptibility. Overall, technical limitations in capturing the dynamic physical and chemical context of bacterial microenvironments directly impact the predictive value of antibiotic susceptibility testing (AST) and hinder the development of effective antimicrobial combinations.

Microfluidics offers unique capabilities in creating and precisely controlling physiologically relevant cellular microenvironments. Microfluidics enables complex culture geometries and dynamic control of the chemical environment at high spatial and temporal resolution^11,12^. Recent advances in microfluidic automation^13^, programmable fluid handling^14^, machine learning-driven analysis^15^, and 3D cell culture^8,16^ significantly impacted microbial and other live-cell applications. However, scaling microfluidic live-cell analysis systems remains a challenge^17,18^, particularly when dynamic fluidic control and 3D culture need to be combined^19,20^. Previous platforms^21,22,23^ such as microfluidic chemostat^24,25^ rely on liquid cultures of bacteria and do not replicate the complex heterogeneous 3D conditions found in bacterial ecosystems. 3D cell culture often requires embedding cells in natural or engineered gels, which physically prevents rapid and dynamic fluidic delivery to cells. As a result, no technological platform thus far integrated high-throughput, dynamic, and real-time profiling of cellular responses through 3D microbial cultivation^26,27^.

Here, we introduce a microfluidic culture and live-cell analysis system with hundreds of independently programmable 3D culture chambers for bacterial cultivation and time-lapse analysis under dynamic drug perturbations. Our system represents a massive increase in multiplexing capability by generating a wide-range of chemical and fluidic perturbations across different types of molecules, doses, combinations, and temporal modulations. Technical advances in design, large-scale integration, automation, and active control of soft materials culminated in a capable system for high-throughput and high-content study of microbes and microbial interactions.

First, we developed a new method to build inner-compartmentalized microfluidic chambers incorporating hydrogel islands that trap and cultivate bacterial cells in 3D, while still retaining dynamic fluid delivery to the cultured cells. The combination of these capabilities is enabled by precise control of hydrogel formation inside microfluidic flow chambers—with high spatial and temporal resolution—allowing culture in 3D and immobilization of cells for time-lapse imaging and tracking. Gels with varying stiffness, chemical compositions and geometries can be loaded into each chamber, and different cell types can be trapped and cultured in them. The use of hydrogel for culture enables immobilization of non-adherent cells, viable formation and imaging of 3D colonies, and modulating the matrix stiffness supports diverse morphological features.

Furthermore, an integrated input-generation architecture automatically produces thousands of drug or media combinations and temporal sequences, enabling a massive chemical “input space” to be screened, and delivers unique formulations to 512 independent culture chambers at predetermined times. Our system automatically generates an extremely wide range of drug combinations, dose ranges, and temporal schedules (pulses, increasing-decreasing concentrations, or sequences of drugs). Dense microfluidic multiplexing ensures that each culture chamber receives a different drug program without influencing the others, and these programs can be changed over time when desired. To achieve these functions, our system integrates 1,400 automated switchable PDMS valves driving ∼6×10^5^ valve actuations per experiment to perform 4,000 micro-pipetting–equivalent fluidic procedures such as gel formation, cell seeding, individualized media supply, washes, dilutions, and combinations. Automation of image acquisition results in generation of nearly half a million images of individual colonies per experiment. We also developed custom bacterial colony segmentation and tracking algorithm for automated analysis of colony growth and morphology. These functionalities are combined in an integrated system, achieving unprecedented multiplexing, throughput and functionality for 3D microbial cell culture applications.

Using our system, we studied the influence of matrix stiffness, nutrient timing, drug combinations, and drug temporal sequences on bacterial growth, antibiotic susceptibility, resistance and morphology. We first investigated how hydrogel matrix stiffness (10–300 kPa) influences morphology and antibiotic susceptibility alteration in *E. coli* (*Escherichia coli* Nissle 1917). We then performed antibiotic susceptibility testing (AST) and evaluated resistance across four bacterial species (*E. coli, S. aureus, P. aeruginosa, K. pneumoniae*). Thirty-six pairwise formulations from nine clinically important antibiotic compounds (Amikacin, Cefepime, Clindamycin, Levofloxacin, Nitrofurantoin, Sulfamethoxazole, Rifampin, Tetracycline, and Trimethoprim) were delivered into each of the 512 chambers in simultaneous (i.e. A and B together; denoted A+B) and sequential manner (i.e. A then B; denoted A→B). Individual colonies were tracked over time to measure colony growth, and antibiotic susceptibility was analyzed in each condition.

Analysis of 2,700 distinct drug conditions revealed several order-dependent and order-independent synergistic and antagonistic interactions among commonly used drugs. Most importantly, we mapped the dynamic temporal landscape of antibiotic–antibiotic interactions in live bacterial cells and colonies. We found several dynamic interactions— synergistic, neutral, or antagonistic—between antibiotic pairs when delivered in different orders. These findings have the potential to improve antibiotic susceptibility testing, while minimizing antibiotic-related toxicities and recurrent infections that drive the evolution of resistance. Overall, these studies demonstrate the unique capabilities of our microfluidic live-cell analysis system for massively multiplexed studies of bacteria-drug interactions under time-dependent perturbations.

## Results

### Building hydrogel-based 3D culture compartments within microfluidic chambers

Hydrogels are widely used across microbial and mammalian cell culture applications because they recapitulate essential features of the native extracellular matrix (ECM) or extracellular polymeric substances (EPS), providing structural and biochemical cues that regulate cellular behavior^28^. Engineered gels offer tunable mechanical and chemical properties—such as stiffness, viscoelasticity, degradability, and ligand density—and can be functionalized to present adhesion peptides, growth factors, and signaling molecules^20^. These features support physiologically relevant growth conditions and cell–cell communication. The porosity of hydrogels facilitates diffusion of gases, nutrients, metabolites, and drugs, while the 3D matrix protects embedded cells from shear forces^29^. Importantly, embedding otherwise non-adherent cells in hydrogels protects them from displacement due to flow during drug delivery and chemical stimulation. This enables long term imaging and tracking of individual colony growth and morphological changes. It also prevents cell density changes during media or drug delivery, which is critical for evaluating drug efficacy and treatment success^27^.

While gel-supported culture is the favorable mode of cultivation for many cell types, repeated delivery and removal of reagents, drugs and liquid media to 3D hydrogel cultures is a difficult task^19^, especially at the microscale. Loading gels to microfluidic systems leads to clogging of flow channels^20^, and it is difficult to precisely control where and when gelation processes initiate. To overcome these limitations, we developed a process for controlled formation of hydrogel-supported 3D compartments within microfluidic chambers (Fig 1a, Extended Data Fig 1a). To control the location and timing of gel formation, we incorporate a ring-shaped PDMS membrane valve (i.e., encapsulation valve) in the microfluidic culture chamber that can be automatically opened and closed with external pressure (Δ*p* = 35 psi). Closing of the valve allows physical isolation of a central area defined by valve geometry within the chamber (Extended Data Fig 1b,c). A liquid hydrogel precursor can be loaded into this isolated region, which is then exposed to crosslinkers to form a hydrogel-island. To facilitate the gelation processes, three pneumatic valves (inlet, outlet, and encapsulation valves) act as a functional unit (Fig 1d), where a hydrogel gel precursor can be crosslinked or de-crosslinked through sequential actuation steps (Supplementary Note 1).

**Fig 1.**
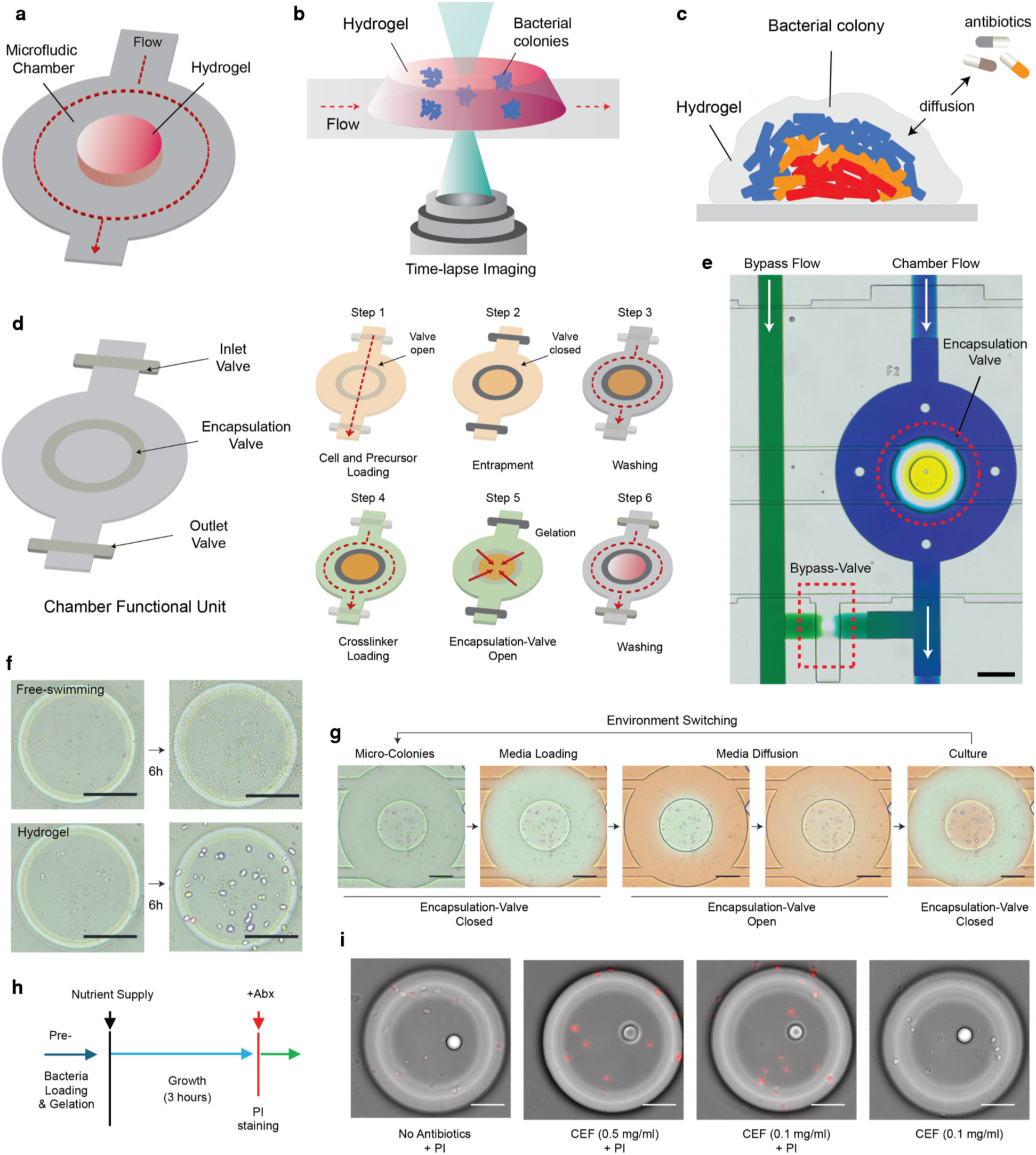
Microfluidic chambers for hydrogel-based cell encapsulation, 3D culture, and dynamic chemical stimulation. **a**. Schematic of a microfluidic culture chamber with hydrogel island showing flow direction (red dashed arrow) and chamber structure. Cells are encapsulated and cultured in the hydrogel island. The surrounding microfluidic structure is used for controlled and time-dependent delivery and removal of reagents and media. **b**. Time-lapse imaging setup for real-time monitoring of bacterial colonies in hydrogel, with flow direction indicated. **c**. Conceptual diagram of the spatial arrangement of cells in hydrogel, highlighting subpopulation heterogeneity, antibiotic exposure via diffusion, and bacteria shielded by polysaccharide/eDNA hydrogel barrier. **d**. Functional unit of the microfluidic chamber with inlet, outlet, and encapsulation valves for flow control and cell isolation. Sequential steps for hydrogel island construction: precursor (0.5-2% w/v alginate solution) with inoculum loading, valve closure for gel entrapment, encapsulation-valve open for gelation via exposure to a crosslinker, and washing of un-crosslinked precursor (see Methods, Supplementary Note 1, and Supplementary Videos 1 and 2). **e**. Microscopic image of an actual unit chamber, showing bypass flow line (green), chamber flow (blue), valve positioning, and inner-chamber compartmentalization (yellow); scale bar, 200 μm. **f.** Time-series images comparing free-swimming (planktonic) and hydrogel-embedded *E. coli* over 6 hours, highlighting colonized growth pattern; scale bar, 100 μm. **g**. Chemical environmental switching illustrating liquid media loading, reagent diffusion into hydrogel island, and culture stages with encapsulation-valve status (open/closed). Inner circle shows hydrogel with *E. coli* colonies; scale bar, 100 μm. **h**. Experimental procedure for nutrient supply, antibiotic administration, 3-hour growth, and PI staining, with timeline of bacterial loading and gelation. **i**. Microscopic images of individual *E. coli* colonies in the hydrogel island under scheduled exposures to various drug conditions (no antibiotics + PI, CEF 0.5 mg/ml + PI, CEF 1 mg/ml + PI, CEF 1 mg/ml + no PI), merging bright-field and rfp channels; scale bar, 50 μm.

To start the gel-forming process, first a high viscous gel precursor (viscosity, 50-200 *cP*) is flown into the microfluidic chamber by opening all the valves. Then, the encapsulation valve is closed, isolating a central region and the un-crosslinked precursor. The precursor solution from the surrounding area is then washed away. In the final step, a crosslinking reagent is flowed in, and the encapsulation valve is opened. The contact between the crosslinking reagent and the precursor leads to near-instantaneous formation of a gel island at the center of the microfluidic chamber. Microbial cells can be inoculated in this region by initially mixing them with the precursor during the first step.

This automated procedure results in the formation of a gel island containing bacterial cells, directly placed in the middle of a microfluidic chamber (see Supplementary Video 1). The empty space around the island is used for flow and rapid fluidic exchanges, leading to reagent delivery to cells via diffusion. The chamber retains inlet and outlet valves, allowing controlled delivery of fluids and reagents to the gel island. Rapid molecular transport via diffusion through the porous gel enables effective exchange of nutrients, waste, and reagents between the cells in gel, and their environment (Supplementary note 2 and Supplementary Fig 3). Furthermore, the 3D support provided by the gel enables formation of cultures and colony structures that mimic micro-physiological 3D environments. The encapsulation valve can be used for further operations such as decrosslinking of the gel, removal of cells for downstream analysis off the chip, or for complete physical isolation of cells from their environment. Multiple hydrogel islands can be placed inside the same microfluidic chamber and can be individually controlled, creating complex culture geometries such as co-culture islands in the same microfluidic environment (Supplementary Fig 4).

We screened nine natural polysaccharide polymers (e.g., alginate, gelatin, agarose) to evaluate their properties, including viscosity (*cP*), gelation process, timing, reaction reversibility, pH, and their adaptability to microfluidic experiments and live cell culture (Supplementary Table 1). Among these, alginate hydrogel showed the highest compatibility for our inner-compartmentalized microfluidic design. Its reversible crosslinking properties^30^ effectively prevented hydrogel channel clogging and ensured a controlled crosslinked gelation processes with the valve actuation process in chambers (Extended Data Fig 1d). The integration of a bypass flow line enabled rapid washing and media transitions within each chamber (< 5-6 seconds) (Fig 1e). Further, we assessed the structural integrity of the hydrogel environment based on alginate stiffness factors^30^ for 3D culture environments (Supplementary Table 2).

Overall, our hybrid microfluidic-hydrogel environment sustains microbial population growth, facilitates molecular diffusion, and maintains structural integrity under time-dependent chemical perturbations, while enabling undisturbed 3D microbial culture and imaging (Fig 1b). The 3D spatial arrangement of bacterial colonies in hydrogels closely replicates natural microbial ecosystems, influencing antibiotic exposure and diffusion while enabling dynamic colony changes (Fig. 1c), thus offering a physiologically relevant environment suitable to study bacterial physiology and resistance mechanisms.

### Colony formation and dynamic reagent delivery to cells cultured in 3D gel islands

Our hybrid gel-microfluidic culture chambers solve the challenging problem of rapidly delivering soluble reagents to cells cultured in a gel environment. *E. coli* cells entrapped in hydrogels are enclosed within the chamber by closing of the encapsulating valve, which isolates them from the surrounding environment. Fresh culture media is then supplied to support their colonization and growth. Various molecules within the fluidic and hydrogel environment diffuse freely into the gel to enable molecular delivery to colonies, and removal of culture waste^31^. This *E. coli* strain showed distinct colonization patterns before gelation and after gelation in a 3D matrix (Fig 1f and Supplementary Video 2). Before gelation, the uncontrolled proliferation of free planktonic bacteria filled the chamber space. At this state, active flow cannot be used for media exchange or delivery of nutrients^26^ because the cells would be diluted or removed altogether. After gelation, bacterial inoculum were immobilized in the hydrogel island where a single microbe can form a colony, and fresh media can be delivered via flow, repeatedly. This enabled viable cell culture, colony formation and time-lapse tracking of colonies under growth conditions.

A key feature of our hybrid culture chamber is its capacity to perform repeated environmental switching, enabling multiple media or reagent delivery cycles at specific time points. (Fig 1g, Extended Data Fig 1e,f). To demonstrate dynamic drug administration to cells entrapped in the gel island, *E. coli* colonies were treated with cefepime (CEF, 0.1–0.5 mg/ml) and propidium iodide (PI, 5 µg/ml) after 3 hours of incubation (Fig 1h,i, Extended Data Fig 1g). Dead cells in the colonies were visualized by PI within 3 minutes of CEF administration while the hydrogel environment supported undisturbed imaging. Even after multiple reagent administrations, the hydrogel structure withstood the mechanical stress from repeated valve actuation, and cells remained viable (Supplementary Video 1). This functional design facilitates precise regulation of flow trafficking, retention, and timely delivery, supporting robust phenotypical characterizations in hydrogels.

### Automated colony tracking and quantification

We monitored microbial culture chambers via automated time-lapse microscopy (Fig 2, and Supplementary Video 3). For automated colony tracking, we developed image-processing algorithms through two tracks (Extended Data Fig 2a): 3D reconstructed visualization of microbial colonies and 2D contouring analysis for quantification. To quantify colony dynamics, a post-processing algorithm incorporates morphological decision criteria, such as circularity and convexity of contours, and enables setting different detection thresholds for segmenting colonies (Extended Data Fig 2b, Supplementary Note 3). This resulted in quantifying individual colony growth and tracking of spatial distribution patterns under different population densities.

**Fig 2.**
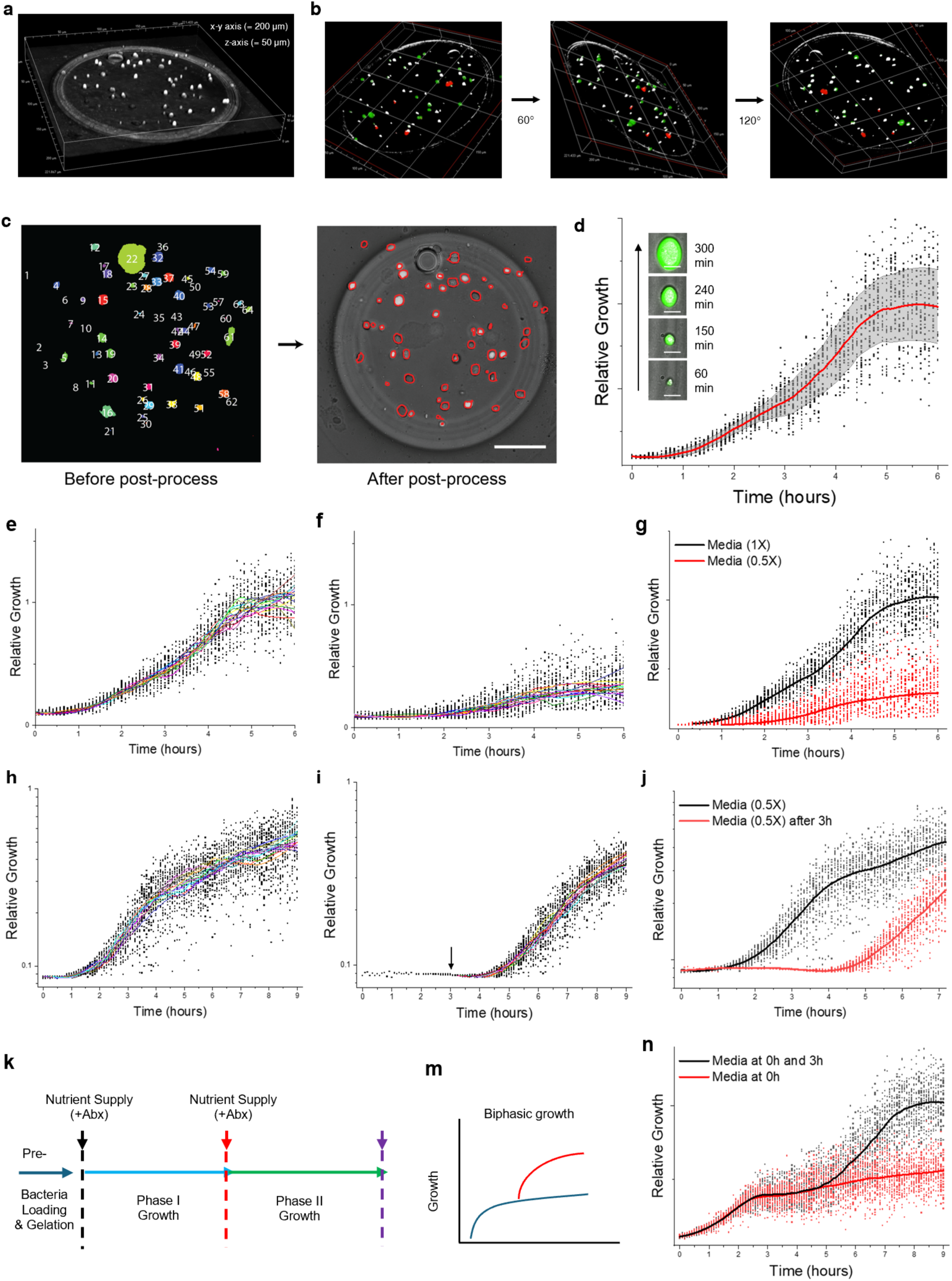
Automated time-lapse microscopy and image processing pipeline enables high-throughput colony tracking and quantification of colony growth. **a.** 3D volumetric visualization of *E. coli* colonies cultured in the hydrogel islands (200 μm × 200 μm x-y, 50 μm z-depth, bright-field). **b.** 3D visualization of colony distribution via 3D rendering (rotations at 60° and 120°) in a hydrogel culture chamber. Wild-type, gfp-expressing, and rfp-expressing E. coli Dh5α colonies are shown by merging bright-field, gfp, and rfp channels. **c.** 2D image analysis before and after post-processing, with pseudo-colored labels (left) and refined contoured colonies (right) for enhanced segmentation accuracy. Scale bar, 50 μm. Also see Supplementary Video 3. **d.** Real-time colony growth tracking over 6 hours from a single microfluidic chamber, showing relative growth of multiple colonies normalized by area measurement (see Supplementary Note 3). Measurement was taken every 5 minutes. Red line indicates average colony size over time, each dot represents a single colony. Mean ± s.d. (n = 30 colonies from biological replicates), with mean colony size (red line), scatter points (individual measurements), and black-shaded region (s.d.); inset images of gfp-expressing E. coli at 60, 150, 240, and 300 minutes. Scale bar, 5 μm. **e-g.** Growth tracking analysis over 6 hours for LB media concentrations: (e) 1X and (f) 0.5X. Solid lines show mean colony size per chamber (n = 15 chambers), dots show individual colonies (n = 30, subsampled). **h-j.** Growth tracking analysis comparing (h) initial LB 0.5X media to (i) fresh 0.5X media supplied at 3 hours. Solid lines show mean colony size per chamber (n = 15 chambers), dots show individual colonies (n = 30, subsampled). (e-j) For presentation purposes, individual colony data were randomly subsampled from raw data. **k.** Experimental setup for two growth phases: Phase I followed by Phase II (re-growth). **m.** Depiction of biphasic growth pattern. **n.** Colony growth tracking plot over 9 hours with initial LB 0.5X media at 0h and fresh 0.5X media at 3h; Mean colony size (red lines, n = 30 colonies per condition, biological replicates).

Through our custom image-processing pipeline, we reconstructed volumetric images of colony distribution cultured in a single hydrogel island (Fig 2a,b, Extended Data Fig 2c,d). In addition, the pipeline facilitated high-throughput phenotyping such as colony count and size, morphology, spatial distribution and quantitative analysis of growth patterns of bacterial colonies after post-processing (Fig 2c, Extended Data Fig 2e). Real-time colony tracking analysis of *E coli* generated growth curves (n = 30 colonies) over a 6-hour period to capture population-level growth trends, which were visualized as average relative growth trajectories (Fig 2d). In Figure 2d, each dot represents the relative size of a single bacterial colony, and solid lines indicate the average colony size over time for each independent culture chamber in our integrated device. Our pipeline generates a high number of single colony measurements per experiment, and many panels in Figure 2 (specifically e, f, g, i and j) had to be randomly sub-sampled for presentation purposes.

### Nutrient timing modulates colony growth in microfluidic hydrogel islands

Nutrient availability and timing are critical factors that significantly influence the progression and characteristics of growth phases in bacterial cultures. Automated and dynamic reagent delivery, precisely timed by our platform, enables control over nutrient availability in the microfluidic environment. We analyzed the impact of varying nutrient levels (0.5–1X LB media, Fig 2e-g) and the specific timing of nutrient supply at 0 and 3 hours (0.5X LB media, Fig 2h-j) in shaping the observed growth phases. *E. coli* population growth reached a plateau after 3-4 hours of incubation during nutrient baseline monitoring.

When nutrients (0.5X LB media) were delivered at 3 hours, a secondary growth phase began (Fig 2k, m). This response is noteworthy because it is similar to in vivo observations from living organisms^32^. The additional nutrient supply at the 3-hour mark induced the secondary phase of proliferation, which shows distinct biphasic growth patterns (Fig 2n). This biphasic response underscores the dynamic adaptability of the cells to changes in nutrient availability and highlights the importance of nutrient timing in optimizing bacterial growth.

### Large-scale device integration, dense multiplexing and automation

To improve throughput, multiplexing capabilities, and testing of a wide range of reagent combinations on live cultures, we used large-scale microfluidic integration methods and built a single device that incorporates 512 independently programmable hydrogel culture units. For integration and fluidic control, we designed key components including an input generation module for automated mixing and preparation of reagent combinations and dilutions (Fig 3a). We also developed a large-scale parallelized chamber array and integrated them into the same device (Extended Data Fig 3a). Individualized fluidic control in each chamber is critical for high throughput experiments in parallel units^18^. Row and column multiplexers were implemented for maximizing the parallelized device’s throughput and individualized control of culture chambers (Fig 3b, Extended Data Fig 3b, Supplementary Videos 4 and 5).

**Fig 3.**
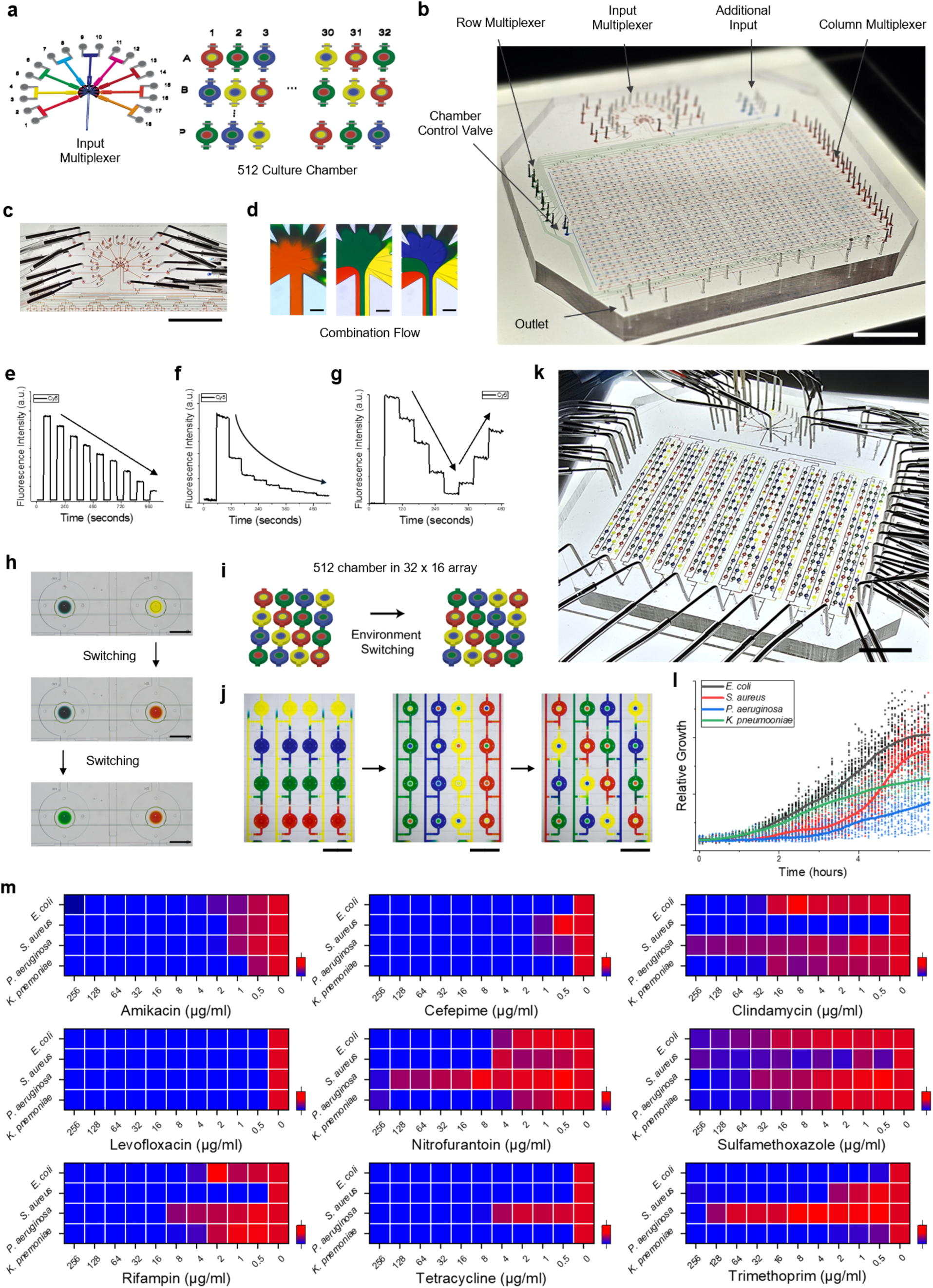
Integrated device with parallel hydrogel colony cultures, serial chemical formulation, and dynamic delivery of drug regiments to each unit. Different colored food dyes were used for visualization of device geometry and fluid flow. a. Schematic of chemical input generator (Supplementary Video 4) and 512 independent culture chambers arranged in a 32 × 16 array (Supplementary Video 5). b. Image of the integrated microfluidic device with multiplexed modules (row/column multiplexers, input, chamber control valve, see Supplementary Note 4). Scale bar, 1.5 cm. c. Input multiplexer image with radially arranged fluidic inputs; scale bar, 1.5 cm. d. Images of combination fluid states with 1, 3, and 4 reagent mixtures; scale bar, 150 μm. Different colored food dyes were used for visualization. e-g. n-fold dilution profiles over time: (e) linear (9:0, 8:1, 7:2 to 1:8) and (f) logarithmic (1:0, 1:1, 1:2 to 1:8), with cy5 fluorescence intensity control showing multiplexer precision. h. Encapsulation-chamber color switching demonstration; scale bar, 500 μm. The fluids inside the encapsulated area can be quickly changed by using the valve. i. Checkerboard-like pattern switching for each chamber, demonstrating individual addressability and flow of each culture chamber. Each chamber can be loaded with a different fluidic formulation independent of the others. j. Images of fluid switching in a 4 × 4 chamber array; scale bar, 1.5 mm. k. Whole integrated chip configuration with 512 chambers filled with red, yellow, blue, and green dyes, indicating multiplexing controllability and independent chamber loading; scale bar, 1.5 cm. l. Relative growth tracking curves for four bacterial species (*E. coli, S. aureus, P. aeruginosa, K. pneumoniae*) in MH2 media over 6 hours in the integrated device; scatter plots with mean curves (n ≥ 30 colonies per condition, biological replicates). m. Antibiotic susceptibility testing heatmaps of four bacterial species, each showing susceptibility to nine antibiotics (AMI, CEF, CLI, LEV, NIT, SUL, RIF, TET, TRI, see Supplementary Table 4), across a wide range of concentrations (from 256 to 0.5 µg/ml, 2-fold dilution); color gradient from blue (high susceptibility, no growth) to red (resistance, maximal growth) after 6 hours. Overall, 396 different antibiotic conditions have been tested in a single integrated device, with tens of thousands of individual colony measurements over 6 hours.

Accurate generation of reagent combinations with varied dilutions has proved to be difficult in high throughput testing across parallel conditions^8^. To ensure symmetrical and reproducible combinatorial fluid mixing, we radially arranged reagent input components around a central outlet with nine nodes (Fig 3c, Extended Data Fig 3c,d). Multiple fluidic streams converged into the outlet as they flowed through a merging channel design, leading to a more homogenized mixture (Fig 3d, Extended Data Fig 3e,f and Supplementary Video 4). We then evaluated the input multiplexer’s performance by generating n-fold concentration gradients for targeted concentrations^33^(Fig 3e-g). By adjusting mixing ratios with a buffer solution, the input multiplexer achieved both linear and logarithmic concentration profiles with high precision (<1% variability for targeted concentrations). Our approach also eliminated delays in reagent delivery typically seen in separated microfluidic module systems^16,34^, where a time delay of 30–60 seconds is common^35^. Our system reduced reagent fluid travel time through channels to 1-2 seconds across 512 chambers, significantly minimizing experimental delays. Overall, our fluidic multiplexing module allowed efficient, rapid and accurate generation of drug combinations and dilutions across a wide range.

The functional integrity and fluidic transitions across 512 chambers were assessed by visualization of flow patterns and food-dye color transitions between chambers (Fig 3h, Extended Data Fig 3g,h, and Supplementary Video 5). We inspected our system to verify the absence of fluid leakage or any functional defects for subsequential reagents delivery and switching, both within and outside the chamber area (Fig 3i-k, Extended Data Fig 3i,j). To dynamically alter all parallel conditions in the device, we automated control of 1,420 monolithic valves, enabling valve actuation with a latency < 100 ms (Extended Data Fig 4a). Culture chamber and multiplexing modules were controlled through a fully automated workflow^36^ where our custom code and (GUI) executes over 4000 fluidic sequence steps per experiment (Extended Data Fig 4b). Through this automated framework, we calibrated the whole system, addressing hardware design, fluid dynamics, culture control systems, and data acquisition with full-automation (Extended Data Fig 4c, Supplementary Note 4). Overall, our integrated design results in a massively multiplexed experimental pipeline, with parallel and serial multiplexing of a wide range of experimental conditions (drug formulations, delivery sequences, timings, and imaging steps), loading these conditions to 512 independent chambers repeatedly (Supplementary Video 5), and quantitative testing of a high number of independent experimental sequences.

### High-throughput antibiotic susceptibility testing

Prior to antibiotics testing in the device, we evaluated the growth dynamics of clinically significant bacterial species in standard culture media: *Streptococcus pneumoniae* (ATCC 49619), *Pseudomonas aeruginosa* (ATCC 10145), and *Klebsiella pneumoniae* (ATCC 13883). The relative growth of these species over a 6-hour period revealed distinct differences and established the baseline growth metrics with the absence of drugs (Fig 3l). The slowest growth phase was observed for *P. aeruginosa,* and *S. aureus* that exhibited delayed growth around 2 hours of incubation (Extended Data Fig 5a–e).

We then tested bacterial growth in the presence of nine antibiotics (abbreviated AMI, CEF, CLI, LEV, NIT, SUL, RIF, TET, and TRI; classified by family class and mode of action, Supplementary Table 3) in our 512-chamber array to measure susceptibility against CLSI (Clinical & Laboratory Standards Institute) ranges^37^ (0.5–256 µg/ml, tested by 2-fold dilution) (Fig 3m). Overall, 396 distinct antibiotics and dilutions have been tested in a single device, and growth data has been collected over a 6-hour period. *P. aeruginosa* exhibited greater resistance to the antibiotics compared to the other test species, a finding attributable to its inherent antibiotic resistance^38^(Extended Data Fig 5f-j). Interspecies (between microbial species) variability in growth and susceptibility is clearly seen, which underscores species-specific responses to different antibiotics. Nevertheless, all the minimal inhibitory concentration (MIC) results obtained from the platform were consistent with the known CLSI susceptible ranges established by other measurement methods (Supplementary Table 4). Taken together, our system effectively evaluates a wide range of species-specific AST responses in 3D hydrogel environments.

### Evaluation of matrix-stiffness driven antibiotic resistance

Soft hydrogel mimics an EPS-like matrix, known for protecting microbial colonies^39^ from environmental insults by forming a physical barrier (Fig 4a). The spatial arrangement of microbial subpopulations (i.e., biofilm heterogeneity^10^) affects antibiotic exposure. The hydrogel matrix may play varied functional roles in microbial physiology. We examined various gelation conditions for colonization of *E. coli* by controlling stiffness factors: alginate concentration (0.5 – 2% w/v) and CaCl₂ concentration (0.1–100 mM) (see Supplementary Table 2, Extended Data Fig 6a). *E. coli* growth exhibited a sharp contrast under varying hydrogel stiffness (10–300 kPa) within the culture chambers (Fig 4b, Extended Data Fig 6b). Specifically, the liquid (sol) state (alginic acid 1% w/v only) led to more dispersed *E. coli* growth, while the gel state crosslinked by 0.08 M CaCl₂ solution resulted in more condensed *E. coli* growth (Fig 4c). To ensure experimental consistency, we narrowed the range of gelation conditions by adjusting the alginate hydrogel stiffness to the 150–300 kPa range (see Supplementary Note 3).

**Figure 4.**
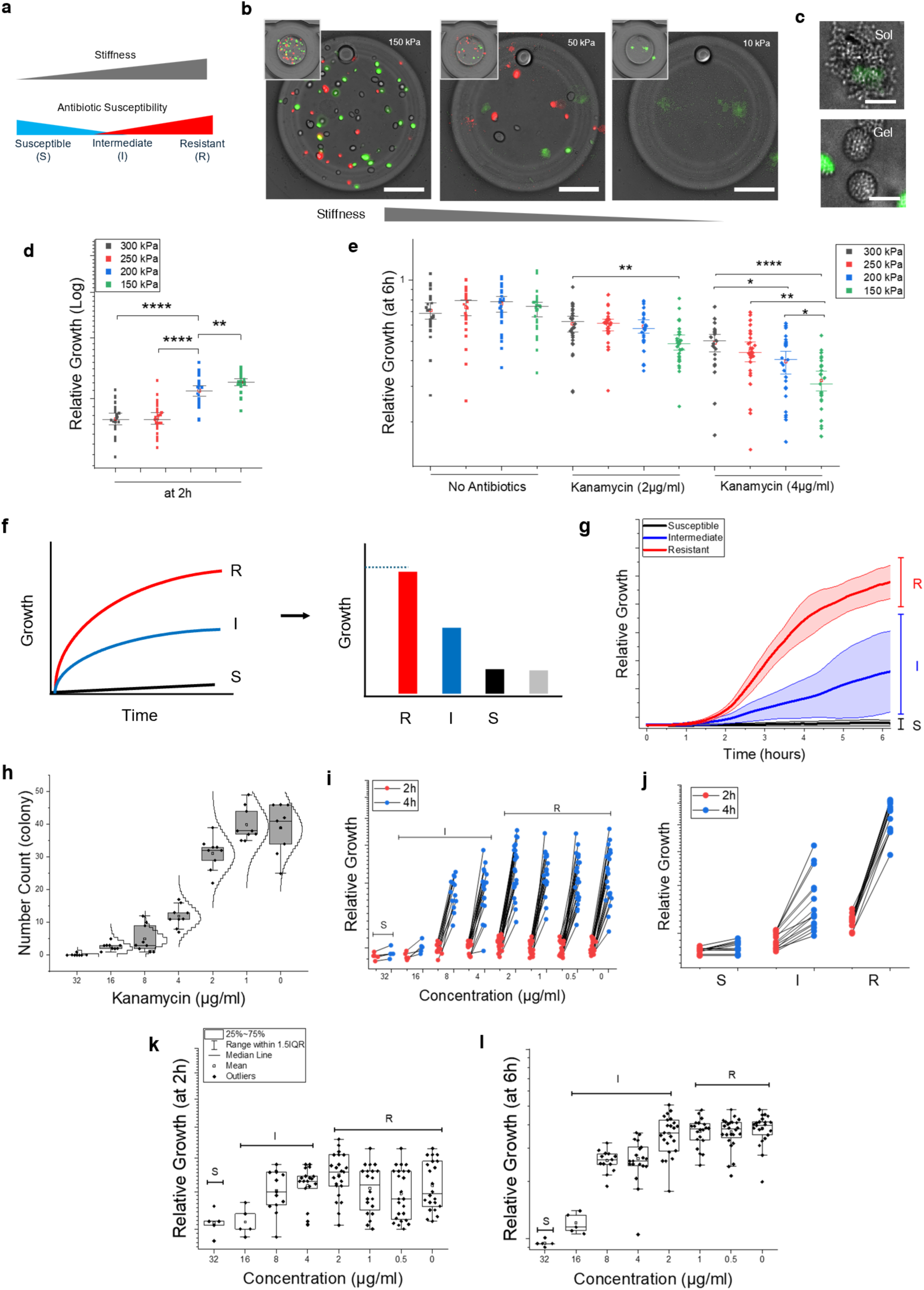
Hydrogel stiffness influences bacterial colony growth and antibiotic susceptibility categorization. **a.** Hydrogel stiffness can influence categorization of microbes due to their antibiotic susceptibility (SIR: Susceptible [S], Intermediate [I], Resistant [R]). **b.** Fluorescence microcopy of three hydrogel islands with varied stiffness (150, 50, 10 kPa), showing morphological patterns in wild-type, gfp-expressing, and rfp-expressing *E. coli* Dh5α colonies. Microscopy channels in bright-field, gfp, and rfp channels have been merged for analysis. Scale bar, 50 μm. **c.** Comparative images of sol (liquid, 0.5% w/v alginate solution) and gel (solid, crosslinked by 0.08 M CaCl₂) states for wild-type and gfp-expressing *E. coli*, analyzed by merging bright-field and gfp channels. Scale bar, 10 μm. **d.** Relative *E. coli* growth at 2 hours under stiffness conditions 150, 200, 250, 300 kPa. **e.** Relative *E. coli* growth with kanamycin (2 and 4 µg/ml) at 6 hours under stiffness conditions 300, 250, 200, 150 kPa; (d, e) data from n = 24-30 colonies per gelation condition from biological replicates, geometric mean ± 95% confidence interval; statistical significance (unpaired two-tailed t-tests): *p < 0.05, **p < 0.01, ***p < 0.001, ****p < 0.0001. **f.** Schematic of growth curves for resistant (R, red), intermediate (I, blue), and susceptible (S, black) populations under antibiotic inhibition, with a bar chart categorizing colonies into R, I, and S groups by normalization. **g.** Relative growth curves over 6 hours for S (black), I (blue), and R (red) populations at varying kanamycin concentrations; data is presented as mean ± s.d., with average colony size and shaded region (n = 5-30 chambers, independent replicates). **h.** Colony counts, modeled with a Poisson distribution for population density across kanamycin concentrations (128, 16, 8, 4, 2, 1 µg/ml). **i.** Comparison of relative growth at 2 hours (red circles) and 4 hours (blue circles) across kanamycin concentrations (32, 16, 8, 4, 2, 1 µg/ml). **m.** SIR decision-making plots comparing relative growth at 2 hours (red) and 4 hours (blue) for S, I, and R populations, categorizing colonies into distinct SIR profiles. **j-k.** Colony growth measurement at 2 hours and 6 hours across kanamycin concentrations (32, 16, 4, 2, 1, 0.5 µg/ml). (h,j,k) Whisker box plots; median, range within 1.5 IQR, and outliers.

To investigate stiffness effects, we measured *E. coli* colony growth curves over 6 hours of incubation and their antibiotic response to kanamycin (ranging from 0–32 µg/ml) across hydrogels of varying stiffness (150, 200, 250, and 300 kPa)^30^ (Extended Data Fig 6c, Supplementary table 5). Dynamic growth curves of *E. coli* colonies showed that gelation conditions within the 150–300 kPa range did not significantly alter growth rates. Growth arrested at similar levels by 6 hours of incubation. However, higher hydrogel stiffness (>250 kPa) delayed early colonization phases (Fig 4d, Extended Data Fig 6d). Additionally, *E. coli* growth alterations with kanamycin at intermediate susceptibility concentrations (2-4 µg/ml) demonstrated that the hydrogel environment reduces antibiotic efficacy at high stiffness (∼250–300 kPa) (Fig 4e). We confirmed that matrix-driven effects dominate kanamycin resistance in the 2–4 µg/ml range (Extended Data Fig 6e,f). For all subsequent AST experiments, we chose a single gelation condition (∼150 kPa, not oversaturated by Ca^2+^ ions for gelation^29^) to maintain experimental consistency.

### Antibiotic susceptibility decision-making in high throughput

Building on the matrix-stiffness effects, we established a high-throughput AST profile for SIR decisions (Susceptible (S), Intermediate (I), Resistant (R); Fig 4f) with a standard CLSI MIC range up to 128 µg/ml for kanamycin. This analysis was enabled by automated adjustments across thousands of experimental steps including fluid washing, exchange and retention. Growth curves over time indicated no growth for S, moderate growth for I, and full growth for R (Fig 4g), while colony counts increased with decreasing kanamycin concentration (Fig 4h). Analysis of relative growth at varying concentrations (Fig 4i, j) identified an intermediate MIC range of 4-16 µg/ml, with susceptibility confirmed at concentrations exceeding 16 µg/ml of kanamycin. Furthermore, colony comparisons at each condition reinforced SIR decision-making at 2 and 6 hours, respectively (Fig 4k,l). We validated the protective role of the hydrogel, observing decreased kanamycin effects on *E. coli* colonies at a stiffness of ∼150 kPa, which agrees with our matrix-stiffness findings. Timely SIR decisions for clinical isolates could reduce turn-around time of diagnoses^15,40^. Our platform thus offers substantial potential for rapid clinical diagnostics via automated culture and harvest processes through SIR decision-making steps.

### Combinatorial and sequential administration of antibiotics map synergistic drug interactions

Finally, we employed our device and the AST framework to explore combinatorial and sequential antibiotic regimens. We focused on analyzing time-dependent colony effects in simultaneous (A+B) and sequential (A→B or B→A) administration of antibiotics. We tested various scenarios: a single combined antibiotic A+B formulation, sequential delivery with antibiotic A preceding antibiotic B (with a 3-hour delay, *Δt*), and the reverse order (Fig 5a). We characterized *E. coli* under these conditions to elucidate pattern changes between simultaneous and sequential dosing studies. A total of 36 drug pairs derived from 9 antibiotics (listed in Supplementary Table 6) were evaluated under 25 conditions per pair with increasing concentrations, utilizing a 5×5 checkerboard assay (Fig 5b). The evaluation of three dosing strategies at each A+B, A→B, and B→A type combinations resulted in creation of 2,700 distinct drug conditions, which were tested by tracking of a total of 81,000 *E. coli* colonies (30 colonies per condition, 6 different chip experiments).

**Figure 5.**
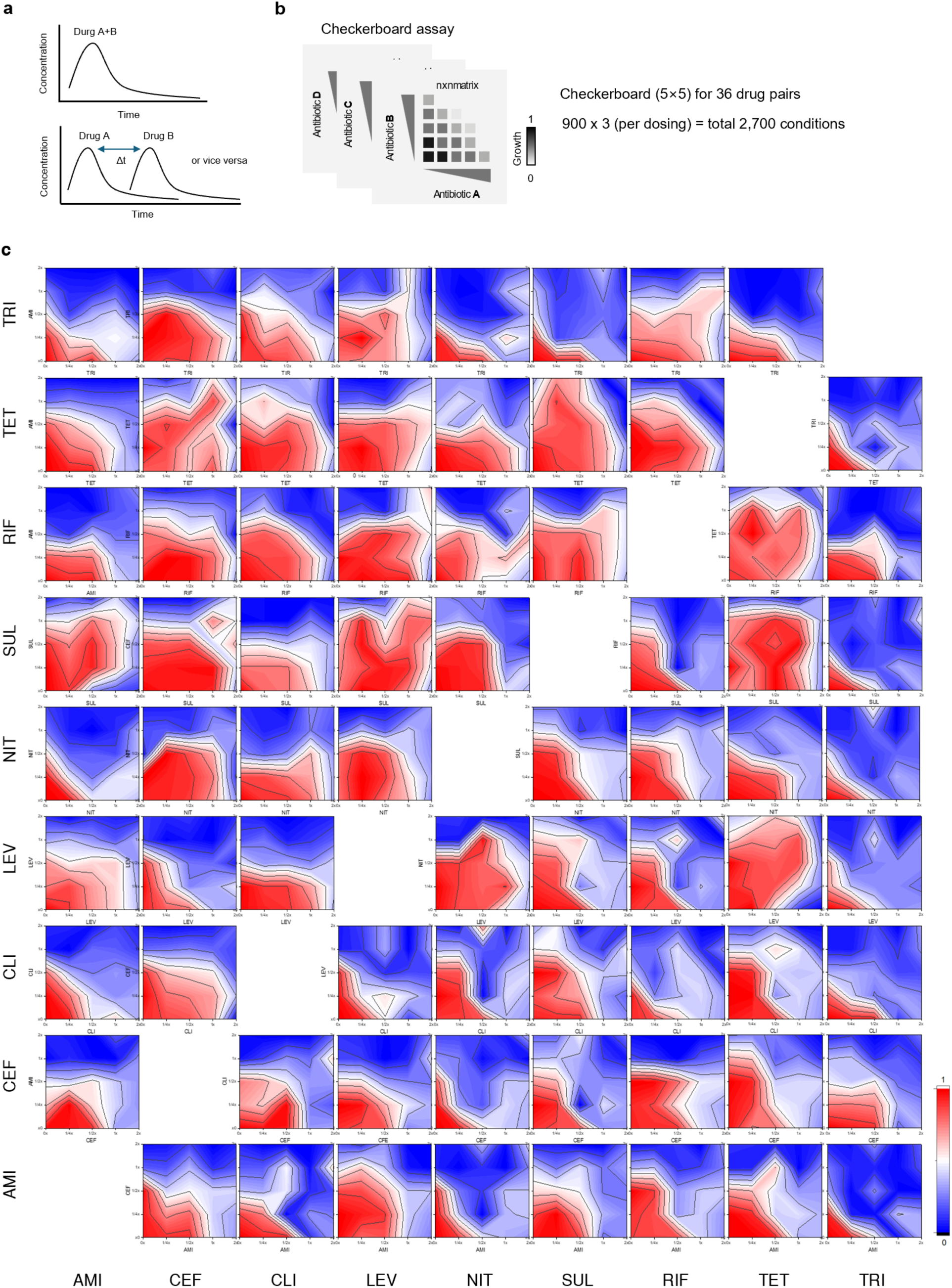
Microfluidic combinatorial and sequential drug administration maps synergistic and antagonistic antibiotic interactions in E. coli. **a.** Schematic of simultaneous administration of antibiotics A and B (A+B), and sequential administration of antibiotics A and B (A → B or B → A) with a 3h time delay, enabling assessment of time-dependent effects on bacterial response. All conditions have been tested in cultures grown in 3D hydrogel islands within the microfluidic device. **b.** Checkerboard assays performed in our device, showing a 5×5 matrix of antibiotic combinations with a gradient representing growth levels (0 = no growth, 1 = full growth). Evaluation is based on 2,700 experimental conditions (25 conditions per drug pair × 36 drug combinations, tested under 3 dosing regimens). Six independent chip experiments have been performed (N=6). The checkerboard assay examines 0, 0.25, 0.5, 1, and 2x concentration for each drug (1x concentration is MIC for each antibiotic, see Supplementary Table 4) **c.** Isobolographic contour heatmaps showing comprehensive growth patterns of E. coli, under a total of 72 pairwise drug conditions with sequential administration of nine antibiotics (AMI, CEF, CLI, LEV, NIT, SUL, RIF, TET, TRI). X-axis: 1st administration, y-axis: 2nd administration. For example, AMI → CEF heatmap contour indicates AMI at x-axis first, followed by CEF at y-axis; CLI → LEV heatmap contour indicates CLI at x-axis first, followed by LEV at y-axis. The color gradient bar indicates growth levels from blue (minimal growth) to red (maximal growth). Overall, 2700 drug conditions were tested in approximately 81,000 *E. coli* colonies tracked over time.

The contoured heatmaps distinctly delineated the temporal sequence effects of A→B and B→A administrations, respectively (Fig 5c), with the result of the A+B administration shown in Extended Data Fig. 7. The contour plots provide a detailed landscape of antibiotic interactions, revealing that sequential administration can enhance synergy (e.g., AMI → NIT) or lead to antagonism between drugs (e.g., TET → LEV) depending on the order and concentration. This data underscores the importance of timing and combination strategies in antibiotic therapy, with potential applications in optimizing treatment protocols based on synergy scores from the analysis of 2,700 distinct antibiotic conditions over time.

### Synergistic and antagonistic shift evaluation of combined drugs

To compare the drug administration conditions, we used a scoring metric to assess synergistic (red arrows) and antagonistic (blue arrows) shifts, based on the Loewe Additivity model^41^ (Fig 6a, Supplementary Note 5). This model converted our high throughput measurements to a quantitative basis for determining whether the combined effect of two drugs exceeds, matches, or falls below the expected additive effect (simple additivity), thus identifying synergy, additivity, or antagonism (Extended Data Fig 8a-d). The calculated combination index (CI) is visually represented through heatmaps, which correspond to three different administration protocols: A+B, A→B and B→A administration, respectively (Fig 6b-c). The heatmap data quantitatively displayed isobolographic patterns to reveal synergistic, additive, and antagonistic interactions, with intensity reflecting synergy or antagonism strength. In addition, these heatmaps were consolidated into a three-tile configuration for a comparative overview of the interaction patterns across the three dosing strategies (Fig 6e). The analysis of 108 antibiotic pairs spanned to highlight the shift patterns of synergy and antagonism, underscoring the significant role of drug administration sequence (Fig 6f).

**Figure 6.**
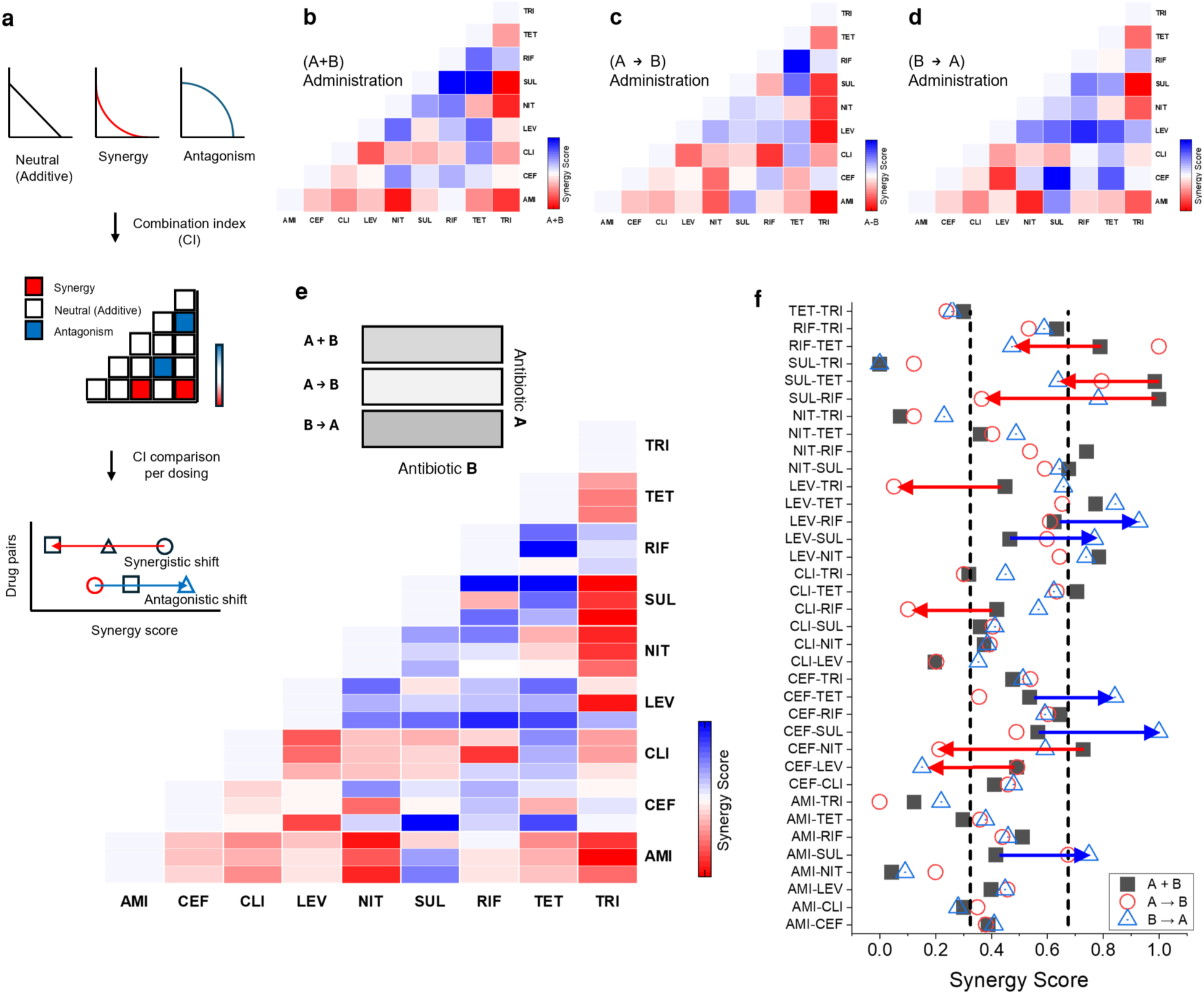
Combination index calculation reveals synergistic and antagonistic shift between drug pairs. **a.** Diagram of combination index (CI) calculation and synergy score evaluation using the Loewe additive model. Isobologram shows neutral/additive (diagonal), synergy (concave), and antagonism (convex). Lower-triangle heatmap indicates synergy score intensity (red: synergy, white: neutral/additive, blue: antagonism). Synergy score shifts include synergistic (red arrow, right to left) and antagonistic (blue arrow, left to right). **b-d.** Lower-triangle heatmaps of 108 pairwise results from nine antibiotics (AMI, CEF, CLI, LEV, NIT, SUL, RIF, TET, TRI); antibiotic A (y-axis), antibiotic B (x-axis), showing the results of (b) A+B simultaneous, (c) A→B sequential, and (d) B→A reverse sequential administration. Color bar indicates synergy (red), neutral/additive (white), and antagonism (blue). (e.g., RIF → TET for synergy, and SUL → CEF for antagonism) **e.** Three-tile heatmap configuration across three administration strategies (top: A+B, middle: A→B, bottom: B→A), showing an overview of administration sequence and timing effects on synergy/antagonism pairwise interactions (see Supplementary Table 6 for details). **f.** Synergy score shift plot for 36 antibiotic pairs across three dosing strategies (symbols: black squares (A+B), red circles (A→B), blue triangles (B→A)), with x-axis showing synergy score (0-1 scale) and y-axis listing 36 combination pairs. Vertical dashed lines mark thresholds (<0.35 synergy, >0.65 antagonism). Twelve pairs highlighted; blue arrows (n=5) indicate antagonism shifts (left to right), red arrows (n=7) indicate synergy shifts (right to left), with score change >0.3 from A+B to A→B or B→A. A scoring metric is detailed in Supplementary Note 5.

A major finding from our data set is that 12 of 36 drug pairs we tested (33.3%) exhibited a score shift >0.3 on a 0-to-1 scale, when simultaneous (A+B) dosing is compared to either sequential A→B or B→A dosing (Extended Data Fig 8e-j). For simultaneous administration, several synergistic and antagonistic drug interactions aligned with previously well-studied drug mechanisms in pairs such as SUL+TRI and AMI+TRI^42,43^. Sequential dosing studies of pairs, mapping synergistic shift for CEF→NIT and antagonistic shift for CEF→SUL, highlighted how administration sequences and timing influence antibiotic efficacy (Supplementary Note 6). *In vivo* host metabolism is often cited as a main factor influencing drug interactions in clinical studies^44^. In our experiments, we have observed strong synergistic or antagonistic effects between drugs even in the absence of host factors. This suggests that factors intrinsic to the pathogens and the drugs alone are sufficient to create many synergistic drug interactions. Our data and the observed shift changes underscores the necessity for in-depth mechanistic studies to unravel the underlying pathogen and drug related biological and chemical processes driving these interactions.

## Discussion

We combined microfluidic Large-Scale Integration^17,18^, soft-materials, and automation to develop a platform for robust microbial 3D cultivation, dynamic drug perturbation and high-throughput phenotypic colony analysis. Advanced functional modules were integrated to perform various drug formulations, cell culture, chemical stimulation, and imaging tasks automatically. Our system incorporates a dense microfluidic architecture and automation for multiplexed generation of tens of thousands of potential chemical stimulation conditions. To create favorable cultivation conditions, we utilized a polysaccharide-based natural hydrogel environment^45^, and solved technical problems in spatially and temporally controlling gel formation and fluidic delivery at the microscale (Fig 1). By integrating high-resolution imaging with automated reagent control, our platform enabled observation of 3D colony dynamics in high throughput (Fig 2 and Fig 3), capturing population growth and spatial interactions often missed in conventional 2D assays^26^.

These capabilities improve the accuracy of cell culture and growth measurements and provide insights into the spatial organization of microbial communities, with potential for understanding complex interactions in realistic 3D environments. We performed extensive studies of antimicrobial susceptibility in physiologically relevant culture conditions. The ability to precisely switch experimental conditions like drug type and concentrations across hundreds of chambers, while dynamically adjusting to varying requirements, makes this platform a powerful tool for advancing microbiology research.

We investigated the impact of nutrient supply timing on microbial growth phases (Fig 2e-n) and found that sequential administration of growth medium at defined intervals exhibited biphasic growth, with an initial lag phase followed by renewed proliferation, which highlights how nutrient availability shapes microbial behavior. This diauxic growth pattern indicates how environmental pressures may encourage the emergence of multidrug-resistant (MDR) pathogens or limit their evolutionary escape routes^46^. Several potential factors may contribute to this outcome including differences in alterations in target enzyme activity and changes in intracellular metabolic processes^47,48^.

Our system enables modulating the culture matrix stiffness to study how mechanical cues shape microbial behavior. We found that changing hydrogel stiffnesses induced differences in microbial growth, and susceptibility to kanamycin (Fig 4). Overall, these insights are consistent with past research highlighting the role of 3D microbial ecosystems (i.e., biofilms^10,39^) in enhancing resistance via physical barriers, altered microenvironments, and physiological alteration.

The timing and sequence of antibiotic delivery are key factors in the complex interplay of pharmacokinetic profiles and susceptibility, which influence treatment efficacy and resistance development^4,7^. Our technology enables systematic screening of dynamic drug interactions in live cultures and maps resistance mechanisms due to pathogen-drug interactions. For example, simultaneous dosing of SUL and RIF exhibited antagonism, whereas sequential dosing (SUL followed by RIF) produced an additive effect. In clinical studies, SUL, metabolized by liver enzymes (CYP2C9)^44,49^, shows reduced plasma concentrations when co-administered with RIF. This co-administration may alter their effectiveness against community-acquired methicillin-resistant *Staphylococcus aureus* (CA-MRSA)^50^. Our extensive studies showed that such drug interactions are observable even in the absence of host factors.

Future incorporation of host cells to create host-pathogen co-cultures would improve the clinical relevance of *in vitro* systems like ours. Our approach is suitable for such improvements, as multiple gel-islands containing different cell types like host and pathogen cells can be created in the same microfluidic chamber (see Supplementary Fig 4). In addition, variations in population density^51^ (number of colonies per chamber) influences population dynamics, which necessitates statistically rigorous exploration such effects. Access to many parallel culture chambers in the same device alleviates this limitation. Finally, image acquisition speed is a limiting factor in high throughput microfluidic experiments, because imaging the entire culture chamber array takes a substantial amount of time, especially when imaging in multiple channels is required. Future advances in high speed and large area microscopy would further improve the temporal resolution or throughput of parallelized microfluidic devices.

In summary, our ultra-multiplexed platform provides complex, dynamic, and sequential drug perturbations to cells in 3D culture and automatically monitors them over time. This is achieved by integrating hydrogel-based culture with automated fluid delivery, real-time analysis, and stable colony cultivation. This capability allows creating microenvironments closer to *in vivo* conditions compared to other platforms, enabling the study of microbial responses to tailored therapeutic regimens. These advancements make our microfluidic platform a transformative tool for dissecting complex microbial interactions and optimizing antibiotic treatment strategies. Future research will elucidate critical mechanisms—such as genetic clone selection, metabolic adaptation, and phenotypic memory—that drive the emergence of resistant subpopulations in 3D microbial communities, and co-cultures of host cells and pathogens, paving the way for innovative therapeutic strategies to combat antimicrobial resistance.

## Methods

The following describes the general methods and analytical approaches employed in this study to investigate bacterial dynamics and antibiotic susceptibility in a high-throughput microfluidic platform. Detailed descriptions of the microfluidic device design, fabrication, bacterial culturing, and synergism evaluation are provided in Supplementary Tables 1-6, Supplementary Notes 1-6 and Supplementary Figs 1–11.

### Microfluidic Device Design and Fabrication

Microfluidic device was fabricated using a well-established soft lithography method, adapted from prior literature^36^ with minor modifications. Molds were created on a 4-inch silicon wafer (WAFERPRO) using negative photoresist (SU-8 3025, MicroChemicals) to form fluid layer channels and control layer features (∼30–35 µm height), and positive photoresist (AZ40XT, MicroChemicals) for pneumatic valve sections (∼20–25 µm height). Patterns, designed in AutoCAD (Autodesk, Inc.) and converted using KLayout (https://www.klayout.de/), were exposed with a 375 nm laser on a Heidelberg MLA150 maskless aligner. Two-layer PDMS (polydimethylsiloxane, 10:1 ratio, Monsanto RTV-615A/B) structure, consisting of control and fluid layers, was mounted on a glass substrate (127.8 × 85.5 × 1 mm, Marienfeld), with fluid channels (∼150 µm wide) and 18 input points enabling multiplexed fluidic control. Fabrication involved pouring ∼70 g for the control layer and ∼10 g of PDMS for the fluid layer, spun at 2,200–2,300 RPM to achieve a 50–60 µm thickness, and cured at 80°C overnight for 12 hours. Fluid inputs and outputs were punched. The fluid layer was aligned with the control layer using a custom stereomicroscope equipped with an XYZ translation stage and plasma-bonded (Harrick, PDC-001) to the glass substrate.

### Multiplexer Control

Combinatorial multiplexer was evaluated for fluid trafficking and n-fold dilution capabilities within the microfluidic chip, utilizing nine distinct mixtures in this study. Four food dyes (McCormick; red, green, blue, and yellow) were introduced at nine input nodes, each assigned to a unique color, and controlled via a custom MATLAB-based graphical user interface (GUI, MATLAB R2024b). Fluid dynamics were visualized using brightfield imaging on an inverted optical microscope (Nikon ECLIPSE Ts2R), enabling qualitative assessment of dye mixing and flow patterns. Dilution gradients, including linear and serial dilutions with varied mixing ratios, were quantified using 10 µM Cyanine5 (Cy5, Invitrogen) mixed with deionized (DI) water. Fluorescence microscopy (50 ms exposure, Nikon Ti-E Eclipse, RFP filter: 600 nm excitation/50 nm bandwidth, 665 nm emission/50 nm bandwidth) measured diluted fluorescence levels at the chip’s outlet. Fluorescent intensities (arbitrary units) were quantified via line scans or regions of interest, normalized to the maximum input fluorescence, ensuring precise control of targeted concentrations for antibiotic delivery.

### Microorganisms and Growth Conditions

Microbial strains used in this study were *Escherichia coli* Nissle 1917, *E. coli* DH5α, GFP/RFP-expressing *E. coli* DH5α (provided by Dr. Mark Mimee, University of Chicago), *Streptococcus pneumoniae* (ATCC 49619), *Pseudomonas aeruginosa* (ATCC 10145), and *Klebsiella pneumoniae* (ATCC 13883). Cultures were grown to mid-exponential phase (OD₆₀₀ ≈ 0.5–1.0, depending on the strain) in Luria-Bertani (LB) broth (pH 7.0, Fisher, BP1426) or Cation-adjusted Mueller-Hinton II (MH2) broth (pH 7.3, MilliporeSigma, 90922) at 37°C. Stocks were stored at-80°C in tryptic soy broth (MilliporeSigma, 41298) with 20% glycerol. Bacterial strains were revived from-80°C glycerol stocks by streaking onto LB or MH2 agar plates (1.5% agar) and incubating at 37°C for 12–16 hours. A single colony was inoculated into 5 mL of LB or MH2 broth in a sterile 15-mL conical tube and incubated at 37°C with shaking (200 rpm) for 6–12 hours. Optical density was measured using a spectrophotometer (Thermo Scientific, GENESYS 20).

### Bacterial Loading and Hydrogel gelation

Bacterial cultures were diluted 1:100 into 10 mL of fresh LB or MH2 broth and grown at 37°C with shaking (200 rpm) until reaching mid-exponential phase (OD₆₀₀ ≈ 0.5). Bacterial concentration (cfu/mL) was estimated using the standard plate colony counting method. A 1–2% (w/v) alginate hydrogel solution (alginic acid sodium salt, Sigma-Aldrich, 180947) was prepared in deionized (DI) water. A 10–25 µL aliquot of bacterial broth with targeted concentrations was mixed with 1 mL of 1–2% (w/v) alginate solution. Chip surfaces were cleaned and filled with DI water to remove air bubbles. Three 2 mL input vials (DI water, CaCl₂ [Sigma-Aldrich, C4901], and sodium citrate [Sigma-Aldrich, S4641]) were connected to additional input lines in the chip and pressurized at 2–5 psi. 1–2% (w/v) alginate solution with bacterial inoculums was loaded by reverse flow for 2–3 minutes (see Supplementary Fig 5), and pneumatic valve actuation was used to trap the inoculums. A 0.1 M CaCl₂ solution was introduced to induce gelation for 3–10 seconds in the inner chambers. After gelation, the bacterial inoculum was cultivated at 25–37°C.

Chamber valve actuation, bacterial loading, gelation conditions, and media exchange processes are followed by Extended Data Fig 3b,d. Additional descriptions are detailed in Supplementary Note 1-2 with Supplementary Table 1,2.

### Automation Experimental Setup

Experimental workflow integrated reagent retention, fluid trafficking, time-lapse imaging, automated image analysis, and data storage, conducted on an inverted fluorescence microscope (Nikon Ti-Eclipse) under controlled conditions (25–37°C, >95% humidity, ambient air). The system was mounted in a culture chamber equipped with a microscope stage controller to maintain stable temperature and humidity, ensuring consistent experimental outcomes, as detailed in Extended Data Fig 1. Automation was achieved through a custom MATLAB-based GUI (MATLAB R2016b and R2024b, available at [https://github.com/Uchicago-TAY/LIST]) controlling a valve pneumatic solenoid manifold (Festo, 197334) connected to electronic control units. The manifold, operated via a USB controller box with a 5–40 psi pressure range, actuated the chip’s control layer and was connected to control lines using water-filled Tygon tubing (0.02-inch ID × 0.06-inch OD, VWR). Input manifolds for reagent delivery, linked to low air pressure (2–10 psi), pressurized reagent 1ml vials. The chip was mounted on the microscope, cleaned with a razor blade and scotch tape to ensure optical clarity, and rigorously tested for leaks and defects at 35 psi. To test full automation from reagent loading to washing steps, a checkerboard-like pattern was created to demonstrate environment switching across all 512 chambers in a 32×16 array, with each chamber filled with distinct food color dyes.

### Automated Image Analysis

Time-lapse imaging was conducted using Nikon’s Ti2 control software (µManager, Nikon), capturing images of chambers every 5 minutes with 20×, 40×, and 60× objectives for phase-contrast and GFP/RFP fluorescence (GFP filter: 450 nm excitation/40 nm bandwidth, 500 nm emission/50 nm bandwidth; RFP filter: 600 nm excitation/50 nm bandwidth, 665 nm emission/50 nm bandwidth). The Nikon Ti2 microscope was equipped with an LED source (Lumencor Spectra X) and a CMOS camera (Hamamatsu ORCA-Flash4.0 V2). The Nikon’s perfect focus system ensured consistent focus with the 40× objective. Bacterial inoculums were initially fed at 0 hours and, after 3 hours of incubation, programmed through the GUI to replace the outer chamber volume (∼20 nL) with LB or MH2 broth via on-chip peristaltic valve actuation. Growth dynamics were monitored through phase-contrast imaging to maintain stable culture conditions. The GUI enabled chamber-specific delivery, with baseline images captured before antibiotic delivery and continuous imaging every 5 minutes per chamber in a cycle during exposure to track bacterial responses. Raw images were processed using a custom MATLAB algorithm for 3D volume rendering and 2D segmented analysis, applying an intensity threshold to delineate bacterial colonies (see Extended Data Fig 4). MATLAB scripts tracked growth trajectories of multiple colonies, analyzing changes in colony count and size to evaluate bacterial dynamics and antibiotic effects. Relative growth on growth curve data was normalized using the pipeline’s area measurement on a 0-1 scale. The 0-1 scale, which is derived from control groups, is used to compare each experiment.

### Antibiotic Susceptibility Testing (AST) in microfluidics

Antibiotic susceptibility was assessed for four strains: *E. coli* Nissle 1917, *S. pneumoniae*, *P. aeruginosa*, and *K. pneumoniae* with 10 antibiotics (amikacin [Sigma-Aldrich, A3650], cefepime [Sigma-Aldrich, PHR1763], clindamycin [Sigma-Aldrich, PHR1159], levofloxacin [Sigma-Aldrich, 28266], nitrofurantoin [Sigma-Aldrich, 46502], sulfamethoxazole [Sigma-Aldrich, 31737], rifampin [Sigma-Aldrich, R3501], tetracycline [Sigma-Aldrich, T7660], trimethoprim [Sigma-Aldrich, 46984], and kanamycin [Sigma-Aldrich, K1377]) (as detailed in Supplementary Table 3, except for kanamycin). Microfluidic minimum inhibitory concentrations (MICs) were determined by the concentration range (CLSI guidelines^52^, 0.5–256 µg/mL) for 6–9 hours of incubation in the chip using cationic adjusted MH2 broth. Modified MATLAB scripts tracked growth trajectories of at least 30 colonies per conditions, as detailed in each figure legend with experimental conditions (hydrogel stiffness, incubation time, colony numbers), analyzing changes in colony count and size to evaluate bacterial dynamics and antibiotic effects.

The MIC breakpoints for each antibiotic and strain were compared in Supplementary Table 4 (this study vs. CLSI database)^37^.

### Sequential Dosing Experiments and Synergy evaluation

Simultaneous and sequential dosing studies in the chip assessed 36 antibiotic pairs, from 9 antibiotics (listed in Supplementary Table 6), using a 5×5 checkerboard assay with concentrations of 2x, 1x, 0.5x, 0.25x, and 0x, yielding 25 conditions per pair. Microfluidic MICs (1x) for each antibiotic were obtained from AST experiments for *E. coli*. Bacterial growth was monitored for 9 hours via phase-contrast imaging, employing microfluidic AST methods to evaluate combination effects. In sequential dosing studies, the first antibiotic was administered at 0 hours, followed by a media exchange to MH2 broth containing the second antibiotic after 3 hours, with growth tracking for an additional 6 hours of incubation. Colony population density and growth rates were quantified using a customed MATLAB-based algorithm. Additional descriptions for combinational index (CI) and synergism evaluation with modifications are provided in Supplementary Note 5.

### Software and statistics

All schematic figures were generated using Adobe Illustrator (CC2018) and AutoCAD (v.2025) for producing technical schematics related to microfluidic system design or experimental setups. Data analysis and visualization were performed using MATLAB (v.R2024b) and Origin (v.2025) to complement MATLAB outputs. Visual Studio (v1.1) with Graphviz was used for supplementary data visualization, creating structured diagrams. All image-processing algorithms, used to analyze phase-contrast images from microfluidic AST experiments for quantifying bacterial growth metrics, were developed in MATLAB (v.R2024b). Statistical analyses were performed using MATLAB (v.R2024b) software. For all studies, n.s. indicates not significant (p > 0.05), *p < 0.05, **p < 0.01, ***p < 0.001, ****p < 0.0001. Unpaired or paired student’s *t* test was used to compare data from two groups. All tests were two-sided. Details of specific statistical methods and p-value results are included within each figure legend.

## Data Availability

All data supporting the findings of this study are included in the Main Text and Supplementary Information. Additional data are available from the corresponding author upon reasonable request. Source data are provided with this paper.

## Code Availability

AutoCAD design files, GUI function codes, Experimental scripts, and colony-tracking pipeline are publicly available on GitHub (https://github.com/Uchicago-TAY/LIST).

## Acknowledgement

This work was supported by the NIH grant R35GM148231 (S.T.) and Chan-Zuckerbeg Chicago Biohub Investigator Award (S.T.).

## Author contributions

Y.J. conceptualized, designed microfluidic platform and performed experiments, analyzed data, and wrote the first draft of the manuscript. G.M. and A.P. assisted with microfluidic experiments and data analysis. Y.D., M.P., J. M., and S. O. assisted with chip design for multiplexing features and provided experimental resources. E.T assisted with antibiotic susceptibility experiments. S.T. conceptualized, supported and supervised the project. All authors reviewed and edited the manuscript.

---

**Extended Data Fig 1.**
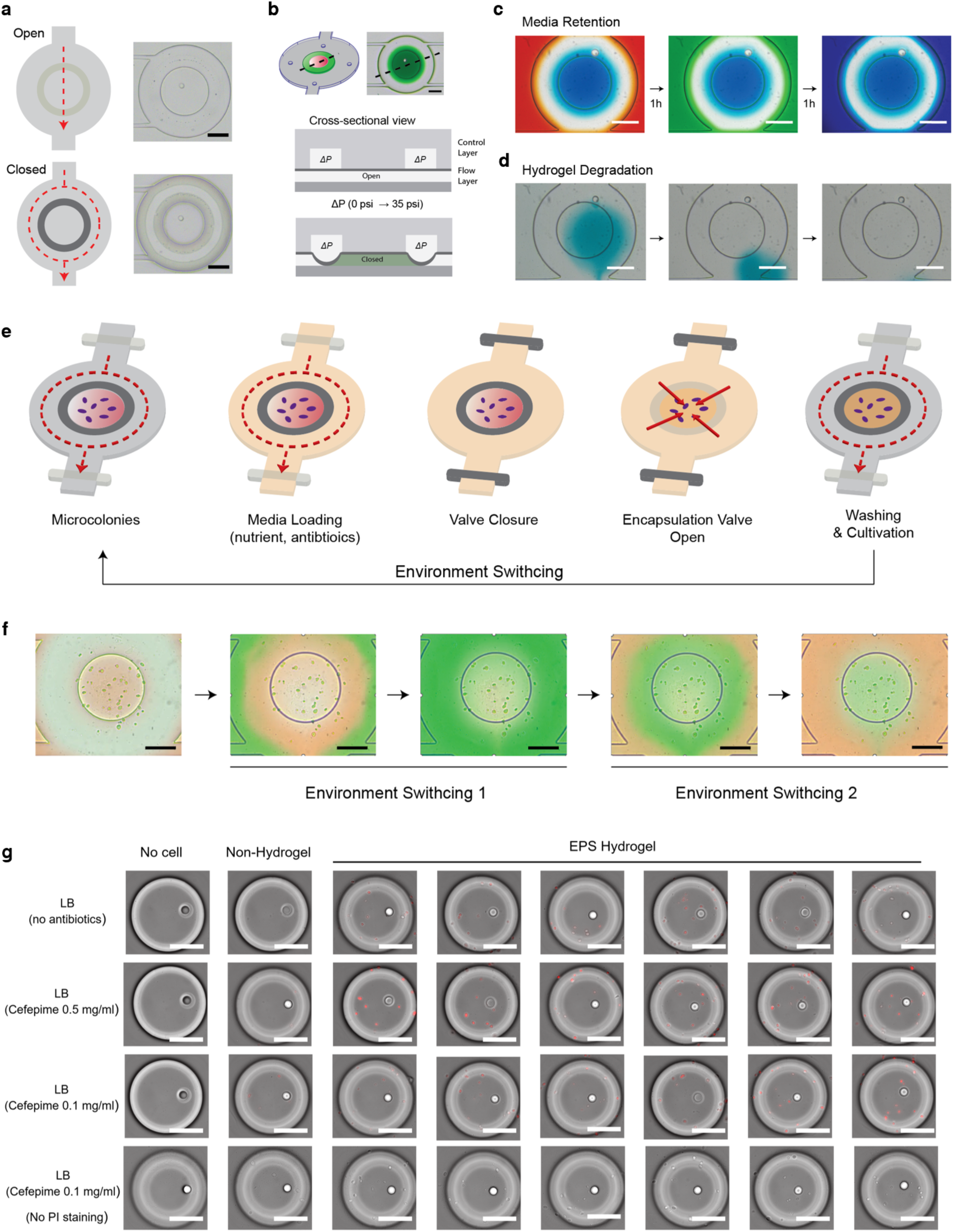
Inner chamber compartmentalization and hydrogel functionality for temporal administration of molecules of interest. **a.** Encapsulation-valve control mechanism (upper: open state; lower: closed state) in a microfluidic chamber, with microscopic images (right) operating at pressures from 0 to 35 psi; scale bar, 100 µm. **b.** Chamber view with cross-sectional representation, showing a thick control PDMS layer and thin flow PDMS layer mounted on a glass substrate; pressure differential (ΔP: 0 psi to 35 psi) controls open and closed states (top: black dashed line indicates cross-section), with microscopic images of the inner chamber state containing green-colored media; scale bar, 100 µm. **c.** Media retention capacity demonstrating no media diffusion across the encapsulation valve upon closure, with each image showing 1-hour retention and transitions in the outer chamber region to test complete compartmentalization; scale bar, 100 µm. **d.** Alginate hydrogel degradation induced by Ca²⁺ ion chelators (e.g., EDTA, sodium citrate solution) for decrosslinking; time-lapse images indicate blue food dye-stained alginate hydrogel degrades within seconds under flow conditions; scale bar, 100 µm. **e.** Schematic of environmental switching in the chamber, including media loading (nutrients, antibiotics) in the outer chamber region, valve closure, encapsulation valve opening for inward molecular diffusion, and washing step for cultivation. **f.** Time-series images of repeated environment switching in a culture chamber containing *E. coli* colonies, with green-colored solution (Switching 1) and red-colored media (Switching 2) administration, highlighting inward molecular diffusion from the boundary to the center of the hydrogel and no disturbance of the hydrogel structure containing *E. coli* colonies; scale bar, 100 µm. **g.** Images of *E. coli* cultures after 5-minute Cefepime and PI administration: no cells, non-hydrogel, and EPS hydrogel with Cefepime (0.5 mg/ml and 0.1 mg/ml, with and without PI staining); scale bar, 100 µm.

**Extended Data Fig 2.**
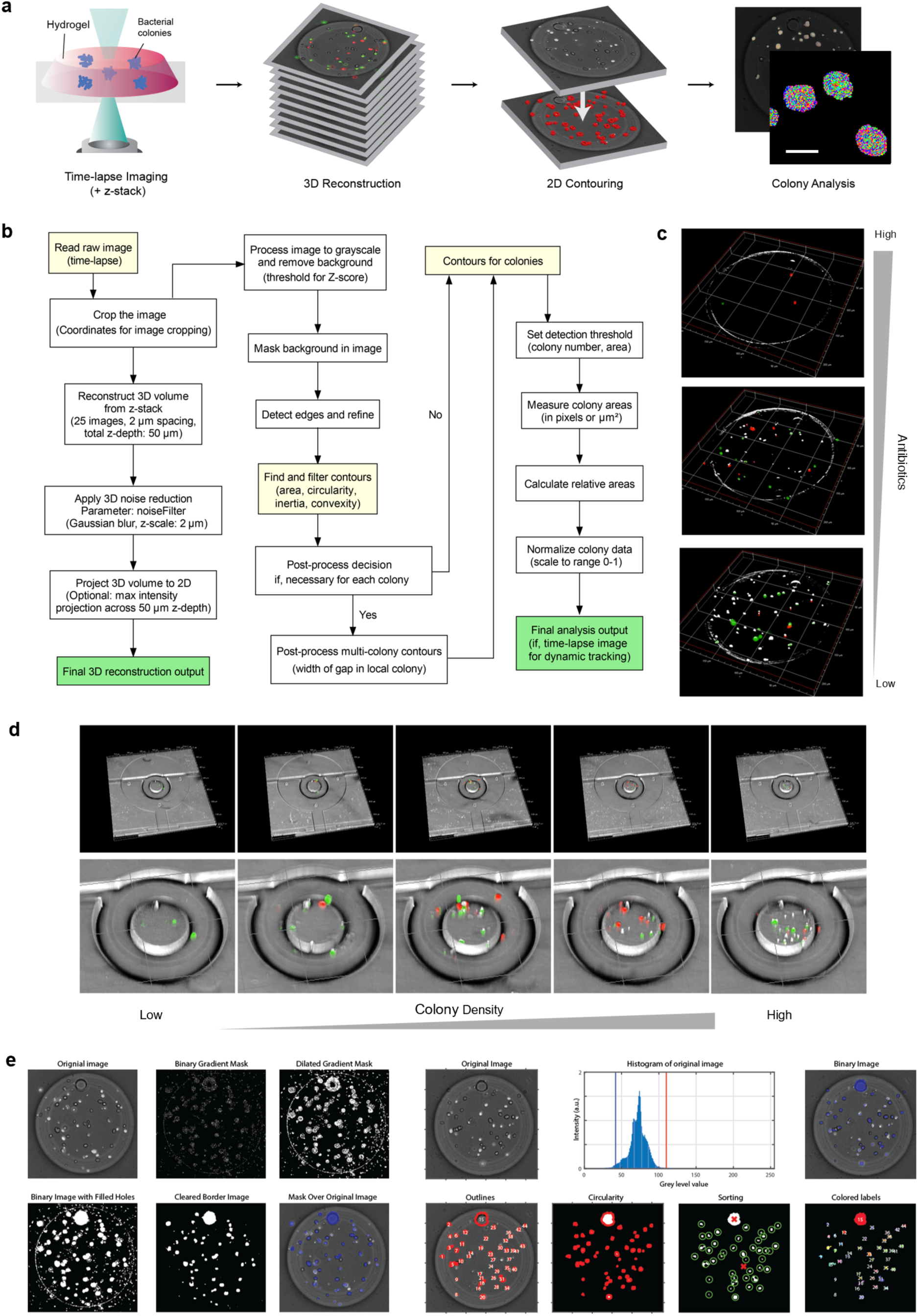
Colony-tracking high-throughput phenotyping and quantitative analysis. **a.** Schematic workflow for colony tracking analysis, starting with time-lapse imaging of a hydrogel with bacterial colonies, followed by 3D reconstruction from z-stack images, 2D contouring, and colony analysis; scale bar, 10 µm. **b.** Flowchart diagram for processing images, including reading time-lapse data, cropping, 3D volume reconstruction, background masking, edge detection, contour finding (area, perimeter, inertia, convexity), and final 3D reconstruction output. It also includes post-processing steps for multi-colony contours if necessary. **c.** 3D reconstructed images of E. coli colonies in a chamber, processed through the pipeline at varying Kanamycin concentrations (top to bottom: 16, 2, and 0 µg/ml), with grey (wild-type), green (gfp-expressing), and red (rfp-expressing) dots representing different E. coli bacterial populations. **d.** 3D chamber views with bacterial colonies with varying population density with a gradient bar (low to high). Grey dots represent wild-type E. coli DH5α bacterial colonies, green dots indicate gfp-expressing E. coli DH5α, and red dots denote rfp-expressing E. coli DH5α. **e.** Intermediate steps of 2D image processing for microbial colony detection, providing a visual representation of how raw images are processed to isolate and segment colonies. The panels demonstrate the transformation of a raw image into a binary image, followed by contour detection and sorting based on geometric features such as circularity. This process aligns with the pipeline steps (e.g., “Mask background in image,” “Find contours,” and “Pre-filter contours”). Detailed flowchart pipeline for processing raw images of microbial colonies, obtained by the platform, is provided in Supplementary Note 3.

**Extended Data Fig 3.**
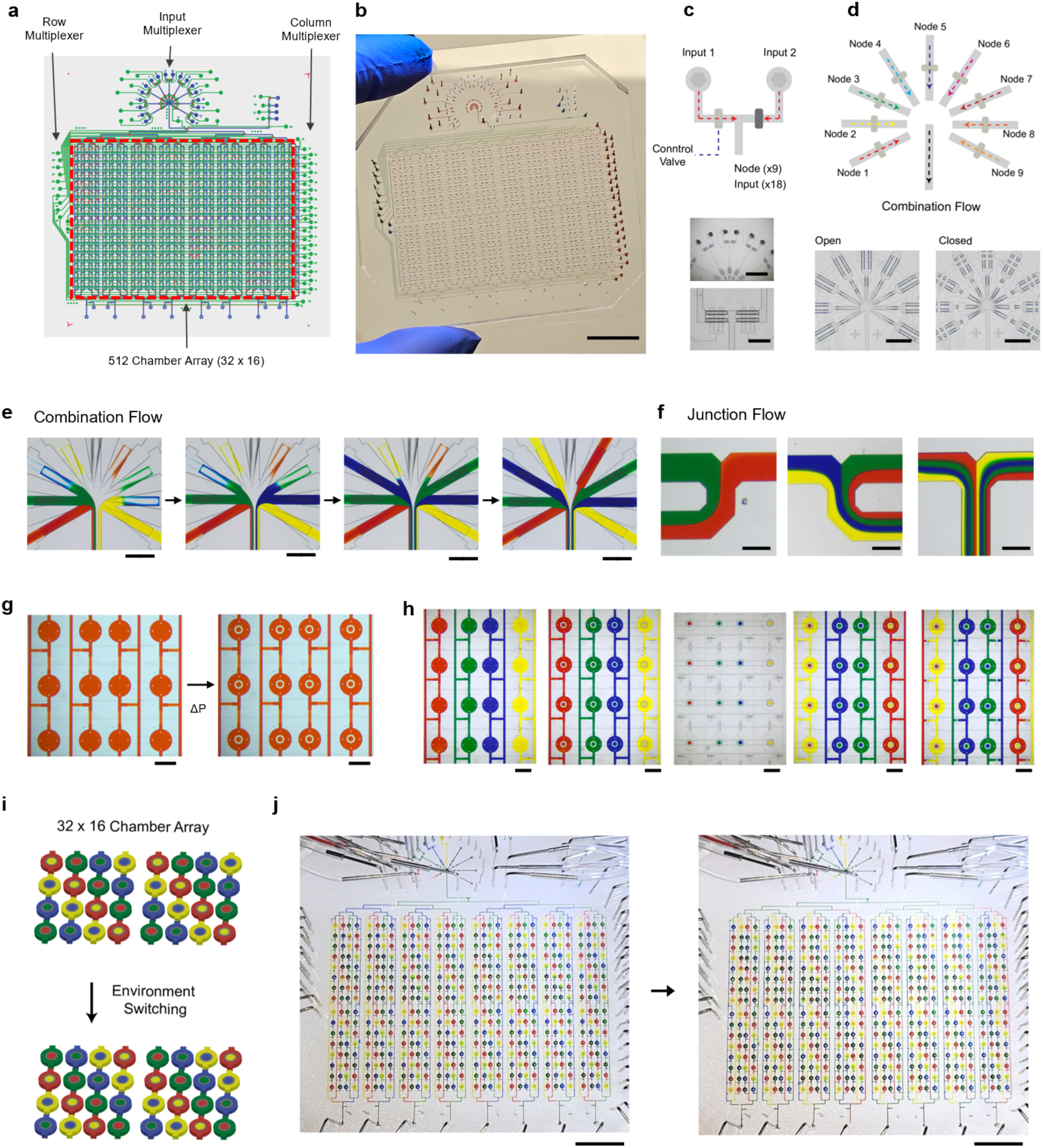
Large-scale Integration for ultra-multiplexed microfluidics system. **a.** Whole-chip layout from original AutoCAD design. Black arrows indicate row/column and input multiplexer configuration and a 512-chamber array (32 × 16) outlined in a red box. **b.** Representative image of the integrated microfluidic device; scale bar, 1.5 cm. **c.** Two inputs connected to a node via pneumatic valves; microscopic images of input nodes (middle image; scale bar, 5 mm; lower image; scale bar, 1 mm). **d.** Input system design layout of the combination multiplexer, with 9 colored input lines (totaling 18 inputs) connected to 2 nodes each, arranged radially around a central outlet for delivering reagent mixtures to culture chambers. Microscopic images show open and closed states by valve actuation; scale bar, 1 mm. **e.** Sequential images of combination fluids (from left to right: 4, 5, 7, and 9 reagent mixtures) with different colored input channels converging into a central outlet, where compounds are mixed and delivered. Each line can connect to a different compound or reagent (up to 18 inputs) to support combination testing; scale bar, 500 μm. **f.** Junction flow images with colored fluids (from left to right: 2, 4, and 8 reagent mixtures) combining at intersection points; scale bar, 150 μm. **g.** Configuration for inner-chamber compartmentalization, showing sequential operation of inner-chambers under varying pressure (ΔP, 0 to 35 psi); scale bar, 1 mm. **h.** Multiplexing chamber configuration with microscopic images of a 4 × 4 culture chamber array; scale bar, 1 mm. **i.** Illustration of a checkerboard-like pattern demonstrating environment switching across all 512 chambers in a 32 × 16 array. **j.** Image of the whole device with all 512-chamber switched by distinct food color dyes; scale bar, 1 cm.

**Extended Data Fig 4.**
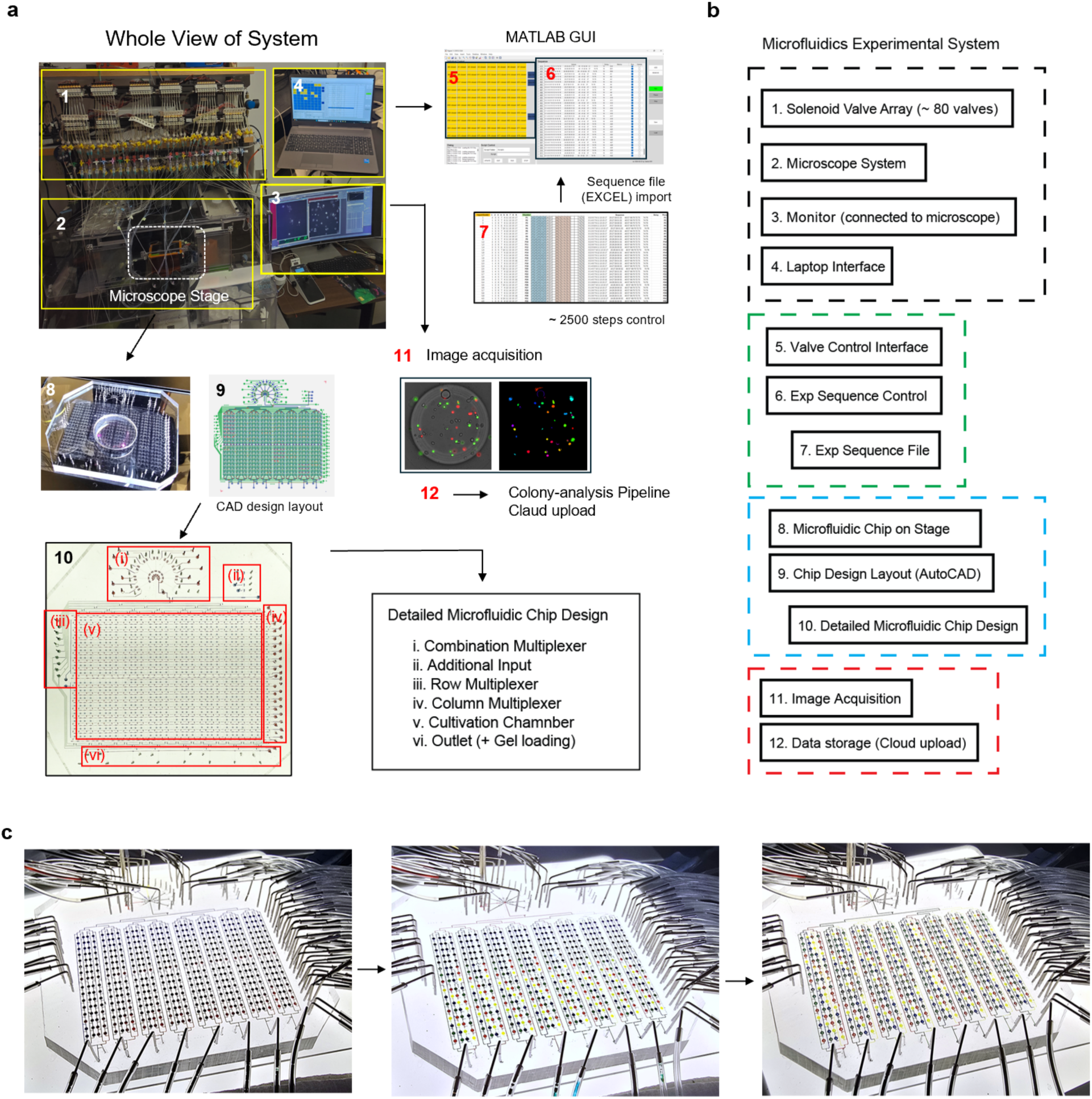
Operation System for Integrated Hardware, Software, and Data Management with Full Automation. **a.** Experimental setup utilizing 80 solenoids to actuate 1,420 valves across 512 chambers. (1) Solenoid valve array (up to 80 valves) pressurized by an air source (0–40 psi), (2) Fluorescence microscope (Nikon Ti-Eclipse) with an incubation system (∼37°C), (3) Monitor connected to the microscope, (4) Laptop interface hosting the MATLAB GUI, (5) Solenoid valve control interface via GUI, (6) Experimental sequence control (up to 2,500 steps in this study), (7) Excel sequence file (“Combinatorial Inputs.xlsx,” detailed in Supplementary File), (8) chip on microscope stage, (9) AutoCAD chip design layout, (10) Image of chip with detailed design components: (i) combination multiplexer, (ii) additional input, (iii) row multiplexer, (iv) column multiplexer, (v) outlet chamber, (vi) outlet for gel loading, and (vii) an implied additional outlet chamber, (11) Image acquisition process, and (12) Data storage with cloud upload for colony-tracking analysis. **b.** Overview of system workflow. The process proceeds sequentially: Excel file import from (7) to (5–6), valve control via GUI from (5– 6) to (1), image acquisition from (2) and (8) to (11), and cloud upload from (11) to (12) for data analysis. **c.** Time-lapse operational images of the chip, demonstrating environmental switching controlled by the sequence file (provided in Supplementary File). Images depict a filling checkerboard pattern starting with columns P1 to P32, followed by rows O1 to O32, and then A1 to A32, each filled with distinct color patterns (red, blue, yellow, green). These colors likely represent different reagents or antibiotic combinations delivered to each chamber via the multiplexer, enabling control of 512 unique combinations.

**Extended Data Fig 5.**
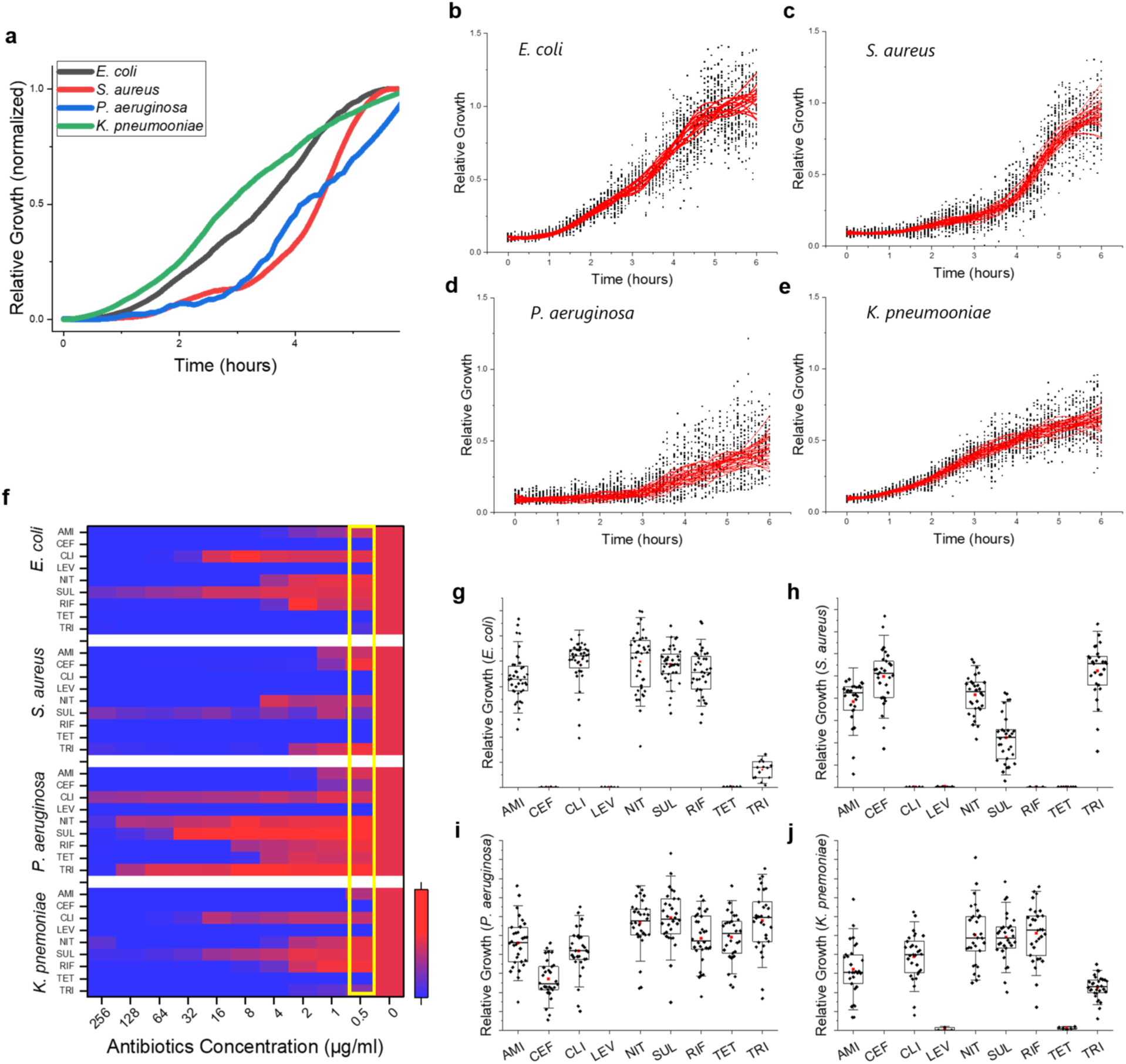
Bacterial colony growth measurement and susceptibility test for four bacterial species. **a.** Normalized growth curves over 6 hours for four bacterial species (E. coli, S. aureus, P. aeruginosa, K. pneumoniae) cultivated in MH2 media without antibiotics. **b-e.** Scatter plots with overlaid mean curves showing relative growth over time for each species (b: E. coli, c: S. aureus, d: P. aeruginosa, e: K. pneumoniae), based on analysis of at least 30 colonies (n ≥ 30, biological replicates). **f.** Susceptibility testing heatmap displaying growth responses of four species (top to bottom: E. coli, S. aureus, P. aeruginosa, K. pneumoniae) across nine antibiotics (AMI, CEF, CLI, LEV, NIT, SUL, RIF, TET, TRI) at concentrations of 256, 128, 64, 32, 16, 8, 4, 2, 1, 0.5, and 0 µg/ml. Growth levels are represented by a color gradient from blue (high susceptibility, minimal growth) to red (resistance, maximal growth). **g.** Relative colony growth for each species (top left to bottom right: E. coli, S. aureus, P. aeruginosa, K. pneumoniae) at 6 hours (yellow box in **f**) for an antibiotic concentration of 0.5 µg/ml across the nine antibiotics; Whisker box plots showing median, range within 1.5 interquartile range (IQR), and outliers. (**g**) Statistical significance is omitted due to the large number of pairwise comparisons.

**Extended Data Fig 6.**
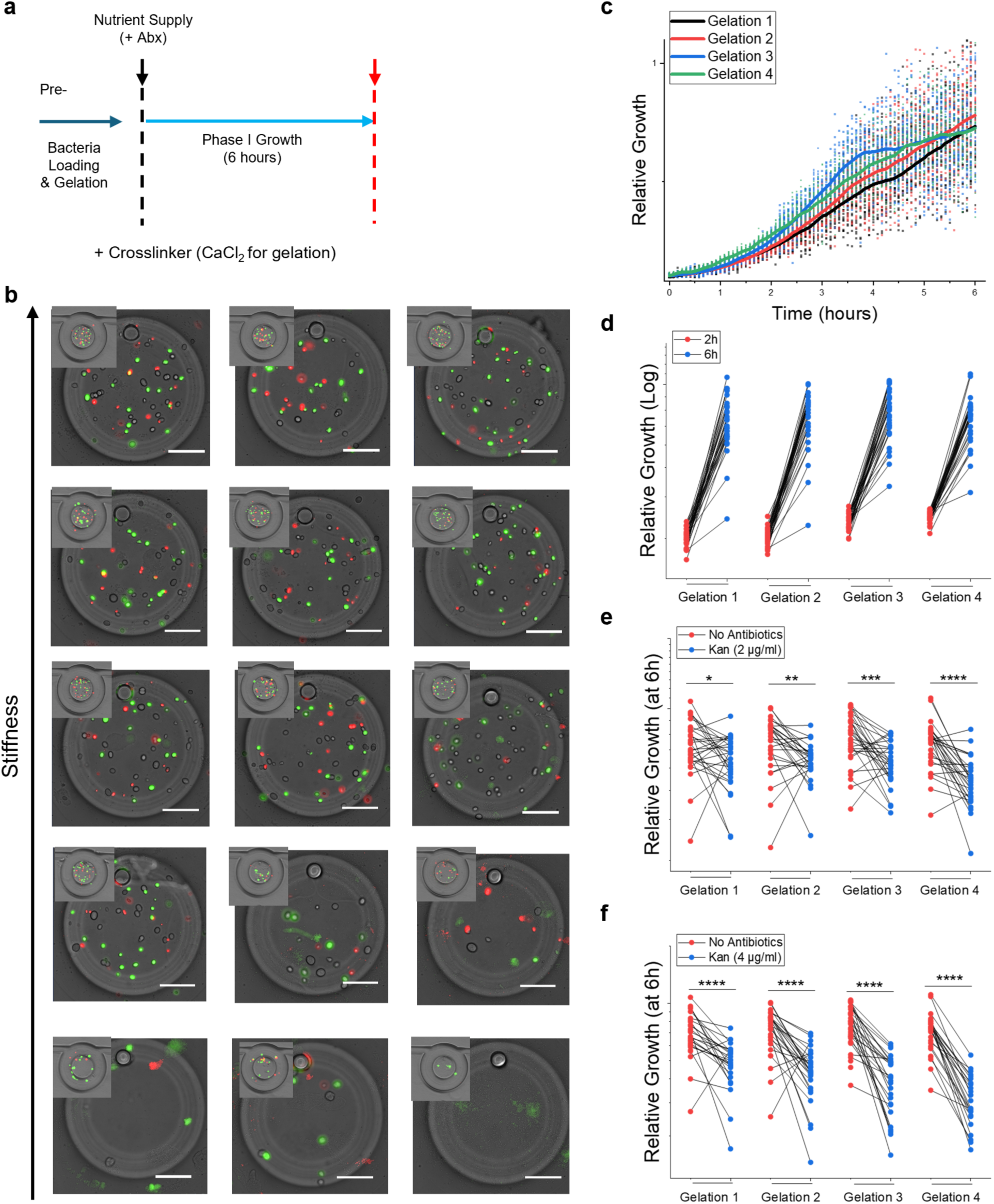
Hydrogel stiffness effect on kanamycin susceptibility testing. **a.** Experimental workflow, including pre-loading of bacteria and gelation with cross-linker (Ca²⁺), followed by fresh MH2 medium supply during Phase I (6-hour growth phase) and image acquisition. **b.** Fluorescent microscopy images of chambers showing bacterial growth patterns (E. coli: wild-type, gfp-expressing, rfp-expressing) in 3D environments with varying hydrogel stiffness (top to bottom: 300, 200, 150, 50, 10 kPa), controlled by Ca²⁺ concentration. Scale bar, 50 μm. **c.** Scatter plots of colony growth tracking at 6 hours under four gelation conditions (Gelation 1–4: 300, 250, 200, 150 kPa), with average colony size represented by colored lines and individual data points. **d.** Line plots comparing relative growth for Gelation 1–4 (stiffness: 300, 250, 200, 150 kPa) at 2 hours (red dots) and 6 hours (blue dots) without antibiotics, highlighting growth variability. **e,f**. Line plots comparing relative growth for Gelation 1–4 (stiffness: 300, 250, 200, 150 kPa) with (red) and without (blue) kanamycin at 6 hours: (**e**) 2 μg/ml, (**f**) 4 μg/ml, showing the association between hydrogel stiffness and antimicrobial effect. (**c-f**) Data based on 25– 30 colonies per gelation condition (n = 8–10 colonies per hydrogel, biological replicates). Significant differences (unpaired two-tailed t-tests) are marked as *p < 0.05, **p < 0.01, ***p < 0.001, ****p < 0.0001.

**Extended Data Fig 7.**
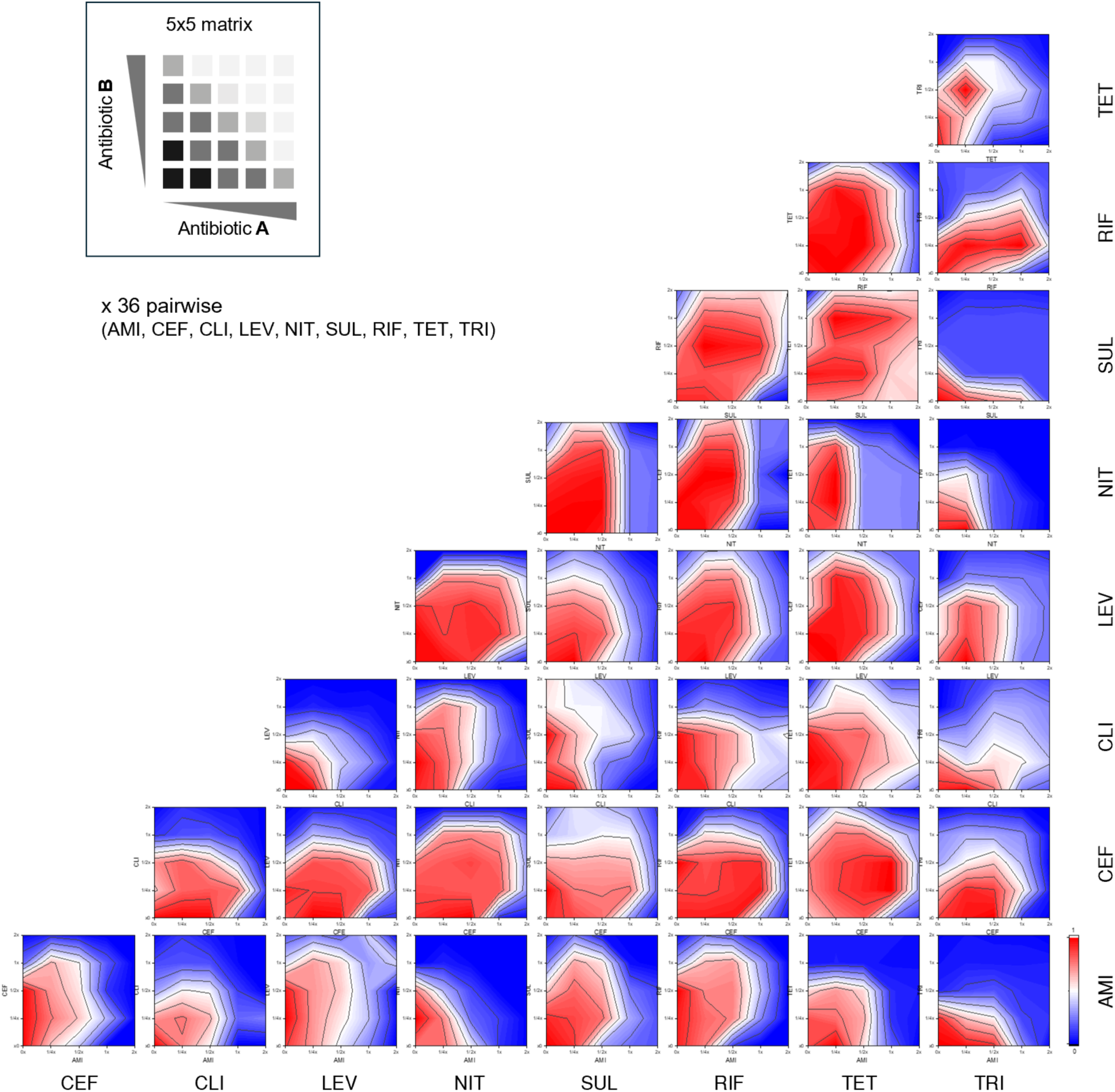
Microfluidic checkerboard assay for simultaneous administration. Isobolographic contour heatmap data illustrating comprehensive growth patterns from 36 pairwise combinations resulting from simultaneous administration of nine antibiotics (AMI, CEF, CLI, LEV, NIT, SUL, RIF, TET, TRI). A color gradient bar indicates growth levels, ranging from blue (minimal growth) to red (maximal growth); Data measured by a microfluidic checkerboard assay, presenting a 5×5 matrix of antibiotic combinations with varying concentration (0, 0.25, 0,5 1, and 2x MIC for each antibiotic).

**Extended Data Fig 8.**
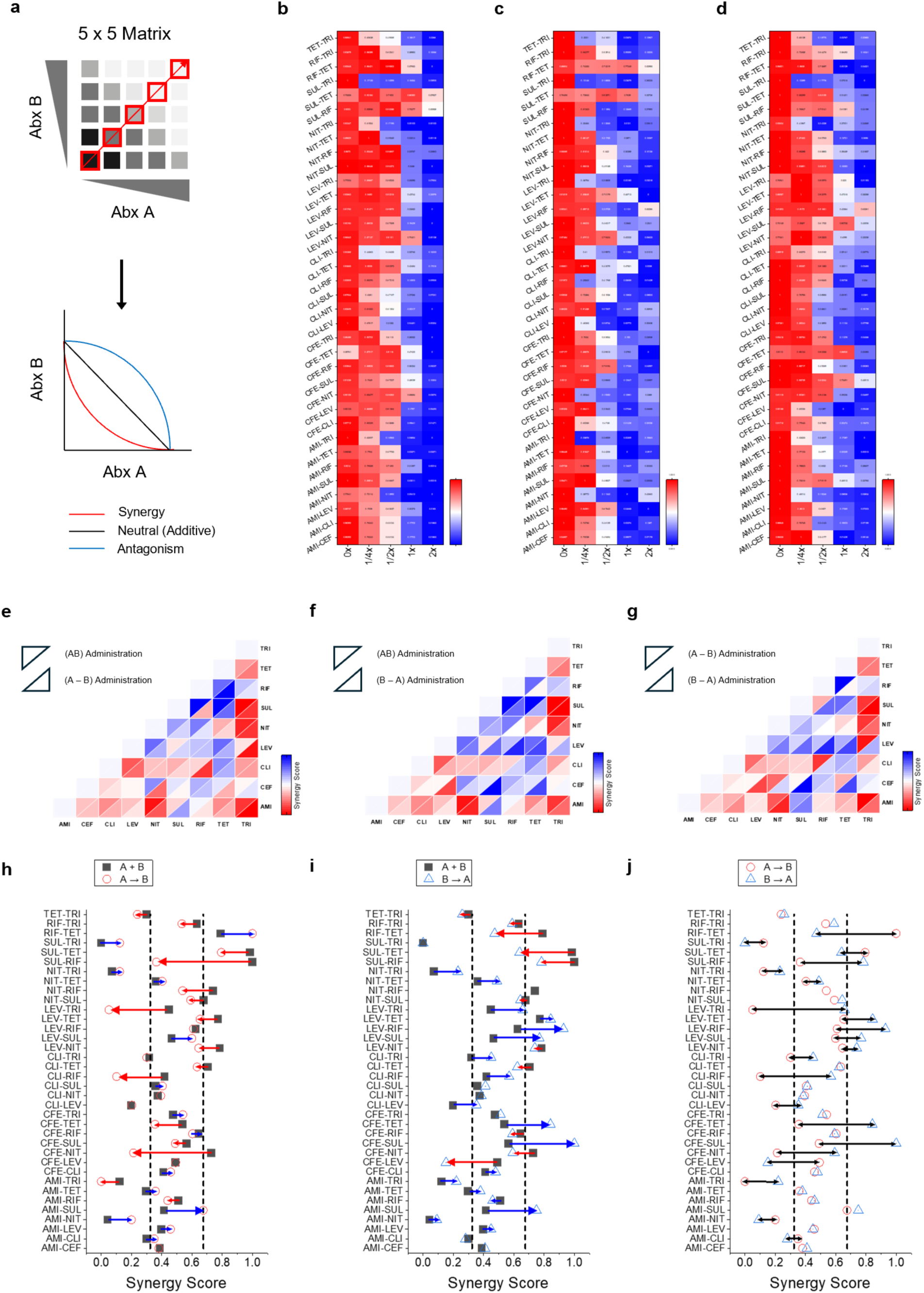
Combination index calculation and comparison of synergy score shift. **a.** Diagonal analysis of a 5×5 checkerboard matrix to approximate isoboles and combination index (CI) calculation. **b-d,** Heatmaps from diagonal analysis for all 36 pairwise combinations of nine antibiotics, with a color gradient from blue (low growth) to red (high growth), highlighting 1:1 two-drug dose responses (red arrow along diagonal calculation in **a**). **e-g.** Comparative heatmaps of synergy scores with two tiles: upper triangle (A+B), lower triangle (A→B) in (**e**); upper triangle (A→B), lower triangle (B→A) in (**f**); upper triangle (A+B), lower triangle (B→A) in (**g**). **h-j.** Synergy score distribution plots with x-axis showing synergy scores (0-1 scale) and y-axis listing pairwise combinations; vertical dashed lines mark thresholds (0.65 antagonism). Symbols: black squares (A+B), red circles (A→B), blue triangles (B→A). (**h**, **i**) Blue arrows indicate antagonism shifts (left to right), red arrows indicate synergy shifts (right to left). (**j**) Black arrow shows score differences between sequential administrations, without directional comparison.

