## supplementary information for "Massively multiplexed microfluidics maps combinatorial and sequential antibiotic responses in 3D"

Yoon Jeong et al.

| Polymer | Natural Source | Viscosity (cP) | Reversibility | pH Range | Microfluidic Adaptability | Suitability for 3D Growth | Notes |
|---|---|---|---|---|---|---|---|
| Alginate | Brown seaweed (e.g., Laminaria, Macrocystis, Ascophyllum) | 50–200 | High (reversible , gel-sol transition with chelators) | 5–9 | High (compatible with ring-structure actuation, clogging prevented by chelators) | Excellent (supports diffusion, structural integrity) | Extracted from cell walls, rich in guluronic and mannuronic acids, used at 1% w/v for 150–300 kPa stiffness. |
| Gelatin | Animal collagen (e.g., bovine, porcine skin/bones) | 20–100 | Moderate (thermo-reversible at 30–35°C) | 5–8 | Moderate (prone to clogging without optimization) | Fair (flexible but less stable) | Derived from collagen hydrolysis, reversible at 30–35°C, used in gelation studies. |
| Agarose | Red seaweed (e.g., Gelidium, Gracilaria) | 100–500 | Low (thermo-reversible after setting) | 6–8 | Low (stable but clogging with optimization) | Excellent (robust structure, diffusion support) | Purified from agar, forms stable gels post-setting, suitable for microbial growth. |
| Chitosan | Crustacean shells (e.g., shrimp, crab) | 200–1,000 | Moderate (pH-dependent gelation) | 4–6 | Moderate (requires pH control, potential clogging) | poor (not compatible due to antimicrobial effects) | Deacetylated chitin, pH-dependent gelation, limited by antimicrobial effects. |
| Pectin | Fruit peels (e.g., citrus, apple) | 30–300 | Low (irreversible gelation) | 2.5–4 | Low (Low pH, viscosity may limit flow, potential clogging) | Poor (less compatible, Low pH) | Plant-derived, low reversibility, pH-sensitive, less compatible with rapid switching. |
| Gellan Gum | Bacterial fermentation (Sphingomonas elodea) | 50–800 | Low (thermo-reversible with cations) | 4–8 | Moderate (compatible with actuation) | Good (versatile, supports diffusion) | Microbial exopolysaccharide, thermo-reversible with cations, supports diffusion. |
| Xanthan Gum | Bacterial fermentation (Xanthomonas campestris) | 600–2,000 | Low (shear-thinning) | 3–10 | Low (high viscosity, requires flow optimization) | Fair (supports viscosity but less reversible) | High viscosity, shear-thinning, less reversible, used in flow optimization. |
| Guar Gum | Guar bean seeds (Cyamopsis tetragonoloba) | 300–1,500 | Low (irreversible gelation) | 4–9 | Low (high viscosity, potential clogging) | Poor (limited diffusion) | Plant-derived, irreversible gelation, high viscosity limits diffusion. |
| Carrageenan | Red seaweed (e.g., Chondrus crispus, Kappaphycus) | 100–400 | Moderate (thermo-reversible with cations) | 5–9 | Moderate (stable with actuation, potential clogging ) | Good (supports structure and diffusion) | Sulfated polysaccharide, thermo-reversible with cations, supports structure. |

**Supplementary Table 1. Properties of naturally-derived polysaccharide polymers[1-4]**

This table presents a comprehensive overview of the properties of naturally-derived polysaccharide polymers evaluated compatibility for the platform, designed to mimic biofilm-like matrices and support antimicrobial susceptibility testing (AST). The table includes: Viscosity (cP) at 25°C with approximate ranges based on 1–5% w/v solutions; Reversibility, assessing the ability to switch between gel and sol states, a critical factor for environmental switching in microfluidic systems to prevent channel clogging; pH Stability Range, indicating optimal pH conditions for microbial growth while maintaining the structural rigidity of the polydimethylsiloxane (PDMS) layer; Microfluidic Adaptability, evaluating compatibility with ring-structure actuation and clogging prevention; and Suitability for Biofilm-like Growth, a qualitative assessment based on support for biofilm-like growth, nutrient diffusion, and structural integrity, informed by pilot study characteristics such as viscosity and reversibility.

| Alginate Concentration (% w/v) | Calcium Ion Concentration (mM) | Stiffness (Young's Modulus, kPa) | Notes |
|---|---|---|---|
| 0.5% | 10 (Low-Moderate) | 5–10 | Lower crosslinking density than 1%, inferred from shear modulus trends. |
| 0.5% | 50 (Moderate) | 25–50 | Moderate stiffness, scaled from 1% data with rheological trends. |
| 0.5% | 100 (Moderate-High) | 50–100 | Increased crosslinking, but lower than 1% due to reduced alginate. |
| 0.5% | 200 (High) | 100–150 | Possible plateau, scaled from 1% with over-saturation effects. |
| 1% | 1.8 (Low) | 6–10 | Based on shear modulus (~2 kPa), E ≈ 3G'. |
| 1% | 10 (Low-Moderate) | 10–20 | Increased crosslinking, inferred from shear modulus trends (~72 kPa at 8 mM). |
| 1% | 50 (Moderate) | 50–100 | Enhanced stiffness with higher $Ca^{2+}$, consistent with rheological trends. |
| 1% | 100 (Moderate-High) | 100–200 | Saturated crosslinking, aligned with tensile property increases. |
| 1% | 200 (High) | 200–300 | Possible plateau due to over-saturation, inferred from high $Ca^{2+}$ effects. |
| 2% | 10 (Low-Moderate) | 20–40 | Moderate stiffness with 10 mM $CaCl_2$, consistent with literature. |
| 2% | 50 (Moderate) | 100–200 | Significant crosslinking, inferred from compressive modulus data. |
| 2% | 100 (Moderate-High) | 200–500 | Optimal range for 2%, aligning with tensile strength trends. |
| 2% | 200 (High) | 300–600 | Potential plateau or softening due to over-saturation. |

**Supplementary Table 2. Stiffness (Young's Modulus) of Alginate Hydrogels (0.5–2% w/v) with different $Ca^{2+}$ Concentration [5-9]**

| Name | Abbr. | Family | Inhibited Pathway | Classification |
|---|---|---|---|---|
| Amikacin | AMI | aminoglycoside | 30S ribosomal subunit | bactericidal |
| Cefepime | CEF | cephalosporin | Penicillin-binding proteins (cell wall) | bactericidal |
| Clindamycin | CLI | lincosamide | 50S ribosomal subunit | bacteriostatic* |
| Levofloxacin | LEV | fluoroquinolone | DNA gyrase/topoisomerase IV | Bactericidal* |
| Nitrofurantoin | NIT | nitrofuran | DNA damage/ribosomal function | bacteriostatic* |
| Sulfamethoxazole | SUL | sulfonamide | Dihydropteroate synthase (folate synthesis) | bacteriostatic |
| Rifampin | RIF | rifamycin | RNA polymerase (β-subunit) | bacteriostatic* |
| Tetracycline | TET | tetracycline | 30S ribosomal subunit | bacteriostatic |
| Trimethoprim | TRI | diaminopyrimidines | Dihydrofolate reductase (folate synthesis) | bacteriostatic |

**Supplementary Table 3. Antibiotics used in this study.**

This table details the nine antibiotics evaluated in our 3D culturomics microfluidic platform for assessing species-specific responses and optimizing combination regimens. The table includes four columns: Name and Abbreviation, listing the antibiotics as Amikacin (AMI), Cefepime (CEF), Clindamycin (CLI), Levofloxacin (LEV), Nitrofurantoin (NIT), Sulfamethoxazole (SUL), Rifampin (RIF), Tetracycline (TET), and Trimethoprim (TRI); Family, categorizing each antibiotic's chemical class; Inhibited Biochemical Pathway, specifying the targeted pathways (protein synthesis, cell wall synthesis, DNA synthesis, RNA synthesis, or folate synthesis); and Classification, indicating whether the antibiotic is bactericidal or bacteriostatic, with an asterisk (*) denoting potential concentration-dependent variations.[10]

| Antibiotics Name | *E. coli* | | *S. aureus* | | *P. aeruginosa* | | *K. pneumooniae* | |
|---|---|---|---|---|---|---|---|---|
| | **Microfluidic** | **CLSI** | **Microfluidic** | **CLSI** | **Microfluidic** | **CLSI** | **Microfluidic** | **CLSI** |
| Amikacin | 4 | ≤16 | 2 | ≤16 | 4 | ≤16 | 1 | ≤16 |
| Cefepime | 0.125 | ≤8 | 2 | ≤8 | 2 | ≤8 | 0.5 | ≤8 |
| Clindamycin | 64 | R (intrinsic) | 0.5 | ≤2 | 128 | R (intrinsic) | 32 | R (intrinsic) |
| Levofloxacin | 0.125 | ≤2 | 0.25 | ≤1 | 0.5 | ≤2 | 0.125 | ≤2 |
| Nitrofurantoin | 8 | ≤32 | 8 | ≤32 | 128 | R (intrinsic) | 4 | ≤32 |
| Sulfamethoxazole | 32 | ≤256 | 64 | ≤256 | 128 | R (intrinsic) | 32 | ≤256 |
| Rifampin | 4 | Not available | 0.5 | ≤1 | 16 | Not available | 4 | Not available |
| Tetracycline | 0.5 | ≤4 | 0.5 | ≤4 | 8 | ≤8 | 0.5 | ≤4 |
| Trimethoprim | 0.25 | ≤8 | 4 | ≤8 | 64 | R (intrinsic) | 1 | ≤8 |

**Supplementary Table 4. MIC comparison between microfluidic platform and CLSI[11, 12]**

This table compares the minimum inhibitory concentrations (MICs) of nine antibiotics—Amikacin (AMI), Cefepime (CEF), Clindamycin (CLI), Levofloxacin (LEV), Nitrofurantoin (NIT), Sulfamethoxazole (SUL), Rifampin (RIF), Tetracycline (TET), and Trimethoprim (TRI)—against four bacterial species (*Escherichia coli*, *Staphylococcus aureus*, *Pseudomonas aeruginosa*, and *Klebsiella pneumoniae*) obtained from microfluidic platform and CLSI. The table comprises: Name and Abbreviation of the antibiotic; Microfluidic MIC (μg/ml) values for *E. coli*, *S. aureus*, *P. aeruginosa*, and *K. pneumoniae*, respectively, assessed within the 256–0.5 μg/ml range; and CLSI MIC Range (μg/ml) for susceptible strains, where, "R (intrinsic)" for intrinsic resistance, and "Not available" for Rifampin against Gram-negatives due to limited clinical relevance. CLSI (Clinical and Laboratory Standards Institute)

| | Stiffness (Young’s Modulus, kPa) | Alginate Concentration (% w/v) | Calcium Ion Concentration (mM) | Notes |
|---|---|---|---|---|
| Gelation 1 | 300 | 1% | 180 (High) | Approaching saturation plateau, inferred from high $Ca^{2+}$ effects (200–300 kPa range). |
| Gelation 2 | 250 | 1% | 140 (Moderate-High) | Further increase from 110 mM, consistent with saturated crosslinking trends. |
| Gelation 3 | 200 | 1% | 110 (Moderate-High) | Adjusted upward from 100 mM (100–200 kPa), reflecting increased crosslinking. |
| Gelation 4 | 150 | 1% | 80 (Moderate) | Interpolated between 50–100 mM range (50–100 kPa and 100–200 kPa), aligning with tensile property trends. |

**Supplementary Table 5. Alginate Hydrogels (1% w/v) with Calcium Ion Concentration (used in this study, for the stiffness range ~150, 200, 250, 300 kPa, respectively) [9]**

| Pair # | Pair (Abbr.) | Inhibited Pathways | Classifications | A+B | A→B | B→A | Score Shift (>0.3) | Notes |
|---|---|---|---|---|---|---|---|---|
| 1 | AMI-CEF | 30S ribosomal subunit–Penicillin-binding proteins | Bactericidal-Bactericidal | Additive | Additive | Additive | No | Effective for E. coli and other Gram-negatives |
| 2 | AMI-CLI | 30S ribosomal subunit–50S ribosomal subunit | Bactericidal-Bacteriostatic* | Synergy | Additive | Synergy | No | CLI resistance in E. coli (Gram-negative) |
| 3 | AMI-LEV | 30S ribosomal subunit–DNA gyrase/topoisomerase IV | Bactericidal-Bactericidal | Additive | Additive | Additive | No | Broad Gram-negative coverage, including E. coli |
| 4 | AMI-NIT | 30S ribosomal subunit–DNA damage/ribosomal function | Bactericidal-Bacteriostatic* | Synergy | Synergy | Synergy | No | 30S overlap may enhance synergy in E. coli |
| 5 | AMI-SUL | 30S ribosomal subunit–Dihydropteroate synthase | Bactericidal-Bacteriostatic | Additive | Additive | Antagonism | Yes | SUL→AMI reduces efficacy in E. coli |
| 6 | AMI-RIF | 30S ribosomal subunit–RNA polymerase | Bactericidal-Bacteriostatic* | Additive | Additive | Additive | No | RIF efficacy limited in E. coli (Gram-negative) |
| 7 | AMI-TET | 30S ribosomal subunit–30S ribosomal subunit | Bactericidal-Bacteriostatic | Synergy | Additive | Additive | No | 30S overlap; TET→AMI reduces efficacy in E. coli |
| 8 | AMI-TRI | 30S ribosomal subunit–Dihydrofolate reductase | Bactericidal-Bacteriostatic | Synergy | Synergy | Synergy | No | TRI→AMI reduces efficacy in E. coli |
| 9 | CEF-CLI | Penicillin-binding proteins–50S ribosomal subunit | Bactericidal-Bacteriostatic* | Additive | Additive | Additive | No | Well-known antagonism; CLI resistance in E. coli (Gram-negative) |
| 10 | CEF-LEV | Penicillin-binding proteins–DNA gyrase/topoisomerase IV | Bactericidal-Bactericidal | Additive | Additive | Synergy | Yes | Effective for E. coli; LEV→CEF enhances efficacy |
| 11 | CEF-NIT | Penicillin-binding proteins–DNA damage/ribosomal function | Bactericidal-Bacteriostatic* | Antagonism | Synergy* | Additive | Yes | NIT bactericidal at high conc.; Bactericidal effects overlap in E. coli |
| 12 | CEF-SUL | Penicillin-binding proteins–Dihydropteroate synthase | Bactericidal-Bacteriostatic | Additive | Additive | Antagonism | Yes | Bacteriostatic → Bactericidal reduces efficacy in E. coli. |
| 13 | CEF-RIF | Penicillin-binding proteins–RNA polymerase | Bactericidal-Bacteriostatic* | Additive | Additive | Additive | No | RIF bactericidal at high conc.; Limited efficacy in E. coli |
| 14 | CEF-TET | Penicillin-binding proteins–30S ribosomal subunit | Bactericidal-Bacteriostatic | Additive | Additive | Antagonism | Yes | Bacteriostatic → Bactericidal reduces efficacy in E. coli. |
| 15 | CEF-TRI | Penicillin-binding proteins–Dihydrofolate reductase | Bactericidal-Bacteriostatic | Additive | Additive | Additive | Yes | — |
| 16 | CLI-LEV | 50S ribosomal subunit–DNA gyrase/topoisomerase IV | Bacteriostatic*-Bactericidal | Synergy | Synergy | Additive | No | CLI resistance in E. coli (Gram-negative) |
| 17 | CLI-NIT | 50S ribosomal subunit–DNA damage/ribosomal function | Bacteriostatic*-Bacteriostatic* | Additive | Additive | Additive | No | CLI resistance in E. coli; Different protein synthesis targets |
| 18 | CLI-SUL | 50S ribosomal subunit–Dihydropteroate synthase | Bacteriostatic*-Bacteriostatic | Additive | Additive | Additive | No | — |
| 19 | CLI-RIF | 50S ribosomal subunit–RNA polymerase | Bacteriostatic*-Bacteriostatic* | Additive | Synergy | Additive | Yes | CLI→RIF enhances efficacy; |
| 20 | CLI-TET | 50S ribosomal subunit–30S ribosomal subunit | Bacteriostatic*-Bacteriostatic | Antagonism | Additive | Additive | No | CLI resistance in E. coli; Different ribosomal targets |
| 21 | CLI-TRI | 50S ribosomal subunit–Dihydrofolate reductase | Bacteriostatic*-Bacteriostatic | Synergy | Synergy | Additive | No | — |

| | | | | | | | | |
|---|---|---|---|---|---|---|---|---|
| 22 | LEV-NIT | DNA gyrase/topoisomerase IV–DNA damage/ribosomal function | Bactericidal-Bacteriostatic* | Antagonism | Antagonism | Antagonism | No | NIT bactericidal at high conc. in E. coli |
| 23 | LEV-SUL | DNA gyrase/topoisomerase IV–Dihydropteroate synthase | Bactericidal-Bacteriostatic | Additive | Additive | Antagonism | Yes | Well-known antagonism; SUL→LEV reduces efficacy in E. coli |
| 24 | LEV-RIF | DNA gyrase/topoisomerase IV–RNA polymerase | Bactericidal-Bacteriostatic* | Additive | Additive | Antagonism | Yes | RIF bactericidal at high conc.; Limited efficacy in E. coli |
| 25 | LEV-TET | DNA gyrase/topoisomerase IV–30S ribosomal subunit | Bactericidal-Bacteriostatic | Antagonism | Antagonism | Antagonism | No | TET→LEV reduces efficacy in E. coli |
| 26 | LEV-TRI | DNA gyrase/topoisomerase IV–Dihydrofolate reductase | Bactericidal-Bacteriostatic | Additive | Synergy | Additive | Yes | TRI→LEV reduces efficacy in E. coli |
| 27 | NIT-SUL | DNA damage/ribosomal function–Dihydropteroate synthase | Bacteriostatic*-Bacteriostatic | Antagonism | Additive | Additive | No | — |
| 28 | NIT-RIF | DNA damage/ribosomal function–RNA polymerase | Bacteriostatic*-Bacteriostatic* | Antagonism | Additive | Antagonism | No | RIF limited efficacy in E. coli |
| 29 | NIT-TET | DNA damage/ribosomal function–30S ribosomal subunit | Bacteriostatic*-Bacteriostatic | Additive | Additive | Additive | No | Different protein synthesis targets in E. coli |
| 30 | NIT-TRI | DNA damage/ribosomal function–Dihydrofolate reductase | Bacteriostatic*-Bacteriostatic | Synergy | Synergy | Synergy | No | — |
| 31 | SUL-RIF | Dihydropteroate synthase–RNA polymerase | Bacteriostatic-Bacteriostatic* | Antagonism | Additive | Antagonism | Yes | — |
| 32 | SUL-TET | Dihydropteroate synthase–30S ribosomal subunit | Bacteriostatic-Bacteriostatic | Antagonism | Antagonism | Additive | Yes | Bacteriostatic effects overlap in E. coli |
| 33 | SUL-TRI | Dihydropteroate synthase–Dihydrofolate reductase | Bacteriostatic-Bacteriostatic | Synergy | Synergy | Synergy | No | Well-known synergy (co-trimoxazole); SUL→TRI disrupts folate pool in E. coli |
| 34 | RIF-TET | RNA polymerase–30S ribosomal subunit | Bacteriostatic*-Bacteriostatic | Antagonism | Antagonism | Additive | Yes | RIF limited efficacy in E. coli |
| 35 | RIF-TRI | RNA polymerase–Dihydrofolate reductase | Bacteriostatic*-Bacteriostatic | Additive | Additive | Additive | No | — |
| 36 | TET-TRI | 30S ribosomal subunit–Dihydrofolate reductase | Bacteriostatic-Bacteriostatic | Synergy | Synergy | Synergy | No | — |

**Supplementary Table 6. 36 antibiotic combination pairs**

This table details 36 antibiotic combination pairs derived from nine antibiotics. The table comprises six columns: Numbers for each antibiotic pair. Abbreviation: Inhibited Biochemical Pathway: Classification: Antibiotic classification as bactericidal or bacteriostatic. Simultaneous (A+B) and sequential (A→B, B→A) dosing regimens. Score shift: Indicates pairs with synergy score shifts across dosing regimens (Yes/No), with 12 pairs showing sequence-dependent changes in this study. Score shift >0.3 (on a scale 0 to 1) in 12 of 36 antibiotic pairs, observed in either A→B or B→A sequential dosing. Notes: Provide resistance patterns (e.g., CLI resistance in Gram-negatives), mechanistic insights (e.g., 30S overlap, folate pool disruption), or pharmacokinetic effects (e.g., SUL metabolism by CYP2C9). with an asterisk (*) indicating potential concentration-dependent variations.

**Supplementary Video 1:** Medium Loading, Hydrogel Structural Stability and Diffusion. This video demonstrates fluidic environment switching within microfluidic chambers, controlled by valve open/close cycles with a latency of approximately 0.5 seconds. Hydrogel stability is shown through multiple valve actuation cycles. Also shown is diffusion-based reagent delivery to the hydrogel structures (left).

**Supplementary Video 2:** Bacterial Growth (Free vs. Hydrogel). This video compares the growth dynamics of E. coli in free solution versus in a 1% (w/v) alginate hydrogel matrix within the chambers, showing morphological differences.

**Supplementary Video 3:** gfp E. coli Growth Monitoring. This video shows gfp E. coli bacterial growth for 5 hours after growth arrest, measured via automated microscopy.

**Supplementary Video 4:** Input Multiplexer (Multicolor Control for n-Fold Dilution). This video illustrates the fluidic control of the combinatorial multiplexing system, enabling n-fold dilution and reagent combination. Multicolor food dyes (red, green, blue, and yellow) connected to 1–9 nodes visualize the delivery of nine reagents via a custom graphical user interface (GUI), as described in Extended Data Fig. 4. All combination of the inputs can be generated automatically and be flown to the cells cultured in individual chambers.

**Supplementary Video 5:** Independently controllable 512-Chamber device with hydrogel islands, and Input Control for Environmental Switching. This video shows the 512-chamber array in operation, integrating input multiplexing and chamber control across a 32 × 16 grid. Ring-shaped valves regulate flow, demonstrating the chip's ability to test chambers under diverse conditions. Each hydrogel-island chamber can be loaded with different reagents without influencing the others, and the reagents can be changed over time. The separated video clips are not synchronized for time control.

### Supplementary Note 1. Bacterial Colony Culture in 3D Hydrogel

The use of a hydrogel matrix is a key feature of our platform, as it replicates biofilm-like conditions, which are more representative of bacterial behavior in natural or clinical settings compared to traditional platforms in liquid culture, or 1D or 2D setting. This note describes the preparation of the 3D hydrogel environment within the ultra-multiplexed chip's chambers, the process of bacterial culturing, and the control of population density.

#### Hydrogel Fabrication and Chamber Isolation

The ultra-multiplexed platform uses alginate-based hydrogels to create 3D cultivation environments within the 512 chambers. Alginate solutions, prepared at concentrations of 1–2% (w/v), are crosslinked with calcium ions ($Ca^{2+}$, 0.01–0.1 M) to form a stable solid matrix that encages bacterial colonies. This 3D environment mimics the extracellular matrix of biofilms, where bacteria are embedded in a protective matrix, altering their growth dynamics and antibiotic susceptibility.[13] Bacterial inoculums, adjusted to an optical density (OD600) of 0.5 to ensure cells are in the exponential growth phase (avoiding stationary phase), is mixed with the alginate solution and loaded into the chambers via the outlet ports

The high viscosity of the 1–2% alginate solution necessitates a controlled gelation process to prevent channel clogging and ensure uniform bacterial encapsulation in chambers. After loading the bacterial-alginate mixture, a 0.1 M calcium chloride ($CaCl_2$) solution is introduced into the outer chamber region through dedicated channels. Ring-shaped valves are actuated altogether for 2–3 seconds to allow the calcium ions to diffuse into the inner space, triggering rapid gelation of the alginate. This forms a stable hydrogel structure with tunable stiffness, which entraps bacterial cells within the central compartments, preventing cross-contamination between chambers. Once the ring valves are closed, the outer culture region can be filled with culture media (LB or MH2) or antibiotics using the same valve system. This precise spatial confinement, validated through dye diffusion imaging, ensures that each chamber operates as an independent microenvironment, supporting high-throughput testing of bacterial responses to various conditions.

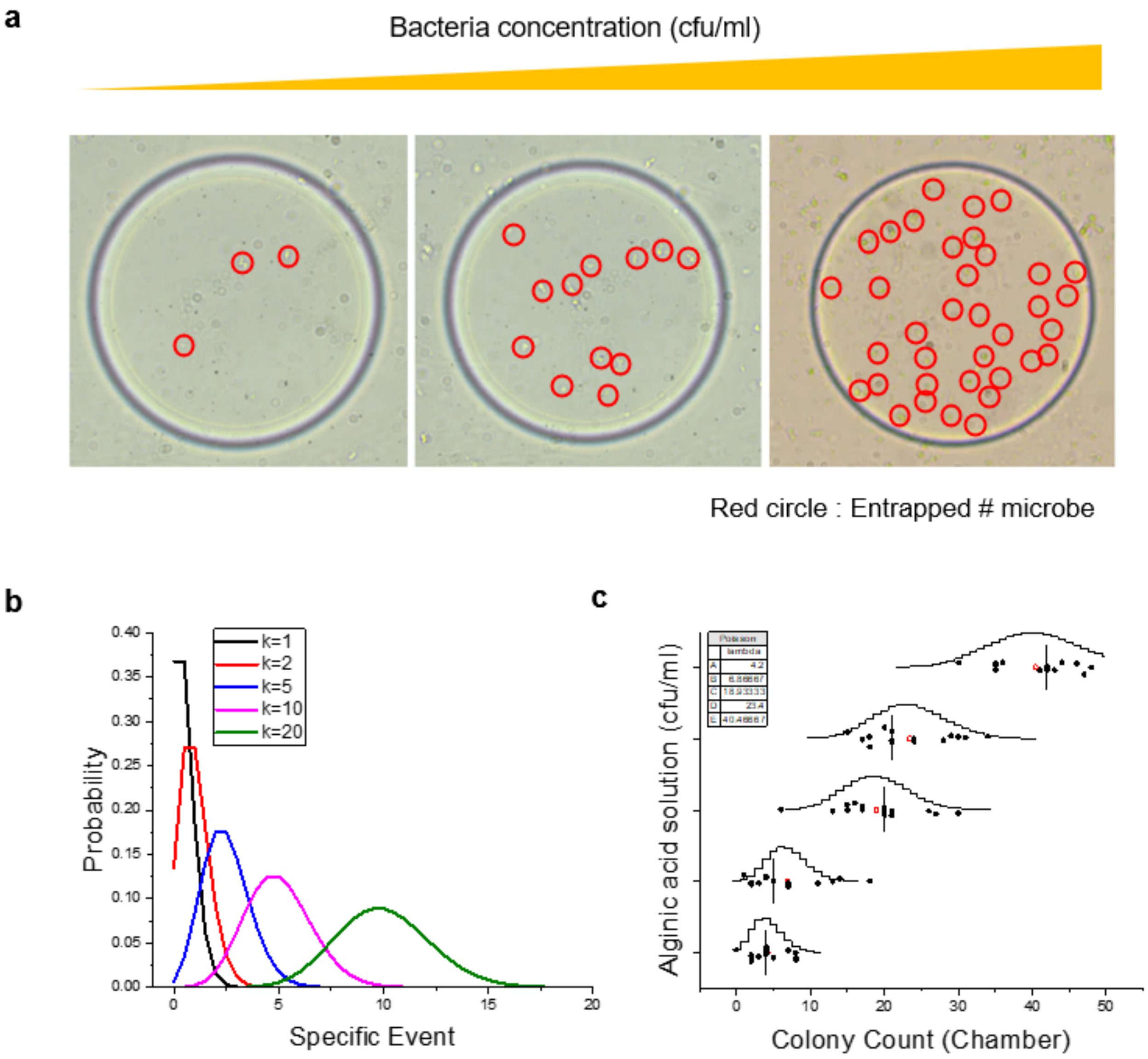


**Supplementary Fig 1.** (a) microscopy images of culture chamber with bacterial entrapment (red circles) at varying concentrations (cfu/mL) within a 2% (w/v) alginate solution. (b) Poisson distribution plot and parameters (λ=1, λ=10, λ=20) (c) Scatter plot of colony counts per chamber across alginate solution concentrations, overlaid with the Poisson distribution model to illustrate probability density of colony counts per chamber. cfu: colony forming unit.

The Poisson distribution model is used to control bacterial population density within the chambers (**Supplementary Fig 1)**

Defined by the probability function,

$$P(X = k) = \frac{e^{-\lambda}\ \lambda^k}{k!}$$

Where, random variable $X$, the number of occurrences $k$, Euler's number $e$. The model optimizes bacterial loading to achieve approximately 25–30 colonies per chamber.[14] The parameters λ=1, λ=10, and λ=20 correspond to sparse (0–10 colonies, ~$10^4$ CFU/mL) to dense (up to 50 colonies, ~$10^6$ CFU/mL) distributions. This controlled loading process ensures colony uniformity across the 512 chambers, critical for high-throughput testing. The scatter plot of observed colony counts, overlaid with the Poisson model, validates the effectiveness of this approach in achieving consistent population densities.

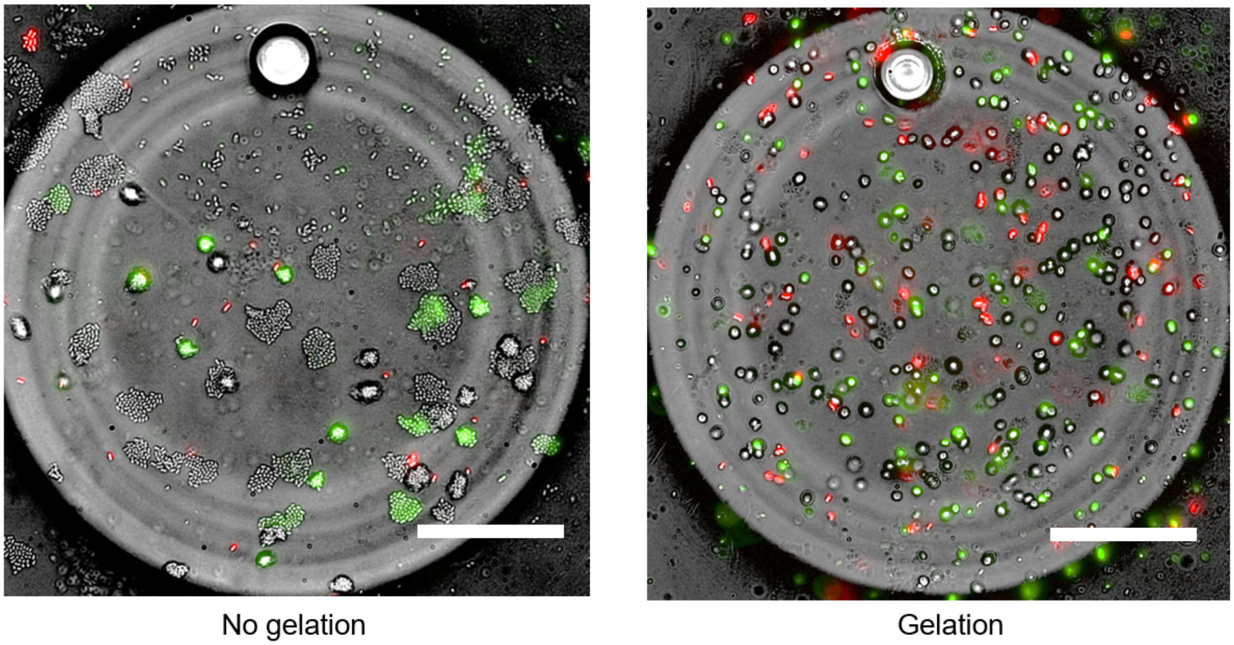


**Supplementary Fig 2**. Microscopic images for *E. coli* in a culture chamber (a) alginate polymer (1 % w/v) only (b) alginate hydrogel (1 % w/v) crosslinked by 0.1 M $CaCl_2$ solution, after 6 hours of incubation; Scale bar, 50 μm.

**Bacteria Immobilization and Colony Culture**

The immobilization of bacteria within the alginate hydrogel is intended for studying colony dynamics in a controlled, biofilm-like environment. Unlike planktonic (free-swimming) bacteria, which disperse rapidly and complicate quantitative analysis, hydrogel-encapsulated bacteria form discrete colonies that can be tracked over time (**Supplementary Fig 2).** After 6 hours of incubation, hydrogel-encapsulated cultures exhibit distinct colony formation, enabling precise quantification of growth and antibiotic effects. The hydrogel matrix restricts bacterial motility, mimicking the sessile lifestyle of biofilms, which is critical for studying antibiotic resistance mechanisms that differ from those in planktonic cultures.[15]

One of the important capabilities of our platform is its ability to execute environmental switching without causing any disruption to the cell cultures embedded within the hydrogel, as illustrated in Fig 1g as well as the detailed process in Extended Data Fig 1e. The workflow begins with the initial establishment of microcolonies with hydrogel stiffness (~150 kPa) within a microfluidic chamber, where cells are allowed to grow and form small colonies. This is followed by the precise introduction of growth media containing essential nutrients (or antibiotics) through microfluidic channels and valve actuation which ensures that the cultures receive the necessary sustenance and experimental treatments. To maintain control over the environment, the system then closes its valves, isolating the microcolonies and stabilizing the conditions within the chamber. Subsequently, the inner valves are opened selectively for media exchange or observation without disturbing the microbial cultures in hydrogels. The process includes washing steps at the outer chamber region to clear out wastes and turn into additional cultivation phases to support ongoing growth. This entire cycle can be repeated multiple times to mimic the dynamic environmental changes that cells might encounter in nature. This repetitive switching is further detailed through two distinct cycles (Extended Data Fig 1f): Each of these transitions is carefully designed to reflect changes in the microcolony's morphology and viability, offering valuable insights into how these communities respond to fluctuating conditions.

**Supplementary Note 2. Hydrogel diffusion Release Kinetics and Modeling**

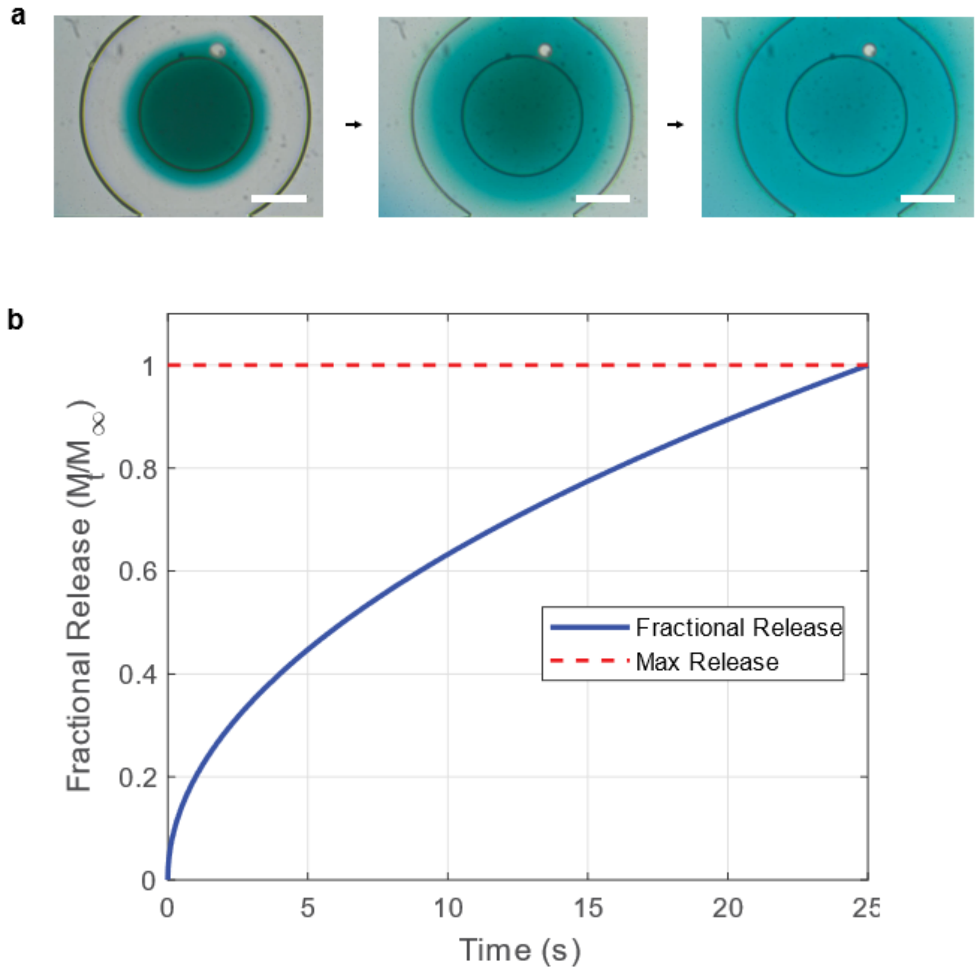


**Supplementary Fig 3**. **a**. Microscopic images of diffusion release from a hydrogel structure, depicting the progression of a circular structure (hydrogel environment with cyan blue dye) over time: stages of a release process by opening encapsulation valve: Scale bar, 100 μm. **b.** Simulation graph of fractional release ($M/M_0$) versus time (s); Practional Release (blue curve), Maximum Release (red dashed line).

We calculated the diffusion release kinetics of a hydrogel, modeled as a finite cylinder (200 μm diameter, 30 μm height, volume ~0.942 nL), into a surrounding aqueous medium (DI water). The simulation examines the release of a diffusant (small dye molecules) under Fickian diffusion with perfect sink conditions, where the surrounding medium concentration remains negligible. The hydrogel's small dimensions enable rapid equilibrium within 25 seconds, a microscale diffusion process influenced by geometry and material properties[16].

The simulation employs an early-time approximation for fractional release, given by the equation.

$$\frac{M_t}{M_\infty} \approx \frac{4}{\pi}\sqrt{\frac{Dt}{l^2}}$$

Where, $l = 30\ \mu m$ is the hydrogel height, $D$ is the diffusion coefficient, $t$ is time, and $\frac{M_t}{M_\infty}$ is the fraction of diffusant released relative to the total releasable amount. This approximation assumes diffusion is primarily governed by the axial direction due to the cylinder's small height. To determine $D$, The slope of this plot is given by:

$$Slope = \frac{4}{l}\sqrt{\frac{D}{\pi}}$$

Rearrange to solve for $D$:

$$D = \frac{Slope^2 \cdot \pi l^2}{16}$$

Fit a straight line to the early data points to extract the slope via dye fading intensity in the image (**Supplementary Fig 3a)**.

For small molecules like water or dyes in hydrogels, we computed the diffusion coefficient. Based on an equilibrium time of 25 seconds, $D$ was estimated at $2.83 \times 10^{-8}\ cm^2/s$ , consistent with small-molecule diffusion in hydrogels, though potentially affected by porosity and swelling. Visualization of the release kinetics was achieved through a MATLAB-based simulation, generating a plot of fractional release versus time (**Supplementary Fig 3b)**

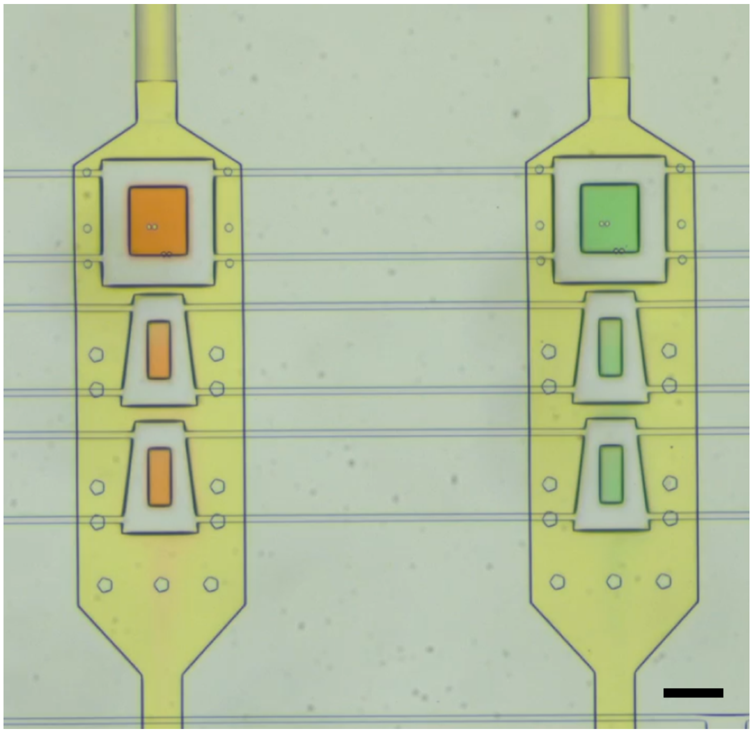

**Supplementary Figure 4.** Microscopy image of the microfluidic chambers featuring three separate hydrogel islands, controlled by with three encapsulation valves. Two individual chambers are shown. Each chamber includes a series of interconnected channels and compartments, allowing for the placement and individual control of hydrogel islands. A different cell type can be loaded and cultured in each different island. This setup enables the creation of complex culture geometries, such as co-culture islands, within a single microfluidic environment; Scale bar, 200 μm.

### Supplementary Note 3. Colony Tracking Imaging Data Processing and Validation

The platform employs automated fluorescence microscopy to capture high-resolution z-stack images, consisting of 25 slices with 2 μm spacing, covering a 50 μm depth. These images are acquired every 5 minutes to monitor bacterial growth dynamics within the hydrogel environment. The MATLAB-based GUI synchronizes valve actuations, imaging parameters, and environmental controls, maintaining a stable culture temperature of 37°C ± 0.1°C.

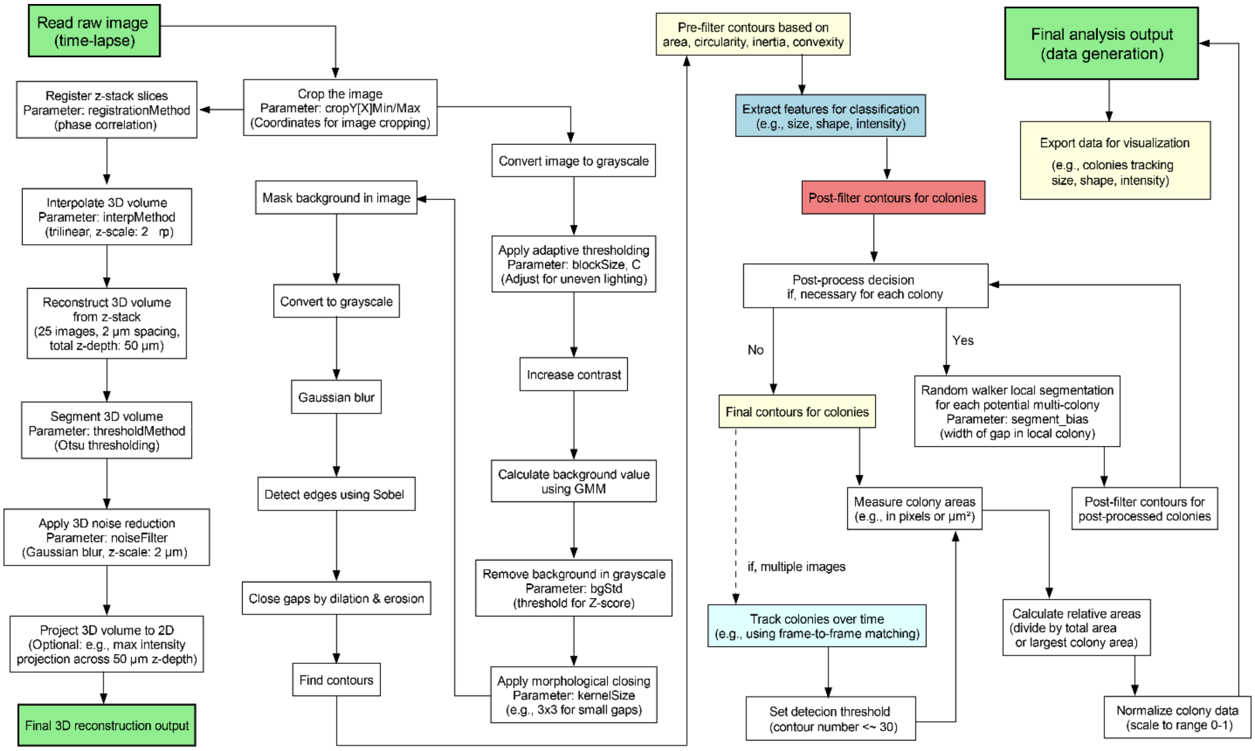


**Supplementary Fig 5**. Flowchart of the image processing pipeline for analyzing time-lapse images. The process initiates with reading a raw time-lapse image, followed by z-stack slice registration, cropping, and grayscale conversion. Key steps include adaptive thresholding, Gaussian blur, and contour detection for feature extraction (e.g., size, shape, intensity) and colony classification. The pipeline incorporates 3D volume reconstruction, background removal, and post-processing, with adjustable parameters (e.g., blockSize, threshold methods, kernelSize). This pipeline supports the phenotypic categorization (e.g., growth, colony formation, phenotypic changes) of up to 15,360 colonies in the microfluidic platform and provides segmented regions for 2D image-based quantification (normalized and scaled to a 0-1 range)

The data processing pipeline includes multiple stages (**Supplementary Fig 5**): image preprocessing, colony segmentation, and quantitative analysis of growth metrics (e.g., colony volume, fluorescence intensity).[17] The process starts with reading a raw time-lapse image, followed by registering z-stack slices using phase correlation (parameter: registrationMethod) and interpolating the 3D volume (parameter: interpMethod, trilinear, z-scale: 2 µm). The image is cropped based on specified coordinates (parameter: cropYX[minMax]) and converted to grayscale. Adaptive thresholding (parameter: blockSize, C) adjusts for uneven lighting, followed by Gaussian blur and contrast enhancement. Background value is calculated using GMM, and the background is removed in grayscale (parameter: bgStd, threshold of Z-score). Contours are found, with gaps closed by dilation and erosion, and morphological closing applied (parameter: kernelSize). Pre-filtering of contours is based on area, circularity, and inertia, leading to feature extraction for classification (e.g., size, shape, intensity). Post-processing decisions determine if further processing such as random walker local segmentation, parameter: gap in local colony bias is needed, followed by measuring areas in pixels or $\mu m^2$ and tracking colonies over time using frame-to-frame matching. For multiple images, a detection threshold (contour number < 50) is set. The pipeline includes 3D reconstruction from z-stack images (2 µm spacing, total depth: 50 µm) with optional 3D noise reduction (parameter: noiseFilter, Gaussian blur, z-scale: 2 µm) and projection to 2D (e.g., max intensity across 50 µm z-depth). The final outputs are a 3D reconstruction and an analysis dataset, exported for visualization (e.g., colony size, shape, intensity tracking), with normalized colony data scaled to a 0-1 range.

**Colony Detection Error and Challenges**

The tracking pipeline enables the identification of antibiotic effects, including synergistic or antagonistic interactions among 36 antibiotic pairs tested across the 512 chambers. However, a significant challenge in image analysis arises from variations in hydrogel stiffness (10 to 300 kPa), which affect bacterial colony morphology and imaging characteristics (**Supplementary Fig 6a**). At higher stiffness (e.g., 300 kPa), colonies appear more compact, while at lower stiffness (e.g., 50 kPa), they spread more diffusely, complicating automated colony detection and segmentation.

Our custom image analysis algorithms struggle to uniformly process images across this range of stiffness conditions, as morphological differences impact contouring and volume estimation (**Supplementary Fig 6b,c**). To address this, the platform adopts a single gelation condition (optimized alginate concentration and crosslinking parameters) for high-throughput screening, ensuring consistency in colony morphology and enabling reliable quantification of antibiotic effects across all 512 chambers.

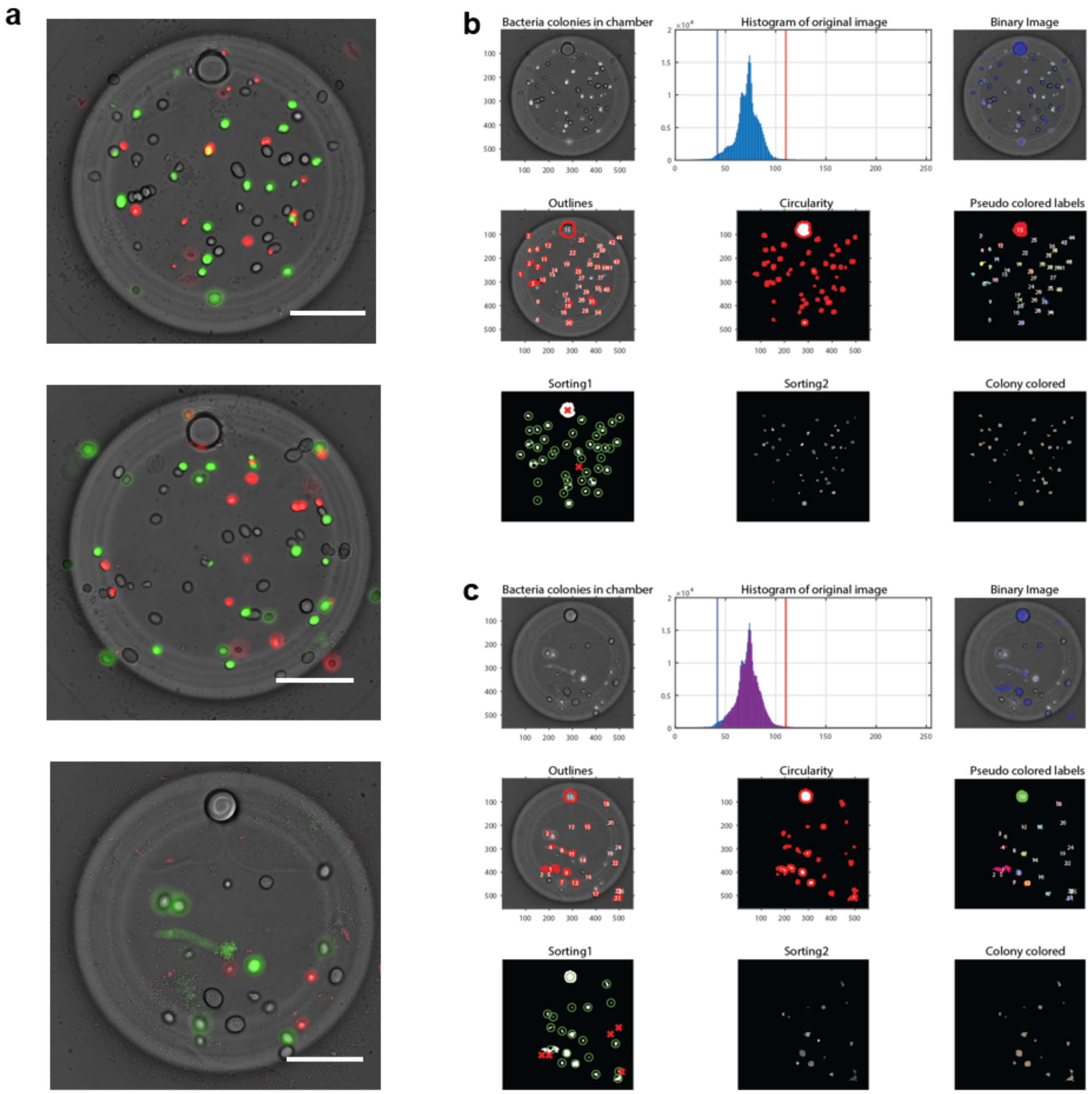


**Supplementary Fig 6**. (a) microscopic images (stiffness, top : 300 kPa, middle : 150 kPa, bottom : 50 kPa). Scale bar, 50 μm. (b-c) Comparison of image processing results for microbial colony detection, providing a visual representation of how raw images are processed to isolate and segment colonies. (b) Stiffness at 300 kPa (c) Stiffness at 50 kPa.

### Supplementary Note 4. Large-scale Integration Platform and Automation Setup

#### Overview of Ultra-multiplexed Experimental Design

The ultra-multiplexed platform facilitates 3D bacterial culturomics not only for enhanced scalability, controllability but also its advanced applicability. Our platform incorporates three multiplexers for its distinct functionality including fluid trafficking and retention in each chamber (**Supplementary Fig 7**). The platform's design—integrating compartmentalized chambers as a functional culture unit to use soft hydrogel and a radial input system with 18 inlets and 9 nodes—supports microbial 3D culturomics, to deliver uniform multi-reagent mixing across 512 chambers with <1% variability for targeted concentrations. To demonstrate its applicability in 3D hydrogel environments, we show advanced AST experiments for evaluation of synergy score shift when co-administered drug combinations. A total of 25 conditions per drug pair across 36 pairs and 3 dosing regimens (A+B, A $\rightarrow$ B, and B $\rightarrow$ A) results in 2,700 conditions tested using an analysis pipeline for 3D colony tracking algorithms.

The platform' system is designed to handle large-scale, parallelized experiments, enabling researchers to screen hundreds of experimental conditions efficiently. The platform's core component is a 512-chamber microfluidic chip, where each chamber has a volume of approximately 1 nL and features inner compartmentalization, including unique ring-shaped structure (200 μm diameter × 30 μm height) to support bacterial encapsulation in hydrogel. These chambers control the growth of approximately 50 bacterial colonies (up to 15,360 colonies across the chip, 30 colonies per chamber) encapsulated within 3D hydrogel matrices, which mimic the complex, biofilm-like environments found in natural bacterial habitats.[18] This 3D setup enhances the physiological relevance of antibiotic testing by replicating conditions where bacteria exhibit altered resistance profiles compared to traditional planktonic cultures. The chip supports up to hundreds of programmable antibiotic regimens, including 36 antibiotic pairs with simultaneous and sequential dosing, enabling comprehensive combinatorial screening. This note details the platform's architecture, fabrication process, automation systems, and data acquisition workflows, with references to supplementary materials for additional context.

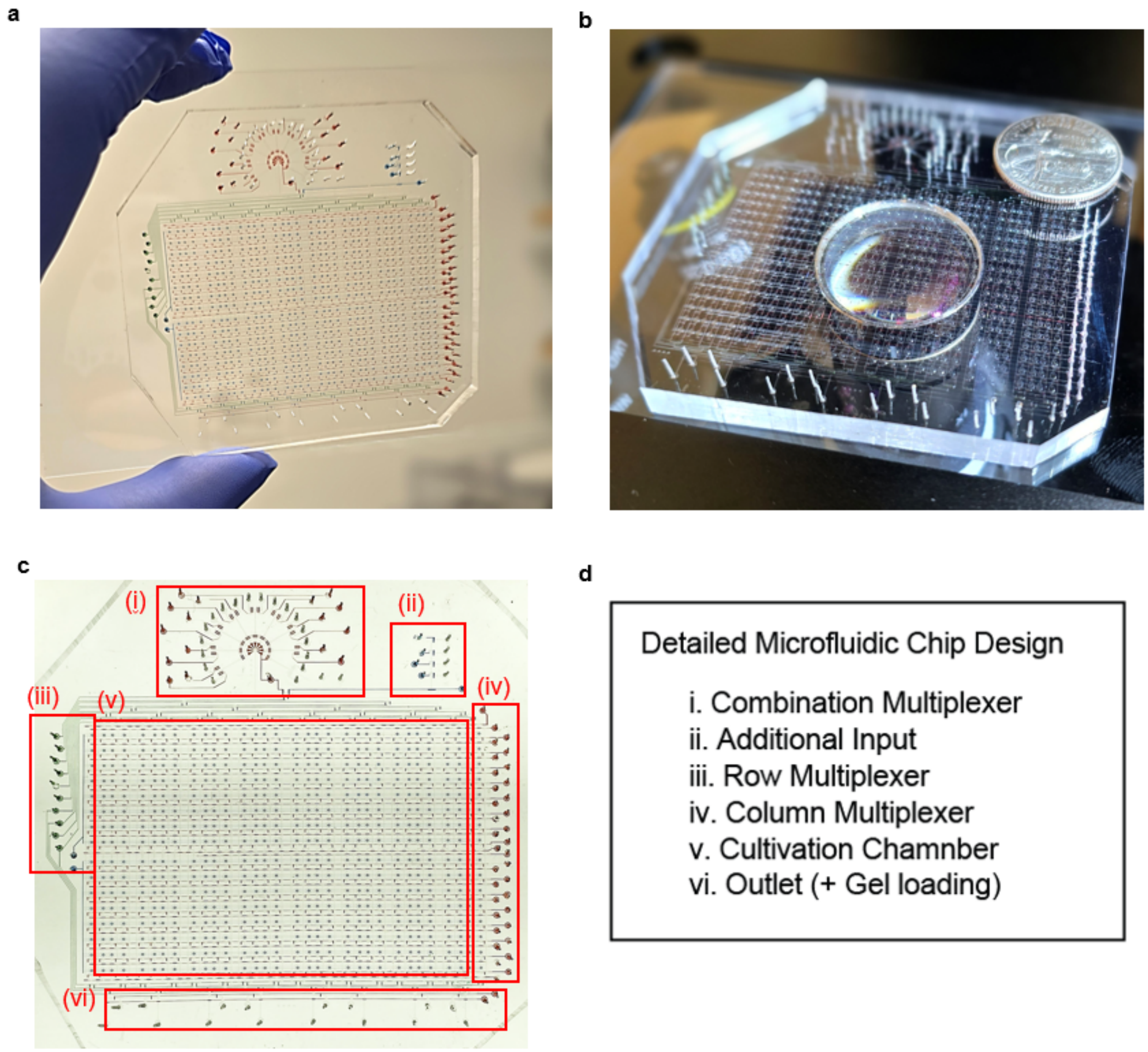


**Supplementary Fig 7.** (a) Image of the fabricated ultra-multiplexed chip (8 x 8 $cm^2$) (b) Image of the chip on a microscope stage with a U.S. quarter (diameter ~24 mm) for scale (c-d) Image of the chip and its architectural components and fluidic control systems.

### Scalability and Automation

The ultra-multiplexed platform achieves exceptional scalability by operating multiple 512-chamber chips simultaneously, allowing us to test up to thousands of unique experimental conditions across multiple chips. This is facilitated by sequential dosing regimens, where reagents such as antibiotics, media, or buffers are delivered in a controlled, time-

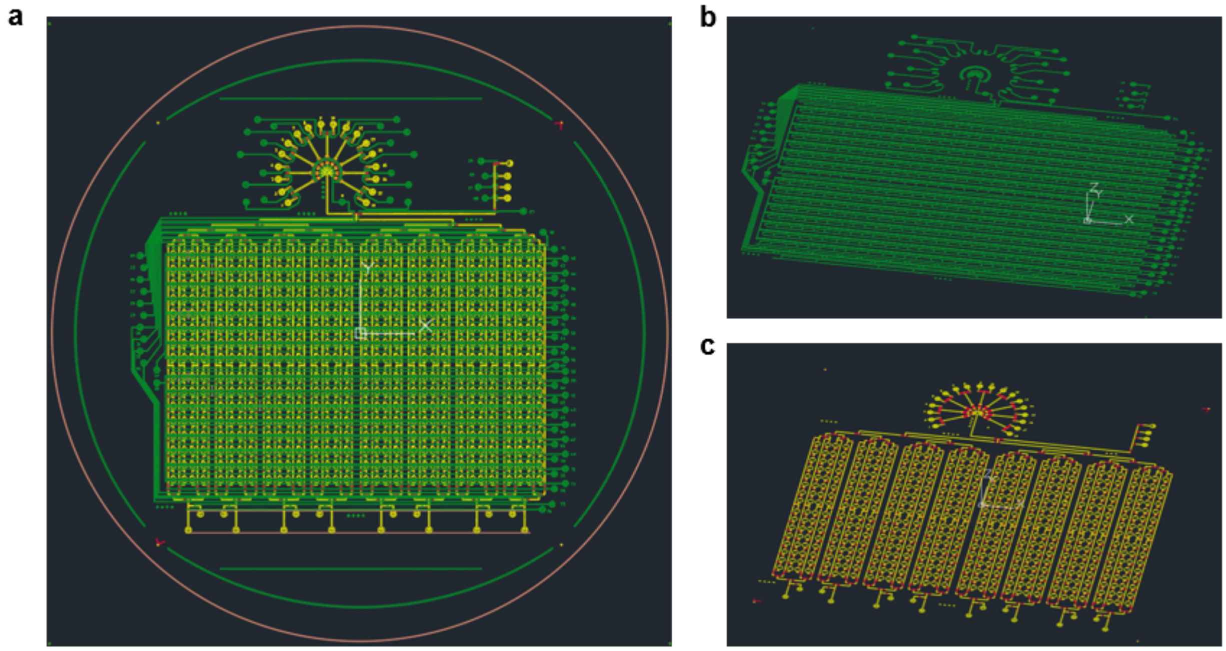


**Supplementary Fig 8.** (a) AutoCAD schematic (left) and (b-c) 3D isometric view of the ultra-multiplexed chip designed for fabrication on a 4-inch silicon wafer: (b) Control layer (green, SU-8), (c) flow layer (red, AZ-40; yellow, SU-8). The chip comprises three layers: a positive photoresist fluid layer (red line, Fig. S2), a negative photoresist fluid layer (green line, Fig. S2), and a negative photoresist control layer (blue line, Fig. S2). Fluid channels for microbial cell loading and media flow are ~150 μm wide, while hydrogel chambers, designed for compartmentalization and media retention, exchange and diffusion, are a volume of approximately 1–2 nL, ~50 bacterial colonies per chamber. Each chamber (200 μm diameter × 30 μm, ~1 nL volume) are connected with a washing channel.

dependent manner to evaluate dynamic bacterial responses. The automation system is a critical feature, ensuring high reproducibility and minimizing manual intervention, which reduces human error and variability in experimental outcomes.[19] Automation is driven by a MATLAB-based graphical user interface (GUI) that integrates with Excel files (attached Supplementary file "Combinatorial Inputs.xlsx") to program the operation of 76 control valve lines in the chip, actuated by external solenoid valves. The GUI import the Excel file to execute a pneumatic system operating at pressures between 10 and 35 psi.

This system controls Quake-style valves[20], which are microfabricated, pneumatically actuated valves that precisely regulate fluid flow within the chip. Reagents are delivered at different flow rates, ensuring accurate dosing across the 512 chambers. The precision of this fluidic control is essential for maintaining consistent experimental conditions, particularly when testing complex antibiotic combinations or sequential treatments. Furthermore, data acquisition is fully automated using fluorescence microscopy (Nikon Ti-Eclipse) to capture time-lapse z-stack images of each chamber (~1 second for 512 chambers, every 7-8 minutes per cycle) over experimental durations of up to 72 hours. These z-stack images provide three-dimensional data, capturing the spatial dynamics of bacterial colonies within the hydrogel matrix. A custom MATLAB-based image processing pipeline analyzes these images, tracking colony growth and morphology to quantify bacterial responses to antibiotics. This pipeline enables high-throughput analysis of colony dynamics across all 512 chambers, ensuring robust and reproducible data collection. By automating fluidic control, microbial cell culture and data acquisition, the platform minimizes manual handling, enhances experimental throughput, and ensures consistency across large-scale experiments.

**System Architecture**

The ultra-multiplexed chip is fabricated using advanced microfabrication techniques, specifically soft lithography, multi-layer PDMS alignment, and plasma bonding, to achieve micron-scale precision in channel and chamber geometries.[21] The fabrication process begins with the creation of master molds using conventional soft lithography. For the fluid layer, negative photoresist (MicroChemicals, SU-8 3025) is spin-coated at 3,200 RPM to achieve ~30–35 µm height features for channels and chambers, while positive photoresist (MicroChemicals, AZ40XT) is applied at a 20 µm thickness to valve sections of the flow layer. For the control layer, SU-8 3025 is spin-coated at 3,800 RPM to generate ~20–25 µm height features. Original graphical patterns (**Supplementary Fig 8**) are designed using AutoCAD (Autodesk, Inc.) and converted with KLayout for standard soft lithography, incorporating 18 distinct input entry points to support multiplex combinatorics ($2^9 = 512$ conditions). All photoresist layers are exposed to a 375 nm laser using a Heidelberg

MLA150 maskless aligner, ensuring precise patterning on a 4-inch silicon wafer (WAPERPRO, Lot number: 21909201). Additional mold preparation details are documented[21-23], and this chip's original design is available in open-source databases (https://github.com/Uchicago-TAY/LIST).

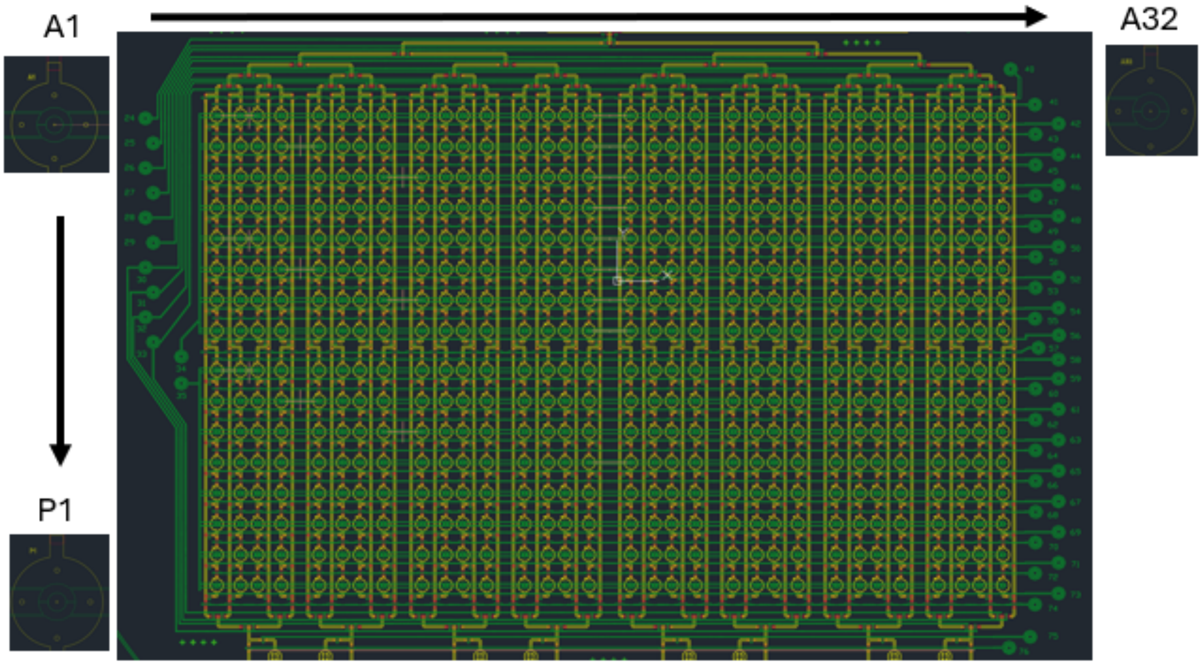


**Supplementary Fig 9.** AutoCAD design: 512 chambers are arranged in a grid, with positional markers labeled A1 to A32 along the top and bottom edges (representing 32 rows) and A1 to P1 along the left and right edges (representing 16 columns).

The chip's fluidic architecture is designed for multiplexing, allowing precise control over reagent delivery to individual chambers. The system includes a combination multiplexer with nine nodes and 18 branched input lines, enabling the delivery of unique reagent combinations (antibiotics at targeted concentrations). An additional input supplies buffer solutions, deionized water, or culture media for washing steps. The row multiplexer, equipped with 10 valves, operates a tree-type flow trafficking system that directs fluid from the inlet nodes to 32 row lines (based on a binary addressing scheme, $2^5 = 32$). The column multiplexer, with 36 valves, controls fluid trafficking and retention to the 512 individual chambers arranged in a 32 x 16 grid. An outlet port facilitates waste purging and hydrogel loading, critical for initializing bacterial cultures and removing excess reagents.

Automation of fluidic control is managed by a multiplexing controller that interfaces with the Excel-based input files. These files map 512 unique reagent combinations to the chamber array, specifying valve actuation sequences and fluid delivery protocols (**Supplementary Fig 9**). The row and column multiplexers work in tandem to direct fluid flow, with pressurized valves isolating chambers during reagent delivery to prevent cross-contamination. The GUI supports both programmable sequential dosing (e.g., delivering reagent A for 3 hours, followed by reagent B) and simultaneous dosing, achieved. The entire experimental protocol involves over 2,500 individual steps, including reagent delivery, washing, and sample recovery, which is facilitated by degrading the hydrogel matrix and purging the chambers. The operational sequence, governed by a sequence file (see the document below), includes valve actuation schedules and fluid delivery protocols. Time-lapse images demonstrate environmental switching, with a filling checkerboard pattern starting with columns P1 to P32, followed by rows O1 to O32, and finally A1 to A32, each filled with distinct reagents or antibiotic combinations. This design supports individual chamber control, enabling the delivery of 512 unique conditions, critical for high-throughput combinatorial experiments.

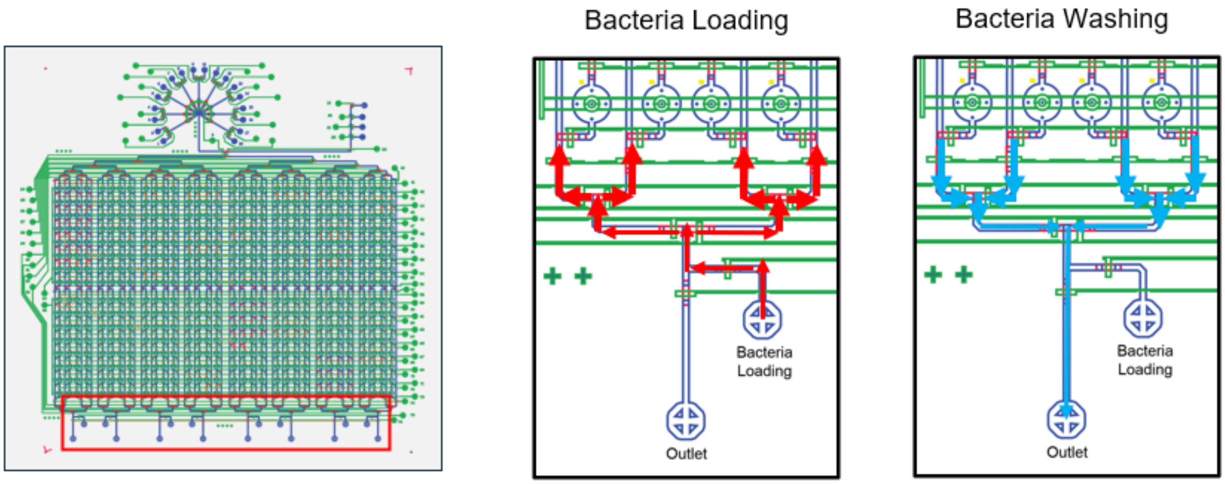


**Supplementary Fig 10.** AutoCAD design of outlet region (red box) for bacteria loading (red arrows) and washing (blue arrows) processes in the chip

**Hydrogel Precursor Instruction through Outlet Port**

To prevent channel clogging and ensure uniform bacterial encapsulation in the chambers, hydrogel precursor (alginate solution, 1-2 % w/v) with bacterial inoculum should be introduced through outlet ports (**Supplementary Fig 10**). Bacteria loading processes (red arrows) indicate the flow direction of the bacterial suspension into the chambers. Washing steps after bacteria loading indicate blue arrows show the flow of washing solution through the system, removing excess bacteria or chemical wastes to prevent cross-contamination. This setup is designed to handle the challenges posed by the viscous alginate solution, ensuring efficient bacterial encapsulation and subsequent processing without clogging, which is critical for consistent experimental outcomes in microfluidic culture studies.

**Reproducibility Test Across Different Spots**

Through the pipelines for colony tracking, we evaluated experimental reproducibility in the chip across different spots (**Supplementary Fig 11a-c,** three chambers, randomly selected) on the same chip and across different chips (**Supplementary Fig 11d**). The growth curves, tracked over a 6-hour period through the pipeline, showed a consistent growth trend across multiple spots within a single microfluidic chip, with local variations likely attributable to environmental factors. These chip consistencies underscore the precision of the microfluidic platform in maintaining uniform growth conditions between chambers. Further, analyses between three chips, supported by detailed box plot representations of normalized average sizes and validated through unpaired t-tests (yielding p-values of 0.412 and 0.968), demonstrate comparable mean values and interquartile ranges, indicating statistically non-significant differences despite minor experimental variability inherent in the fabrication or operational processes of the chips.

In all subsequent experiment, we included a local control group and at least biologically independent replicates (N=3 per condition) further enhance the reliability of our experimental setup, providing a solid foundation for the microfluidic system's performance.

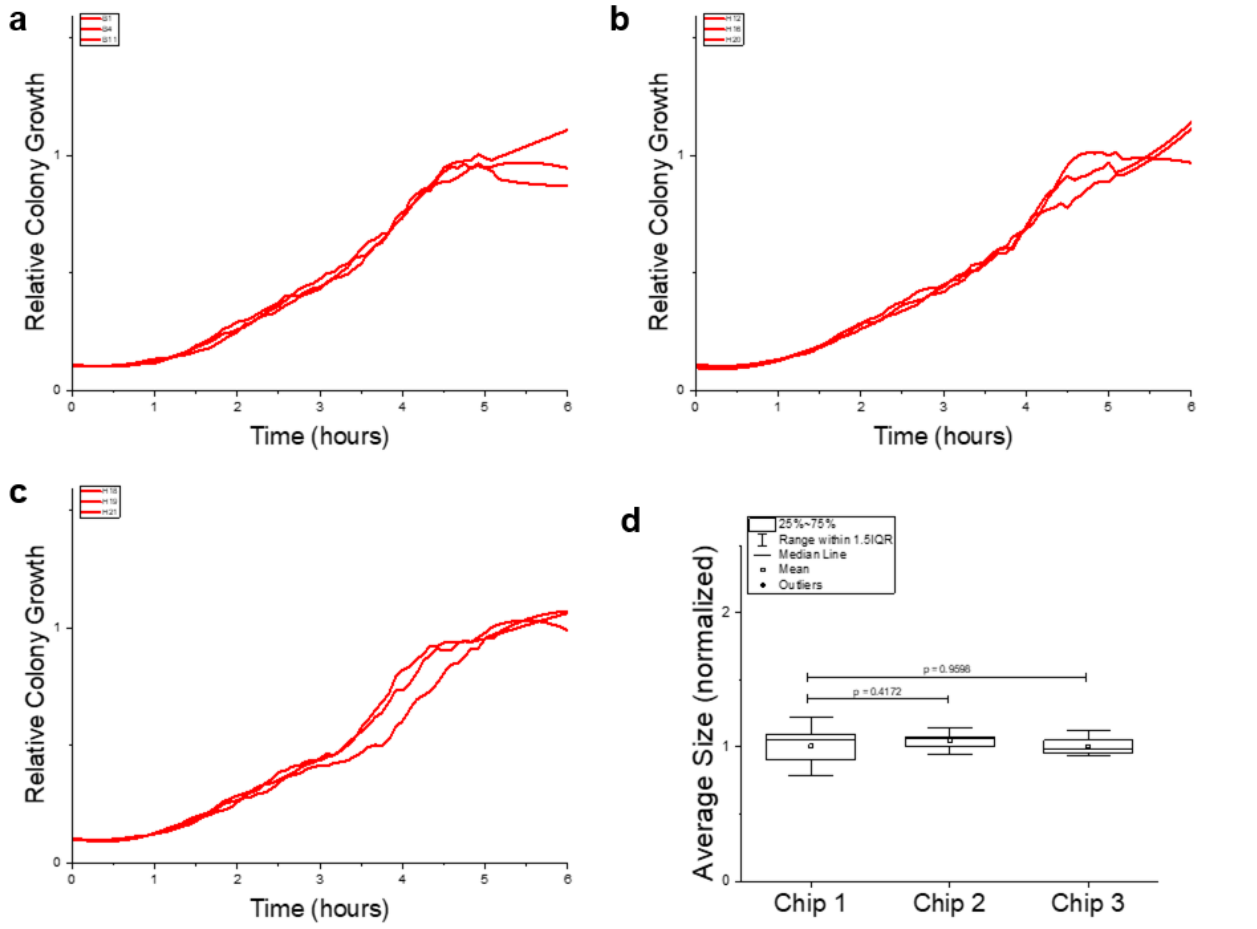


**Supplementary Fig 11**. **Reproducibility Test Across Different Spots in the Same Chip and Across Different Chips** (a-c) Growth curves of *E. coli* relative colony growth over time (0-6 hours) across different spots on the same microfluidic chip, illustrating a consistent growth pattern with minor local variations. The data were normalized on a 0-1 scale using the pipeline’s area measurement and normalization steps, with the mean colony average size represented by a red line (n = 30 colonies per condition). (d) Whisker plot comparison of normalized average sizes across three different microfluidic chips (Chip 1, Chip 2, Chip 3), analyzed via unpaired t-tests (N = 5 per condition, biologically independent replicates). The box plots display the median (line), mean (dot), range within 1.5 IQR (whiskers), and outliers (points), conducted in LB media. The experimental data highlights a robust reproducibility in the growth dynamics of E. coli cultured in LB media across both different spots on the same chip and different chips.

### Supplementary Note 5. Synergy Evaluation and Combination Index

The combined effect of antibiotic pairs was evaluated using a modified Loewe Additivity Model[24-26], defined by two antibiotic A and B:

$$\frac{x_a}{X_a} + \frac{x_b}{X_b} = 1$$

Where,

$x_a$ and $x_b$ : doses of antibiotics A and B in combination

$X_a$ and $X_b$ : doses of antibiotics A and B, when used alone produce the same effect as the combination effect ($f_c(x)$)

This model assumes an additive interaction between two antibiotics, where the doses of the antibiotics in combination produce the same effect as specific doses of each antibiotic alone. The expected effect is modeled using a 4-parameter log-logistic (4PL) function $f(x)$ as a function of dose $x$, characterized by:

$$f(x) = E_{min} + \frac{E_{max} - E_{min}}{1 + (\frac{x}{m})^{-h}}$$

- $E_{min}$ : Minimal effect (no inhibition, full growth = 0 on a 0-1 scale).
- $E_{max}$ : Maximal effect (complete inhibition, ≤ 1 on a 0-1 scale).
- $m$ : Relative EC50/IC50 (concentration for 50% effect).
- $h$ : Shape parameter (steepness of the curve).

Assumptions:

- Both antibiotics contribute to the same effect (growth inhibition), allowing dose equivalence.

- Dose-response curve is monotonic, typically modeled by a sigmoidal function.
- In the platform, $f_c(x)$ the relative growth inhibition (0 = full growth, 1 = complete inhibition)

Given the combination effect $f_c(x)$, $X_a$, $X_b$:

Using the 4PL model, set the effect $f(x) = f_c(x)$.

For antibiotic A,

$$f_c(x) = E_{a,min} + \frac{E_{a,max} - E_{a,min}}{1 + (\frac{X_a}{m_a})^{-h_a}}$$

Rearrange,

$$X_a = m_a \left( \frac{f_c(x) - E_{a,min}}{E_{a,max} - f_c(x)} \right)^{1/h_a}$$

For antibiotic B,

$$f_c(x) = E_{b,min} + \frac{E_{b,max} - E_{b,min}}{1 + (\frac{X_b}{m_b})^{-h_b}}$$

Rearrange,

$$X_b = m_b \left( \frac{f_c(x) - E_{b,min}}{E_{b,max} - f_c(x)} \right)^{1/h_b}$$

Synergy is quantified using the Combination Index (CI), which compares the observed combination effect (relative growth inhibition) to the expected effect under additivity (Fig 6a). The CI is calculated based on the individual antibiotics' doses, their EC50/IC50 values, and the 4PL model parameters.

$$CI = \frac{x_a}{X_a} + \frac{x_b}{X_b}$$

Substitute $X_a$ and $X_b$ from the expression above:

$$CI = \frac{x_a}{m_a\left(\frac{f_c(x) - E_{a,min}}{E_{a,max} - f_c(x)}\right)^{1/h_a}} + \frac{x_b}{m_b\left(\frac{f_c(x) - E_{b,min}}{E_{b,max} - f_c(x)}\right)^{1/h_b}}$$

CI values are interpreted as:

- CI < 1: Synergy
- CI = 1: Additivity
- CI > 1: Antagonism

Since CI values can exceed 1 (e.g., CI = 2 for strong antagonism), normalization is required for visualization and decision-making (CI < 0.35 for synergy, CI > 0.65 for antagonism).[27]

A normalization method is:

$$CI_{normalized} = \frac{1}{1 + e^{k(1-CI)}}$$

Where, $k$ is a scaling factor to map CI values to 0–1.

- CI < 1: Synergy maps to $CI_{normalized} \leq 0.35$
- CI = 1: Additivity maps to $0.35 < CI_{normalized} < 0.65$
- CI > 1: Antagonism maps to $CI_{normalized} \geq 0.65$

Results were visualized in heatmaps (Fig 6b-c), with red indicating synergy, white for additivity, and blue for antagonism, scaled from 0 to 1, noting synergy score shift ($\Delta CI_{normalized}$ > 0.3)

**Modification of the Loewe Model**

Combining the Loewe Additivity Model and CI formular with a biphasic growth curve requires significant modifications to account for the non-standard dose-response behavior of biphasic curves (Fig 2m-o).

To use a biphasic dose-response model, replace the 4PL model with a biphasic dose-response model. A common approach is to model the response as a sum of two logistic functions:

$$f(x) = E_{min} + \frac{w \cdot (E_{a,max} - E_{min})}{1 + (\frac{x}{m^a})^{-h_a}} + \frac{(1 - w) \cdot (E_{b,max} - E_{min})}{1 + (\frac{x}{m^b})^{-h_b}}$$

- $w$ : Weighting factor for the initial phase (0 ≤ $w$ ≤ 1)
- $m_a, m_b$ : EC50/IC50 for the first and second phase.
- $h_a, h_b$ : Shape parameter for each phase.

The Loewe Additivity Model and the CI formula rely on a monotonic dose-response relationship, modeled by the 4PL function[28]. However, biphasic growth curves we measured in the ultra-multiplexed platform challenge this assumption, rendering the standard Loewe model unsuitable without modification. While an alternative such as the Zero Interaction Potency (ZIP) model[29, 30] could address non-monotonic responses, the complexity and lack of analytical tractability of a biphasic dose-response model make it impractical and complicated for high-throughput analysis. To maintain compatibility with the Loewe model's CI framework[25] while ensuring computational feasibility, this study adopts a simplified approach by focusing on the second growth phase following media refresh and antibiotic administration at 3 hours. The 4PL model is applied to describe relative growth as a function of dose during this post-3-hour period, effectively capturing the experimental dynamics while avoiding the challenges of fitting biphasic curves. This strategy reflects the cumulative effects of sequential antibiotic exposure or media adjustments in dosing experiments, providing a robust and practical method for evaluating synergy in a controlled, biofilm-like culture setting.

#### Supplementary Note 6. In-depth Insights of sequential combination therapy

In Fig 6, the SUL-TRI pair showed synergistic effects, consistent with their role in the sequential blockade of folate synthesis[31]: SUL inhibits dihydropteroate synthase (DHPS), TRI inhibits dihydrofolate reductase (DHFR)[32]. Synergistic interactions between AMI and TRI appear to result from parallel inhibition pathway of protein synthesis (30S subunit by AMI) and folate synthesis (DHFR by TRI). Antagonistic interactions in the SUL-TET pair are likely attributable to overlapping bacteriostatic effects[33] on folate synthesis (DHPS by SUL) and protein synthesis (30S ribosomal subunit by TET).

Three pairs (CEF-NIT, CEF-SUL and SUL-RIF) exhibited substantial changes with a score shift >0.5 in sequential (A→B or B→A) dosing studies. The CEF-NIT pair exhibited an additive interaction in simultaneous and NIT→CEF dosing, shifting to synergy in CEF→NIT dosing, likely driven by NIT's concentration-dependent transition from bacteriostatic to bactericidal activity[10]. Our results indicated that bacteriostatic drugs, such as SUL and TET, can inhibit bacterial growth, potentially reducing the effectiveness of bactericidal drugs like AMI and LEV when used sequentially. This is because bactericidal drugs rely on active cell division for killing, and slowed growth induced by bacteriostatic agents can hinder this process. However, it is worth noting that not all combinations show this antagonism, and some can even be synergistic.

We observed that the interaction between SUL and RIF shifted from antagonism in simultaneous dosing to an additive interaction in SUL→RIF sequential dosing (score shift >0.5). RIF's induction of cytochrome P450 enzymes—a crucial class of liver enzymes (e.g., CYP3A4 and CYP2C9) could reduce drug efficacy[34,35,36], particularly due to low medication adherence. In addition, SUL, metabolized by CYP2C9, could decrease in plasma when co-administered with RIF, which compromises the efficacy of SUL-RIF combinations in treating CA-MRSA infections[37]. CLI, metabolized by CYP3A4, is known as to be significantly affected by RIF's induction, which consequently lowers CLI concentrations and risk treatment failure, especially in infectious diseases requiring sustained drug levels[37]. In contrast, TRI could be primarily excreted as unchanged in urine, making this type of drugs less susceptible to RIF's effects[34]. Similarly, aminoglycoside class drugs (e.g. AMI in this study), primarily cleared renally with minimal CYP metabolism, remains unaffected by RIF's induction[36], making it also a stable co-administered agent in such in vivo regimens.

In clinical studies, the combination of SUL and TRI has achieved high cure rates[37 38], comparable to monotherapy in community-acquired methicillin-resistant *Staphylococcus aureus* (CA-MRSA) skin and soft tissue infections[37]. These types of infections can be complicated by biofilms, which promote persister-mediated resistance in multidrug-resistant (MDR) pathogens. We demonstrated time-dependent biphasic growth patterns which could be characterized by initial bacterial killing and certainly associated with slower clearance of persister cells[15]. This offers a robust dynamic model for chronic infections to emulate real world in vivo drug regimens[39 40 41 42]. Sequential dosing of CLI by targeting the 50S ribosomal subunit, followed by RIF inhibiting RNA polymerase[35], reduced bacterial load and targeted persister cells, mitigating resistance[43], consistent with our synergy observation (Fig 6f) in CLI→RIF dosing (score shift >0.3). Similarly, synergy in the CEF→NIT dosing (score shift >0.5) likely depends on achieving a therapeutic NIT concentration, which might be sufficient to shift from bacteriostatic to bactericidal activity[10]. For E. coli urinary tract infections, where NIT is commonly used[44], combining CEF with high-dose NIT in a CEF→NIT sequence could enhance efficacy against MDR strains. Further, sequential administration of LEV, targeting DNA gyrase, followed by CEF, inhibiting cell wall synthesis, enhanced efficacy against biofilm-related resistance[45] in a LEV→CEF sequence (score shift >0.3). Studies incorporating biofilm-disrupting agents (e.g., DNase[46]) or quorum-sensing inhibitors (e.g., hamamelitannin[47]) could further improve outcomes. Many clinical studies for combination therapies align with in vitro research, but efficacy varies by bacterial strain and infection site.

Finally, we rely on the Loewe Additivity model[25 48], for evaluating antibiotic pairs with consideration of similar mechanism of action (e.g., AMI-TET targeting the 30S ribosomal subunit). This model for synergy decision could not fully capture complex drug interactions for antibiotics with independent mechanisms, where the Bliss Independence or other advanced synergism models (e.g., highest single agent [HSA], zero interaction potency [ZIP]) could be more desirable for complementary insights[24 49] (as discussed in Supplementary Note 5). However, it still lacks the experimental depth of our 3D culturomics platform, which captures response shifts in sequential dosing (e.g., SUL-TRI, SUL-RIF), highlighting the importance of experimental models like ours. These ground works need to be a robust theoretical framework for predicting pairwise interactions, supported by references.

**Supplementary Data. GUI control for antibiotic testing ("Multiplexing control.xlsx")**

The " Multiplexing control.xlsx" file (Extended Data Fig 4a, (7)) supports GUI control for antibiotic testing. (A) An excerpt of the Excel file displays the first 10 rows, detailing 512 unique experimental conditions for testing combinations of nine antibiotics (amikacin, cefepime, clindamycin, levofloxacin, nitrofurantoin, rifampicin, sulfamethoxazole, tetracycline, and trimethoprim). The "Input Combinatorial input" column specifies active antibiotics (e.g., "1" for amikacin, "1,2" for amikacin + cefepime), with columns A–I listing corresponding valve states (e.g., [2, 3, 5, 7, 9, 11, 13, 15, 17] for Condition 1). (B) A diagram illustrates the full factorial design, covering individual and pairwise antibiotic interactions. (C) A workflow schematic shows the integration of the Excel file with a custom GUI, mapping valve states to 18 valve controls (multiplexed across 80 total valves) for precise delivery of antibiotic combinations to the 512 chambers. (D) A schematic of the microfluidic chip's layout illustrates how the 512 chambers receive unique antibiotic combinations via 18 valve controls, with fluid routing indicated by color-coded channels. (E) A flowchart details the GUI's role in importing the Excel file and translating sequences into real-time valve operations, ensuring accurate antibiotic delivery to each chamber.

(Also see the additional file "Multiplexing control.xlsx")
